# Functional architecture of cardiac TF regulatory landscapes in control of mammalian heart development

**DOI:** 10.64898/2025.12.19.695499

**Authors:** Virginia Roland, Johannes Tüchler, Andrea Esposito, Mattia Conte, Matteo Zoia, Ekapaksi Wisnumurti, Virginie Tissières, Julie Gamart, Raquel Rouco Garcia, Ines J. Marques, Akshay Akshay, Vincent Rapp, Brandon J. Mannion, Jennifer A. Akiyama, Prateek Arora, Harry Walker, Ali Hashemi Gheinani, Beth A. Firulli, Gretel Nusspaumer, Anthony B. Firulli, Guillaume Andrey, Axel Visel, Nadia Mercader, Javier Lopez-Rios, Mario Nicodemi, Iros Barozzi, Marco Osterwalder

**Affiliations:** Department for BioMedical Research (DBMR), University of Bern, Bern, Switzerland; Center for Cancer Research, Medical University of Vienna, Vienna, Austria; Department of Physics, University of Naples Federico II and National Institute of Nuclear Physics, Naples, Italy; Department of Cardiology, Bern University Hospital, Bern, Switzerland; Department of Genetic Medicine and Development and iGE3, Faculty of Medicine, University of Geneva, Geneva, Switzerland; Institute of Anatomy, University of Bern, Bern, Switzerland; Environmental Genomics and Systems Biology Division, Lawrence Berkeley National Laboratory, Berkeley, CA 94720, USA; Herman B Wells Center for Pediatric Research, Departments of Pediatrics, Anatomy and Medical and Molecular Genetics, Indiana Medical School, Indianapolis, IN 46202, USA; Centro Andaluz de Biología del Desarrollo (CABD), CSIC-Universidad Pablo de Olavide Junta de Andalucía, Seville, Spain; US Department of Energy Joint Genome Institute, Lawrence Berkeley National Laboratory, Berkeley, CA 94720, USA; School of Natural Sciences, University of California, Merced, CA 95343, USA; Centro Nacional de Investigaciones Cardiovasculares Carlos III, Madrid, Spain; Universidad Loyola Andalucía, School of Health Sciences, Seville Campus, 41704, Dos Hermanas, Seville, Spain

## Abstract

Congenital heart disease (CHD), the most common human birth defect, often results from disruptions in gene regulatory networks (GRNs) that control cardiac lineage specification and cell type identity during heart development. A conserved core set of cardiac transcription factors (TFs) orchestrates these processes through combinatorial interactions that are cell type-specific and tightly regulated across space and time. However, the genomic enhancer architecture that integrates upstream effectors to establish precise cardiac TF dosage and downstream transcriptional output remains largely unresolved. Here, we assessed the functional necessity of five developmental heart enhancer modules previously linked to the regulation of *Gata4* and *Hand2*, core cardiac TFs exhibiting overlapping roles in myocardial and endocardial development. While individual enhancer deletions in mouse embryonic hearts revealed a surprising degree of transcriptional resilience, a subset of *Gata4* enhancers proved indispensable for embryonic progression in a genetically compromised background. To achieve higher precision in cardiac cell type-specific enhancer prediction, we applied single-nucleus multiome profiling, enabling the delineation of cardiac cistromes underlying heart morphogenesis. By integrating this resource with deep learning applications, site-directed transgenesis, and chromatin conformation modeling, we mapped the cardiac enhancer repertoire and regulatory signatures that orchestrate *Hand2* dynamics across distinct cardiac compartments and lineages. Genome editing further revealed an essential role for the *Hand2*-upstream regulatory interval (*H2*-URI) in transcriptional control of endocardial lineage effectors and, consequently, trabecular network formation and cardiac cushion patterning. Together our findings highlight substantial resilience in the *cis*-regulatory architectures governing cardiac TF dynamics and demonstrate that combinatorial integration of upstream lineage identities across modular enhancer landscapes establishes the cardiac cell type-specific programs driving heart morphogenesis. These results advance the reconstruction of cardiac GRNs and enhance the functional interpretation of CHD-associated variants.

## Introduction

Congenital heart disease (CHD), arising from disruptions in four-chambered heart morphogenesis, is the most common human structural birth anomaly^1^. It encompasses a broad spectrum of malformations, from isolated defects to complex syndromic abnormalities^2^. CHD also contributes substantially to adverse pregnancy outcomes, with affected fetuses experiencing an estimated 1–13.5% risk of pregnancy loss^3^. Genetic factors are major contributors to CHD risk and comprise chromatin perturbations, copy-number variation, and coding mutations that disrupt transcriptional control, signaling pathways, or cardiomyocyte (CM) function^1,4^. CHDs frequently arise from coding mutations in cardiac transcription factors (TFs), which orchestrate gene regulatory networks (GRNs) governing lineage commitment and spatial patterning of cardiac progenitors^5,6^. Acting together with signaling pathways and cofactors, these TFs maintain an evolutionarily conserved regulatory synergy detectable from bilaterians to vertebrates^5^. A core “cardiac regulatory kernel” including TFs from different families such as *Nkx2-5*, *Gata4*, *Mef2c*, *Tbx5*, and *Hand2* coordinates cardiac progenitor specification and differentiation across both first and second heart fields (FHF, SHF), sculpting the four-chambered heart from an evolutionary conserved template^7–9^. This regulatory program, operating through multi-scale DNA-protein and protein-protein interactions, underlies the myocardial contractile and conductive identities of CMs essential for cardiac function^10–13^. Such GRNs are likewise interconnected with the regulatory framework governing endocardial (EC) development, which encompasses trabecular network formation driven by endocardial–myocardial crosstalk, as well as the generation of cushion mesenchymal cells, a substantial proportion of which arise from endocardial-to-mesenchymal transition (EndMT) and subsequently remodel into valve structures^14,15^. While FHF progenitors form the primitive heart tube and largely contribute to left ventricular (LV) myocardium, SHF cells generate most of the right ventricle (RV), outflow tract (OFT), and parts of the atria^9,16,17^. Heart development relies on additional progenitor populations outside the early heart tube, including cardiac neural crest cells (cNCCs) and proepicardial/epicardial (EP) cells, which contribute to OFT and great vessel smooth muscle (cNCCs), as well as coronary vascular smooth muscle, cardiac fibroblasts, and parts of the coronary vasculature and cushion/valve mesenchyme^9,18,19^. All these cardiac lineages and cell types are dependent on instructive combinatorial and tightly controlled expression dynamics of cardiac TFs^13,20^. Complete loss of expression of most core cardiac TFs leads to embryonic lethality, primarily due to severe cardiac defects such as looping abnormalities or chamber malformations^21–25^. A subset of these genes, including *Tbx1* and *Tbx5*, even exhibit haploinsufficiency that leads to syndromic CHD^24,26^. Compartment- or cell type-specific deficiency of core cardiac TFs expressed across multiple cardiac lineages, such as *Gata4* or *Hand2*, often leads to cell type-restricted or isolated CHD-related defects, including myocardial thinning, chamber hypoplasia, OFT and septal defects, trabeculation defects or cardiac cushion abnormalities^27–30^. Such abnormalities are typically linked to defects in OFT/RV specification or differentiation, EC function or OFT remodeling^18,31,32^, emerging from disrupted cardiac cell type GRNs wired through genomic enhancers^10,33,34^. Together, these findings underscore that precise spatiotemporal regulation of core cardiac TF dosage, tailored to specific compartments or cell types through enhancers, is essential for normal heart development and for preventing cardiac abnormalities^35–37^.

Although protein-coding variants in cardiac TFs and other developmental genes contribute to CHD^2^, they explain only a limited proportion of the genetic risk involved, with more than half of familial and sporadic cases still lacking a molecular diagnosis^1,38^. Remarkably, most CHD-associated *de novo* mutations and common risk variants map to noncoding regions, where a substantial fraction is thought to disrupt TF motifs and developmental enhancer function^37,39–41^. However, due to the limited availability of catalogs comprehensively mapping CHD-associated single nucleotide polymorphisms (SNPs) and the lack of large-scale validation workflows, only a comparatively small number of heart enhancer-localized variants have been functionally validated *in vivo*^33,42–44^. Structural variants, such as large deletions that disrupt enhancer clusters or topologically associated domain (TAD) boundary elements, can also exert pathogenic effects by altering genome topology and causing aberrant enhancer-gene interactions^45–47^.

*In vivo* heart enhancers are typically identified by the convergence of H3K27ac and H3K4me1 histone marks, open chromatin architecture, and binding signatures of lineage-defining TFs^13,37,48^. Predicting and functionally resolving cardiac enhancer activity at cell type resolution remains a major challenge, as this requires precise correlation with target gene activity and *in vivo* perturbation strategies^49–51^. The difficulty is particularly pronounced at TF loci, which often coordinate diverse cardiac and extracardiac regulatory programs and are embedded within large, gene-poor TADs^33,52,53^. Functional studies of enhancers near structural or ion channel genes have uncovered diverse, and sometimes context-dependent, roles of cardiac enhancers in heart development and physiology^40,54,55^. By contrast, few TF-associated enhancers have been systematically characterized in their native genomic setting in terms of functional necessity, and those analyzed displayed remarkable variability in functional importance^33,53,56,57^. As a notable example, a proximal *Nkx2-5* enhancer is critically required for regulation of RV conduction and for conferring protection against neonatal lethality arising from cyanotic conotruncal malformations^57^. Consequently, elucidating the precise roles of individual cardiac TF enhancers is critical for functional interpretation of CHD variants, yet insights into individual enhancer mechanisms are only beginning to emerge^33,37,58^.

In this study, we explored the functional enhancer architecture underlying mammalian heart morphogenesis, with a particular emphasis on the regulatory landscapes controlling the cardiac core regulators *Gata4* and *Hand2*. These TFs exhibit overlapping roles within the SHF-associated GRNs and are indispensable for the proper development and coordination of myocardial and endocardial populations^25,27–30^. In addition, *Hand2*, through its function in cardiac neural crest cells (cNCCs), regulates the development and alignment of the OFT and aortic arch arteries^14,59^. Here, through systematic endogenous deletion of previously identified *Gata4*- and *Hand2*-associated heart enhancer elements^60–63^ in mouse embryos, we observed remarkably widespread transcriptional resilience, consistent with pervasive enhancer redundancy in the genomic landscapes of developmental regulator genes^52,64,65^. However, enhancer loss in the context of reduced target gene dosage caused embryonic lethality associated with respective cardiac TF functions, implicating these sequences as conditional disease modifiers. To achieve a comprehensive delineation of cardiac enhancer landscapes at genome-scale and cell type resolution, we performed single-nucleus multiome profiling of developing mouse hearts during cardiac chamber expansion stages. This integrative approach enabled high-resolution reconstruction of cardiac TF cistromes and facilitated systematic prediction of enhancer activity across distinct cardiac cell types and subpopulations. Using this multiomic framework, we characterized a new repertoire of *in vivo Hand2* heart enhancers implicated in mediating essential compartment- and cell type-specific activities. Combining chromatin conformation polymer modeling and genome editing, we uncover an essential function of the non-coding *Hand2* upstream regulatory interval (*H2*-URI) as a gatekeeper domain that controls cardiac progression and key endocardial lineage functions, including trabecular network morphogenesis and cardiac cushion patterning. Collectively, our findings establish transcriptional robustness as a predominant principle of cardiac TF enhancer landscapes, delineate the enhancer architecture governing dose-dependent, compartment- and cell type–specific TF activity, and underscore the fundamental role of these regulatory landscapes in cardiogenesis and the mechanisms underlying CHD.

## Results

### Resilience of cardiac TF regulatory landscapes to individual heart enhancer loss

To elucidate how the cardiac compartment and cell type-specific expression dynamics of core cardiac regulators are orchestrated, we examined the loci of the *Gata4* and *Hand2* TFs, which are known to integrate several lineage-specific functions, while occupying an epistatic hierarchy that governs morphogenesis of the SHF-derived cardiac compartments such as the OFT and RV^27,60,66,67^. Using transgenic reporter assays in mouse embryos, previous studies have identified several *Gata4* and *Hand2*-associated cardiac enhancers located within the respective topologically associated domains (TADs), yet their individual functional contributions in native context remained elusive^60–63,68^ (**Figs. 1A; S1A, B**). For *Gata4*, this included distant enhancer elements active throughout the heart (mm245, *G4*-124)^61^, within the outflow tract (OFT) and ventricles (mm244, *G4*+182)^61^, or restricted to the endocardium and EC cushions (G9, *G4*-93)^62^ (**Figs. 1A; S1A, C**). To enable a more systematic comparison of enhancer activities during midgestation, we performed site-directed reporter transgenesis (enSERT)^69^ in mouse embryos at E10.5 and E11.5. This analysis revealed that *G4*-124 exhibited predominantly myocardial activity, whereas *G4*+182 drove expression restricted to the ventricular endocardium and the OFT endothelium, similar to *G4*-93 (**Fig. 1B, S1C**). Analysis of accessible chromatin and H3K27ac profiles from mouse embryonic hearts^53,70^ further identified an additional element with cardiac enhancer signatures (*G4*+177) directly adjacent to *G4*+182 (**Fig. 1C**), which drove strong subregional myocardial and endocardial activities restricted to the OFT and parts of the ventricles, as well as within the EC cushions (**Figs. 1B; S1A, C, Table S1**). Section analysis at E10.5 further localized additional *G4*-93 and *G4*+177 activities in the sinus venosus (inflow tract) region known to contain sinoatrial node (SAN) progenitors^43^ (**Fig. S1D**). Unlike *G4*-93, *G4*+177 also exhibited activity in the OFT and AV canal (AVC) myocardium (**Fig. S1D**). For *Hand2*, despite its more pronounced SHF lineage specificity, the number of previously linked murine heart enhancers was restricted to two TSS-proximal elements driving activity in the RV/OFT (*H2*-3)^60^ or exclusively in the OFT cushions (*H2*+7)^63^, and one far-upstream element exhibiting OFT/RV activity (*H2*+437)^68^ (**Fig. 1A, S1B**). Collectively, the enhancer activities associated with *Gata4* and *Hand2* expression dynamics observed in the OFT/RV and EC cushion regions at midgestation corresponded to areas of strong GATA4/HAND2 overlap (**Fig. S1E**), suggesting involvement in shared cardiac cell type GRNs. While GATA4 was detected throughout the myocardium, HAND2 was restricted predominantly to SHF-derived populations, including the OFT, RV and cardiopharyngeal (cp) mesoderm, in accordance with previous expression-based studies^27,28,71^ (**Fig. S1E**). Co-localization of high levels of GATA4 and HAND2 was evident in the ventricular endocardium, as well as within the AV cushion endocardium and mesenchyme (**Fig. S1E**), consistent with their overlapping and essential roles in trabecular development and cushion morphogenesis^29,30,51,72^.

**Figure 1:**
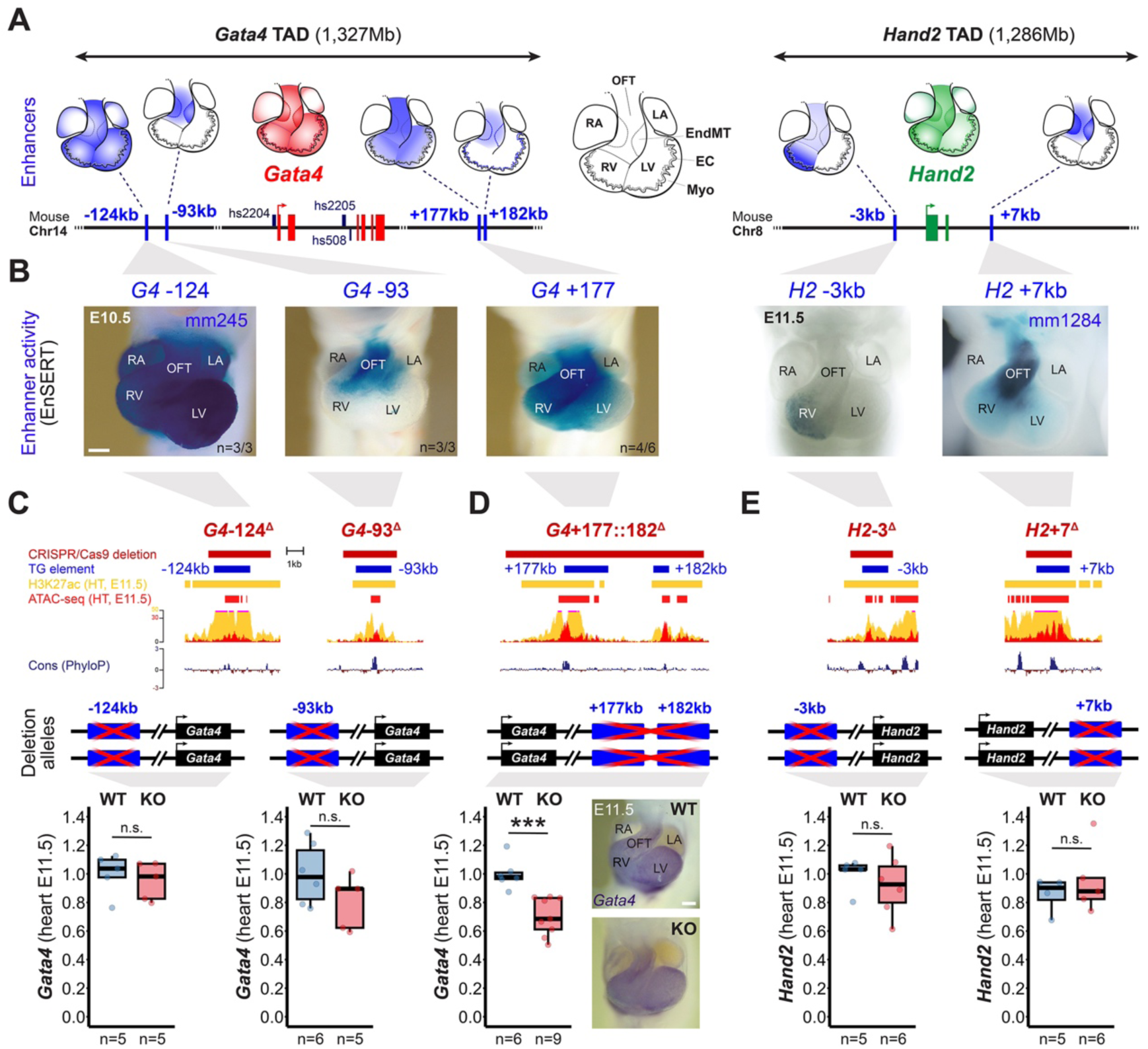
Resilience of cardiac transcription factor (TF) dosage to individual heart enhancer loss. (**A, B**) Location of previously identified mouse heart enhancers within *Gata4* (*G4*) and *Hand2* (*H2*) TADs (**Fig. S1A, B**). Enhancers are designated based on their distance (in kb) either upstream (-) or downstream (+) from the respective transcriptional start site (TSS): *G4*-124 (mm245)^61^, *G4*-93 (G9)^62^, *G4*+177 (this study), *G4*+182 (mm244)^61^, *H2*-3 (*Hand2* -4.2 to -2.7 element)^60^ and *H2*+7 (mm1284)^63^. (**B**) Left: Comparative analysis of Gata4 enhancer activities at E10.5 using site-directed reporter transgenesis (enSERT)^69^ (see also **Fig. S1A-D**). Right: Representative LacZ reporter activities detected for previously described *Hand2* enhancers. (**C-E**) Generation of CRISPR-engineered *Gata4* and *Hand2* enhancer deletion alleles in mice and evaluation of gene dosage effects during heart development. Top panels: targeted genomic enhancer regions with the deleted region (dark red bars), elements validated in transgenic (TG) reporter assays (blue bars), H3K27ac enrichment (ENCODE) (yellow) and accessible chromatin profile (red)^53^ shown (**Table S2**). Cons., placental mammal conservation by PhyloP. Middle panels: allele configurations analyzed. Bottom panels: Quantitative qPCR analysis of *Gata4* or *Hand2* mRNA in homozygous enhancer knockout (KO) compared to WT control embryonic hearts at E11.5. Box plots indicate interquartile range, median, maximum/minimum values (bars) and individual biological replicates (n). P-values are shown, with ***P < 0.001 (two-tailed, unpaired t-test). N.s., not significant. For the *H2*+7 qPCR, one WT and one KO outlier datapoint are outside of the scale shown. Spatial distribution of *Gata4* detected by ISH in E11.5 mouse embryonic hearts carrying the *G4*+177::182 enhancer deletion (D). “n” indicates number of biological replicates analyzed, with similar results. Scale bars, 200μm. OFT, outflow tract. RA, right atrium. LA, left atrium. RV, right ventricle. LV, left ventricle. Myo, myocardial cells. EC, endocardial cells. EndMT, endocardial-to-mesenchymal transition/transformed cells of the cushions.

To evaluate the functional necessity of individual *Gata4* and *Hand2* heart enhancers, we employed CRISPR-EZ^73^ to generate five enhancer knockout (KO) mouse models by individually targeting the three distally located *Gata4*-associated regulatory modules (*G4*-124^Δ^, *G4*+177/182^Δ^, *G4*-93^Δ^) as well as the two proximal *Hand2* enhancers (*H2*-3^Δ^, *H2*+7^Δ^) (**Figs. 1C-E; S2A, B; Tables S2, S3**). In line with the reported weaker direct sequence conservation of heart enhancers relative to enhancers active in other organ tissues^61^, all of these elements showed conserved cores largely restricted to mammals and marked by accessible chromatin flanked by H3K27ac in hearts at E11.5 (**Figs. 1C-E; S2A**). Reprocessing of available cardiac TF ChIP-seq datasets^72,74,75^ further revealed GATA4 and TBX5 occupancy at nearly all enhancer cores, whereas HAND2 was detected at only a subset, suggesting the presence hierarchical or mutual feedback interactions within GRNs (**Fig. S2A**). Across all enhancer KO mouse lines generated, both founder and F1 animals heterozygous for the enhancer deletions were viable and fertile. Remarkably, embryos and adult mice homozygous for each of the enhancer KO alleles analyzed were likewise viable and fertile, and showed no overt phenotypic constraint (**Figs. 1C-E, S2C**). Neither qualitative nor quantitative analyses of *Gata4* or *Hand2* in the heart showed significant changes upon deletion of individual enhancers (**Fig. 1C-E, S2C; Table S4**). Notably, only embryos homozygous for the deletion encompassing both the *G4*+177 and *G4*+182 enhancer modules displayed an approximately 30% *Gata4* reduction in embryonic hearts at E11.5, while spatial expression remained normal (**Fig. 1D**). Together, our findings indicate that distinct compartment- or cell type-specific levels of cardiac TFs are often maintained by a robust regulatory architecture that buffers against the harmful effects of individual heart enhancer deletions. This pattern is consistent with the widespread additive or redundant enhancer relationships associated with critical TF functions in the development of other organ systems, such as the limb^52,65,76–78^.

To explore whether such cardiac enhancer sequences functionally integrate upstream factors across evolutionarily distant GRNs^79^, we examined whether the *Gata4* enhancers retained context-specific cardiac activity in zebrafish embryos despite the lack of genomic conservation (**Figs. S2A, S3**). To this end, we carried out *Tol2*-mediated fluorescent mCerulean enhancer-reporter transgenesis in zebrafish embryos^80^ (**Fig. S3**). Remarkably, screening in transgenic zebrafish founders indicated that each mouse heart enhancer element intrinsically was able to drive transcriptional activity in the zebrafish heart (**Figs. S3A, B; Table S5**). To examine conserved cell type and regional specificity, we generated transgenic F1 zebrafish lines for two enhancers (*G4*-93, *G4*+182) displaying overlapping activity in the mouse OFT endothelium (**Figs. S1C, S3**) and crossed them with *myl7:mRFP* and *fli1:dsRed* reporter alleles to assess respective cardiac cell type activities at 3-4 days post-fertilization. While *G4*-93 failed to drive reporter activity in zebrafish ECs, we observed enhancer activity instead in ventricular CMs and OFT smooth muscle cells of the zebrafish heart (**Fig. S3C, D**). This activity in CMs may reflect a temporal extension of the early myocardial activity driven by the conserved *G4*-93 core element (G9) in the mouse heart at E8.5^62^. In contrast, the EC specificity observed for *G4*+182 in the OFT and ventricles of mouse hearts at E11.5 appeared conserved in zebrafish embryonic hearts (**Figs. S1C; S3E, F**). Therefore, in line with the concept of conserved modular enhancer grammar across evolutionarily divergent GRNs^79,81^, our findings support that mammalian heart enhancers, despite lacking direct genomic sequence conservation, retain the intrinsic capacity to drive cardiac expression in teleosts, albeit with variation in cell-type specificity.

### Essentiality of normally dispensable cardiac TF enhancers in compromised genetic background

To reconcile the strong mammalian conservation of the core sequences within the targeted enhancers (**Fig. S2A**) with the lack of phenotypic effects following individual enhancer deletions (**Fig. 1**), we next investigated whether enhancer function becomes more critical under conditions of reduced target gene dosage. To this purpose, we leveraged a *Gata4^H2B-GFP^* allele representing a *Gata4* null condition^82^ (**Fig. S4A**) and a *Hand2* null allele (*Hand2*^Δ^) generated in isogenic (FVB) background for accurate comparison with enhancer deletion alleles (**Fig. S4B**). As previously reported, *Hand2* heterozygosity did not visibly affect embryogenesis^25,83^, while mice heterozygous for *Gata4* did not present compromising cardiac abnormalities and were viable and fertile^82^. To investigate how the loss of each previously characterized *Gata4* or *Hand2* cardiac enhancer element influences gene function under genetically sensitized conditions, we intercrossed mice heterozygous for a specific enhancer deletion with mice heterozygous for the corresponding target gene deletion (**Fig. 2A**). This framework enabled comparative analysis between embryos carrying both enhancer and gene deletions in trans (sensitized enhancer KO, sKO) and control littermates that were heterozygous for the target gene deletion alone (**Fig. 2A**). Remarkably, transcript analysis of mouse embryonic hearts at E11.5 revealed notable dose reductions in three of the five sensitized enhancer deletions examined (**Fig. 2B**). As expected, relative to wild-type (WT) alleles, the heterozygous gene loss-of-function (LOF) alleles exhibited approximately 50% reduction in transcript levels. Target gene expression was further decreased by about 30%, 20%, and 10% in the *G4*-93, *G4*+177::182, and *H2*-3kb sKO configuration, respectively (**Fig. 2B**). *G4*-93 sKO hearts further revealed reduced superior and inferior AV cushion leaflets, as opposed to *Gata4* heterozygous (*Gata4*^H2B-GFP/+^) controls and *G4*-124 or *G4*+177::182 sKO genotypes (**Figs. 2C, S4C, D**). This observation is concordant with the predominant EC cushion activity of the *G4*-93 enhancer (**Fig. 1B, S1D**)^62^ and aligns with results from *Tie2*-Cre-mediated conditional inactivation of *Gata4* in ECs, known to entail EMT failure leading to hypocellular cushions^29^. Furthermore, OFT leaflets in *G4*-93 sKO embryos appeared unaffected, in line with the reported absence of OFT cushion defects upon conditional *Gata4* LOF in neural crest-derived cells (NCC) populating mid and distal OFT cushions^29^ (**Fig. 2C**). Most notably, deletion of either the *G4*-93 or the *G4*+177::182 enhancer module in sensitized backgrounds resulted in embryonic lethality with full penetrance at E13.5 or E12.5, respectively (**Fig. 2D, E**). These findings were consistent with the timing of embryonic lethality observed in conditional *Gata4* inactivation models targeting either myocardial or endocardial cells^27,29^. Myocardial *Gata4* LOF is known to cause RV hypoplasia due to reduced CM proliferation at E12.5^27^, while *Tie2*-Cre–mediated *Gata4* deletion in ECs leads to embryonic death at a similar stage^29^. In contrast, no apparent morphological or molecular alterations were observed in *G4*-124, *H2*-3 or *H2*+7 sKO hearts at E12.5 in areas of respective enhancer activities (**Fig. S4C, E**), nor did sKO littermates exhibit overt cardiac phenotypes or embryonic lethality. Together, these results demonstrate that certain cardiac TF enhancers, although seemingly dispensable under normal conditions, possess distinct functional capacities that become essential for heart morphogenesis and embryonic progression when genetic robustness is compromised.

**Figure 2:**
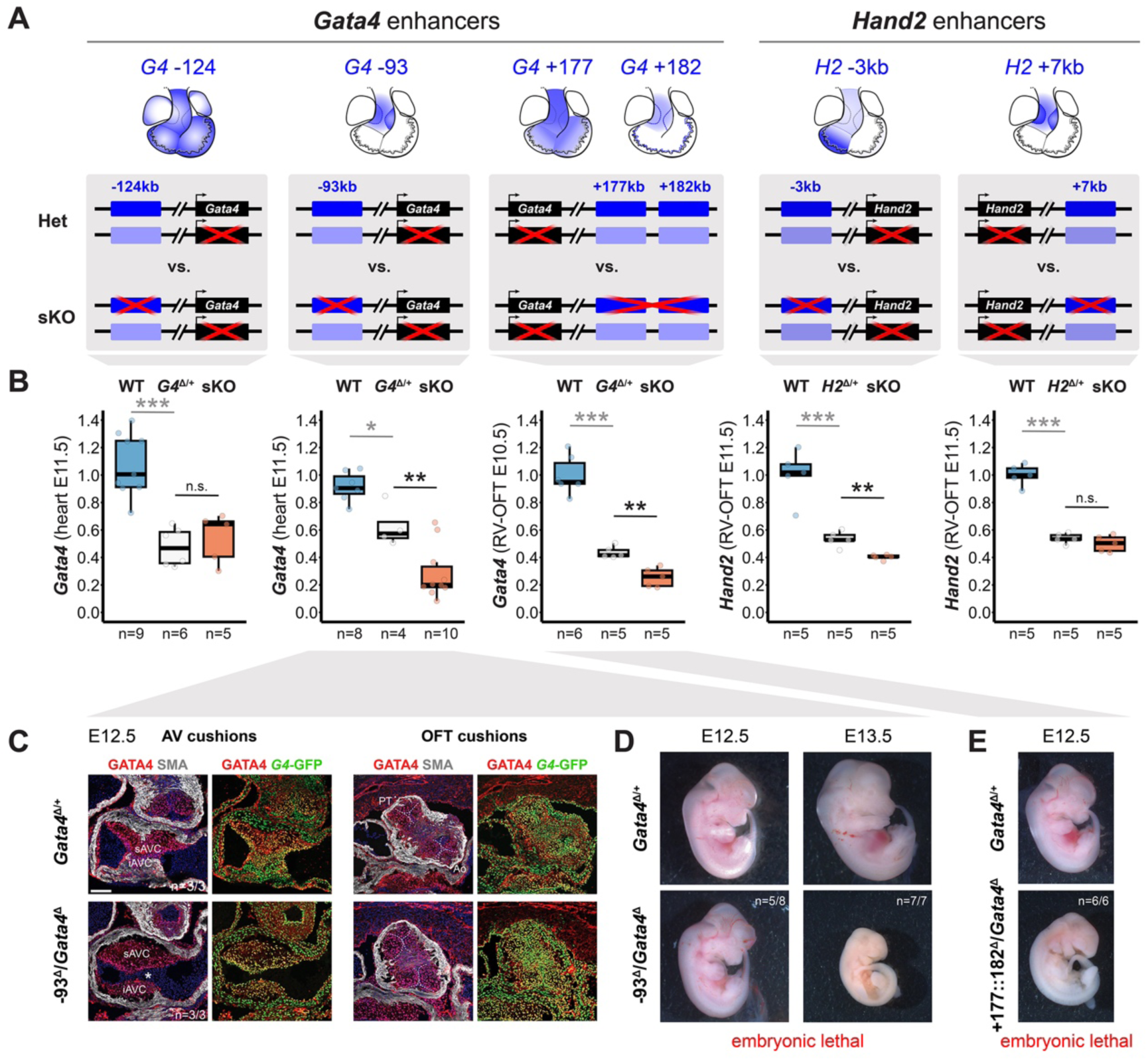
A subset of normally dispensable cardiac TF enhancers is required for heart development and embryonic progression. (**A**) Top: Schematics of *Gata4* (*G4*) and *Hand2* (*H2*) heart enhancer activities (blue) as shown in Fig. 1. Lower panels: Allelic schematics showing deleted target gene and enhancer regions (red crosses) for heterozygous (*Gata4*^Δ(H2B-GFP)/+^ or *Hand2*^Δ/+^) (Het) and sensitized enhancer knockout (sKO) comparisons. Enhancers without functional impact due to target gene deletion in cis are marked light blue. (**B**) Target gene transcript levels (qPCR) from wildtype (WT), Het and sKO mouse embryonic hearts. Box plots indicate interquartile range, median, maximum/minimum values (bars) and individual biological replicates (n). P-values are shown, with ***P < 0.001, **P < 0.01 and *P < 0.05 (two-tailed, unpaired t-test). N.s., not significant. For the *G4*-93 qPCR, one WT outlier datapoint is outside of the scale shown. (**C**) Co-localization of GATA4 protein (red), *Gata4* transcriptional activity driven by the *Gata4*^Δ(H2B-GFP)^ allele^82^ (green) and smooth muscle actin (SMA, gray) in hearts of *G4*-93^Δ^/*Gata4*^Δ(H2B-GFP)^ and *Gata4*^Δ/+^ control embryos at E12.5. White asterisk denotes hypoplastic AV cushions, which are characteristic of decreased *Gata4* levels in endocardial cells^29^. Scale bar, 100μm. (**D, E**) Embryonic lethality observed for *G4*-93^Δ^/*Gata4*^Δ(H2B-GFP)^ (-93^Δ^/*Gata4*^Δ^) and *G4*+177::182^Δ^/*Gata4*^Δ(H2B-GFP)^ (+177::182^Δ^/Gata4^Δ^) genotypes at E13.5 and E12.5, respectively, compared to controls. No embryonic lethality was observed for *G4*-124^Δ^/*Gata4*^Δ(H2B-GFP)^, *H2*-3^Δ^/*Hand2*^Δ^ and *H2*+7^Δ^/*Hand2*^Δ^ sKO genotypes. “n” denotes the number of biological replicates examined, with similar results. E, embryonic day. RV, right ventricle. OFT, outflow tract. s/iAVC, superior/inferior atrioventricular cushion. PT, pulmonary trunk. Ao, Aorta.

### Multiome profiling uncovers regulatory landscapes and TF cistromes across cardiac cell types

The substantial transcriptional resilience observed after homozygous deletion of individual cardiac TF enhancers, even under conditions of sensitized target gene expression (**Figs. 1, 2**), indicated the presence of additional, potentially redundant heart enhancers within the *Gata4* and *Hand2* regulatory landscapes. To achieve more accurate genome-wide prediction of target gene-correlated heart enhancer activities across individual cardiac cell types during chamber expansion stages, we performed simultaneous mapping of chromatin accessibility and transcriptomes in single nuclei (snMultiome, 10x Genomics) of mouse embryonic hearts from E9.5 to E11.5 (**Fig. 3A**). Following stringent filtering (see Methods) we obtained high-quality, joint transcriptomic and chromatin accessibility datasets for 9’387 nuclei (**Fig. S5A, B**). Weighted nearest neighbor (WNN) integration of snATAC-seq and snRNA-seq data followed by Louvain clustering delineated 20 cardiac cell (sub)type clusters, alongside 8 non-cardiac clusters predominantly from heart-adjacent tissues, with discrete temporal profiles, as classified by gene set enrichment and marker gene analysis^18,32,84–87^ (**Figs. 3B, C; S5C-G, Tables S6, S7**). Specifically, we identified respective “early” (e; E9.5 or E10.5) or “late” (l; E10.5 or E11.5) clusters for ventricular CM (vCM), endocardial (EC), epicardial (EP), posterior SHF (pSHF) and cardiopharyngeal neural crest-derived (cpNCC) populations (**Figs. 3B, C; S5D-G**). We further observed two clusters exhibiting vascular endothelial cell-like (VEC) profiles^50^ present across all three timepoints, and three clusters hallmarked by smooth muscle cell and fibroblast -like (SM/FB) signatures detected only at E11.5 (**Figs. 3B, C; S5D-G**). This temporal restriction likely reflects the heterogeneous origins of these cell types^9^ and coincided with the emergence of SM cells during OFT remodelling^88^ as well as differentiation of EP-derived cardiac FBs^89^. Separate myocardial clusters including all three temporal layers comprised AVC, OFT, and atrial CM (aCM) populations (**Figs. 3B, C; S5D–G**), the latter also containing SAN progenitors marked by *Shox2* and *Hcn4*^43,53^ (**Fig. S5G, Table S6**). Another endocardial-associated (Endo) cluster displayed a characteristic profile of regulators involved in endocardial-to-mesenchymal transition (EndMT) and mesenchymal identity within the AV and OFT cushions, such as *Twist1*, *Sox9*, and *Tbx20*, along with the AV mesenchyme marker *Postn*^29,85^ (**Figs. 3B, C; S5D-G**). Finally, a specific TF signature including *Tbx1*, *Tlx1* and *Lhx2* defined a smaller cluster of cells with cardiopharyngeal mesoderm (CPM) identity^87,90^ (**Figs. 3B, C, S5D-G, Table S6**). As expected, both *Gata4* and *Hand2* signatures were prominent in myocardial clusters, including atrial and “early” (E9.5) or “late” (E10.5/E11.5) ventricular CM as well as AVC clusters (**Figs. 3C; S5F, G**). Consistent with the IF results (**Fig. S1E**), *Gata4* expression levels were higher than those of *Hand2* across CM subtypes, while both factors showed strong signals in EC and EC-derived EndMT clusters (**Figs. 3C, S5G**), in line with their respective roles in endocardial lineage regulation^30,51^ and initiation or maintenance of EndMT gene networks required for EC cushion formation^29,72^. In contrast, *Gata4* showed predominant expression in pSHF and EP clusters, whereas *Hand2* was more abundant in aSHF/OFT and cpNCC populations, reflecting distinct functions in other cardiac lineages^29,59,91,92^ (**Fig. 3C, S5G**).

**Figure 3:**
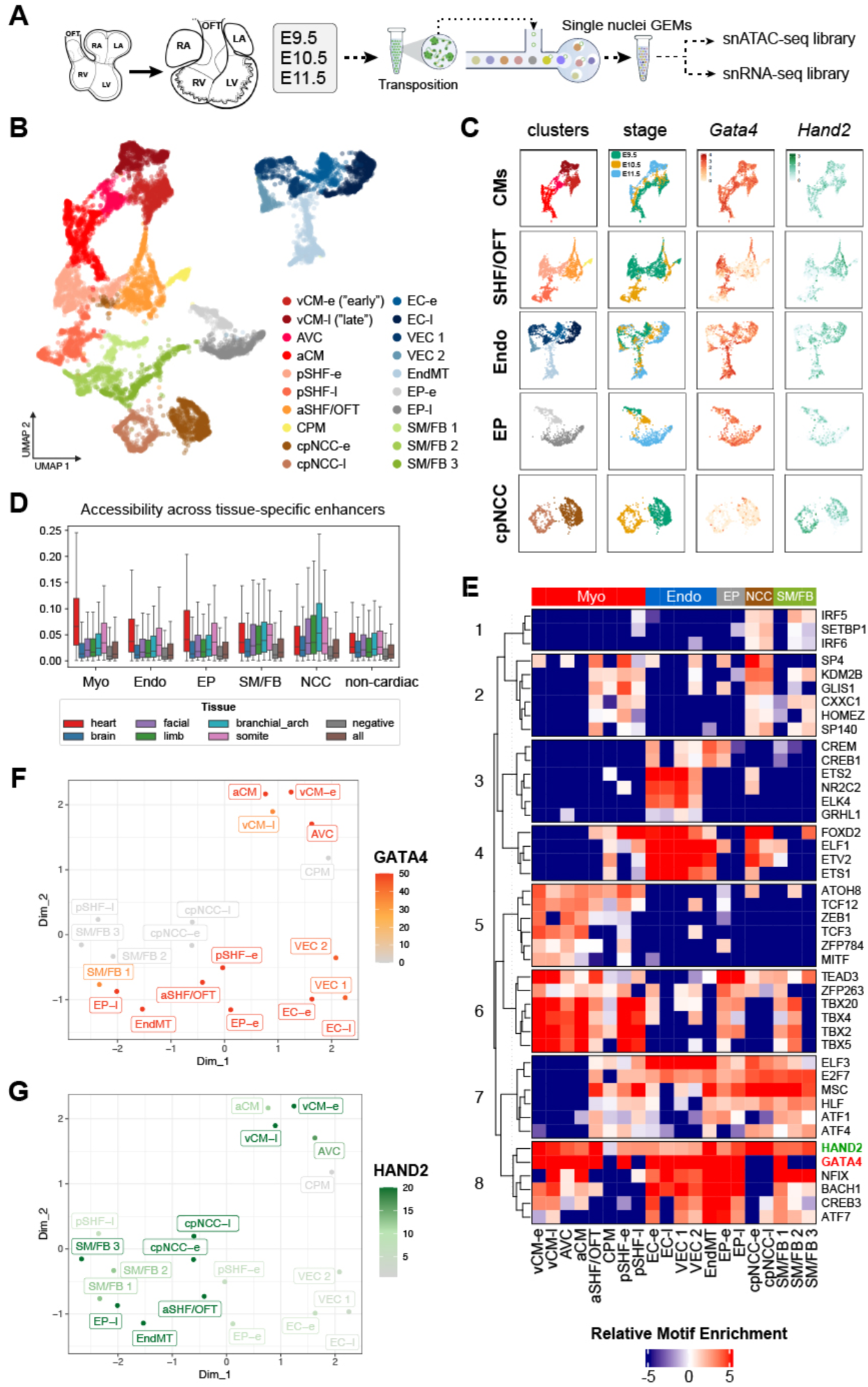
Single-cell multiome profiling of embryonic hearts delineates enhancer landscapes and cistromes across cardiac cell types. (**A**) Schematic of the single-nuclei multiomic (snMultiome) profiling approach. (**B**) UMAP showing nuclei assigned to the 20 cardiac clusters identified. (**C**) Clusters from (B) illustrating subsets of isolated cardiac cell types (rows) for which temporal signatures (stage) and selected TF expression profiles (*Gata4*/*Hand2*) are shown (columns). (**D**) Boxplots showing the average accessibility (snMultiome; split by lineage) of previously validated *in vivo* enhancer regions (VISTA Enhancer Browser), grouped by tissue of detected activity (**Table S8**). (**E**) Heatmap listing TF motifs with the highest differential enrichment across cistromes of lineage-specific clusters (**Table S10**). (**F, G**) UMAPs highlighting the similarities between cell clusters in the cistrome space (see Methods), color-coded by the significance of the enrichment (-log10 of the *p*-value) for the indicated TF-binding sites (GATA4, red; HAND2, green). OFT, outflow tract. RV, right ventricle. LV, left ventricle. RA, right atrium. LA, left atrium. a/vCM, atrial/ventricular cardiomyocytes. AVC, atrioventricular canal. a/pSHF, anterior/posterior second heart field-derived cells. SM/FB, smooth muscle/fibroblast-like cells. EP, epicardial cells. cpNCC, cardiopharyngeal neural crest-derived cells/mesenchyme. EC, endocardial cells. VEC, vascular-endothelial cells. EndMT, endocardial-to-mesenchymal transition/transformed cells of the cushions. -e, early (E9.5 or E10.5). -l, late (E10.5 or E11.5).

Analysis of chromatin accessibility profiles for each identified cell cluster yielded on average 76’822 peaks (range: 17’703-115’656), of which 5-30% showed a statistically significant bias towards higher accessibility in one or more cell clusters (**Fig. S6A, B, Table S7**). In line with previous reports^48,93^, as compared to promoters and CTCF-bound sites, TSS-distal enhancers tended to be more dynamically regulated across cell clusters (**Fig. S6C**). We next examined whether previously validated *in vivo* enhancers (VISTA Enhancer Database)^94^ exhibited specific patterns of chromatin accessibility within these newly defined regulatory maps. Compared with enhancer elements active in non-cardiac tissues, heart enhancers showed the highest chromatin accessibility in cell populations annotated as myocardial (a/vCM, AVC, aSHF/OFT, pSHF and CPM clusters) followed by those assigned to endocardial (EC, VEC, EndMT) and epicardial (EP) lineages (**Fig. 3D, Table S8**). Consistent with the shared neural crest lineage origin of craniofacial mesenchyme and cpNCC-derived cardiac populations^18^, enhancers previously validated *in vivo* for branchial arch activity exhibited highest accessibility in cpNCC clusters (**Fig. 3D, Table S8**). Collectively, these observations confirmed the overall specificity and *in vivo* relevance of the generated multiomic ATAC-seq datasets. Given the presumed relevant role of cardiac cell type-specific *cis*-regulatory elements in the manifestation of CHD^40,41,95^, we next examined the relationship between previously reported SNPs linked to CHD or other fetal/adult heart phenotypes (GWAS catalog) and our cell cluster-specific chromatin accessibility patterns. Whereas broader phenotype categories (e.g., PR interval) showed enrichment within regulatory elements active in CMs and more generally myocardial clusters, the analysis of CHD-associated SNPs did not yield a discernible or consistent regulatory pattern (**Fig. S6D, Table S9**). We next carried out a similar analysis using *de novo* variants identified by whole genome sequencing (WGS) in large cohorts of individuals with CHD^39,50,95^. Unexpectedly, but consistent with a comparable analysis of human snATAC-seq data^50^, we did not detect any individual or subset of clusters, whose putative regulatory elements showed an enrichment for these variants (**Fig. S6E, Table S9**). This result could be attributed to several factors, including an excess of non-functional elements annotated via mutiomics approaches, with most variants being inconsequential to significant gene dosage changes or equally distributed across most of the cell types profiled. To more precisely delineate the GRN architecture underlying heart morphogenesis, we next sought to infer the TF complement associated with the cistrome of each cell type cluster through stringent motif enrichment analysis (see Methods). This approach yielded hundreds of associations and highlighted the most significant TFs for each lineage, thereby providing a map of lineage and cell type-specific regulatory drivers (**Fig. 3E, Table S10)**. Importantly, each cluster showed strong enrichment for motifs associated with well-established cardiac lineage and cell type regulators (**Table S10**). Clustering of TF motif enrichment patterns across different cell types reflected the expected lineage relationship between CM and endocardial/endothelial populations, while cell types linked to OFT formation and remodelling (cpNCC, aSHF/OFT, and CPM) as well as mesenchymal transition/identity (EndMT, EP, SM/FB) grouped together (**Fig. S6F**). GATA4 and HAND2 consistently ranked among the top enriched TF motifs in myocardial and endocardial clusters (**Fig. 3E, S6G**), yet they also exhibit representation across other cell types consistent with their respective cardiac functions, such as EPs (*Gata4*/*Hand2*) or cpNCCs (*Hand2*)^59,91,92^ (**Fig. 3E-G, Table S10**). In addition, motifs for GATA4 and HAND2 showed some of the highest pervasiveness of enrichments across the individual cell clusters identified in the multiome data, indicating that both factors might function as context-dependent pioneer TFs shaping cardiac cell type-specific chromatin landscapes^51,96^ (**Figs. S6H, I**). In conclusion, our results enable the systematic exploration of cardiac cell type cistromes and regulatory element-to-target gene correlations through established gene-enhancer connectivity, thereby providing mechanistic resolution into the role of distinct TFs as key nodes in cardiac GRNs.

### Decoding the *Hand2* cardiac enhancer landscape at cell type resolution

To better delineate the *cis*-regulatory architecture of cardiac TFs at cell type resolution, we mined our snMultiome data and built a ShinyApp that enabled us to (i) fine-map the chromatin accessibility landscape of cardiac TFs at cell type resolution, (ii) compare these profiles with predictions of regulatory element identity provided by the ChromHMM interface^97^, and (iii) track down *cis*-regulatory elements correlating with gene transcription profiles based on computation of “gene-enhancer links” (**Fig. S7A, B**). To determine the effectiveness of this approach in functionally disentangling cardiac cell type-specific enhancer landscapes for core cardiac TF genes, we focused on the *Hand2* locus. Despite its essential functions across all major cardiac lineages and its involvement in regulating cardiac cistromes (**Fig. 3E–G**), the genomic enhancer landscape integrating cardiac cell type-specific upstream TF cues has remained unclear (**Figs. 1, S1B**). *Hand2* resides within a large TAD encompassing an extensive upstream genomic interval of nearly 400 kb devoid of genes or transcripts, extending from the *Uph*/*Hand2os1* lncRNA^98,99^ to the far upstream transcript *Gm33501*, and containing at least six limb-specific as well as additional non-cardiac enhancers^100–102^ (**Fig. 4A**). Interrogation of this region with our multiomic framework identified more than 50 accessible chromatin elements with activity in at least one of the 28 defined multiome clusters (**Fig. S5**), including the previously identified heart and limb enhancer elements^60,63,100,101^ (**Figs. 4A, S7A, Table S11**). Among these, mapping of inferred “gene-enhancer” links uncovered a subset of nine distal *Hand2*-associated heart enhancer candidates (I-IX) (**Figs. 4A, S7A**). These elements showed specific accessibility patterns across *Hand2*-expressing cardiac clusters, with combinatorial signatures present across CM, EC, EP and cpNCC lineages (**Figs. 3C, 4A**). Most elements also lacked accessibility in non-cardiac cell types, and none overlapped with previously identified limb enhancers^100,101^ (**Fig. 4A**). We next defined a set of high-confidence cardiac enhancer candidates by selecting elements with the most pronounced ATAC accessibility across cardiac lineages, including elements with rather broad (-437, -397) or cell type restricted (-389, -307, -68) profiles (**Fig. 4A**). Using enSERT for site-directed transgenic reporter analysis to determine respective *in vivo* enhancer activity patterns in mouse embryos at E10.5 revealed remarkable predictivity of the multiome signatures, as the prioritized elements exhibited cardiac compartment- or cell type-specific enhancer activity largely consistent with multiomic chromatin accessibility profiles (**Figs. 4B; S8A, B, Table S12**). While the heart enhancers located 437 kb (hE1) and 397 kb (hE2) upstream of the *Hand2* TSS drove overlapping activity in the OFT and RV myocardium, hE2 (-397), also showed activity in the EC population and EndMT-derived cells of the OFT and AVC cushions, in line with elevated chromatin accessibility in the corresponding clusters (**Figs. 4A, B; S8A, B**). The suitability of our framework, which integrates gene-enhancer link calls with multiome ATAC accessibility, was further supported by the correct prediction of cell type specificities for the elements located 389 kb (hE3), 307 kb (hE4), and 68 kb (hE5) upstream of *Hand2* (**Figs. 4A, B; S8A, B**). The observed reporter activities driven by these elements in the EP, EndMT (mesenchymal cushion), and EC populations, respectively, consistently matched the profiles of the most pronounced chromatin accessibilities for each region (**Figs. 4A, B; S8A, B**). For comparison, an additional element with cardiac enhancer-predictive ChromHMM signal near hE5 (-72), but without an inferred multiomic *Hand2*-correlation, failed to drive reproducible activity in the heart (**Fig. 4A**). To validate the broad applicability of our multiome resource for reliable identification of gene-linked *in vivo* cardiac enhancers, we also assessed whether the inferred multiome links could accurately recover the previously characterized heart enhancers at the *Gata4* locus^61,62,68^ (**Figs. 1B, S1A, S7B; Table S11**). Indeed, while all previously validated murine heart enhancers were present within the set of accessible regions correlated with *Gata4* expression (n=5/19), the functionally validated *G4*-124, *G4*-93 and *G4*+177 enhancers (**Figs. 1, 2**) ranked among the top-four associated elements, with the most pronounced multiomic accessibility profiles matching the observed cardiac cell type enhancer activities (**Figs. 1B, S1C, S7B**). Together, these findings demonstrate that our multiome-based framework provides a robust strategy to systematically identify *in vivo* heart enhancers, link them to target genes, and predict their cell type specificities.

**Figure 4:**
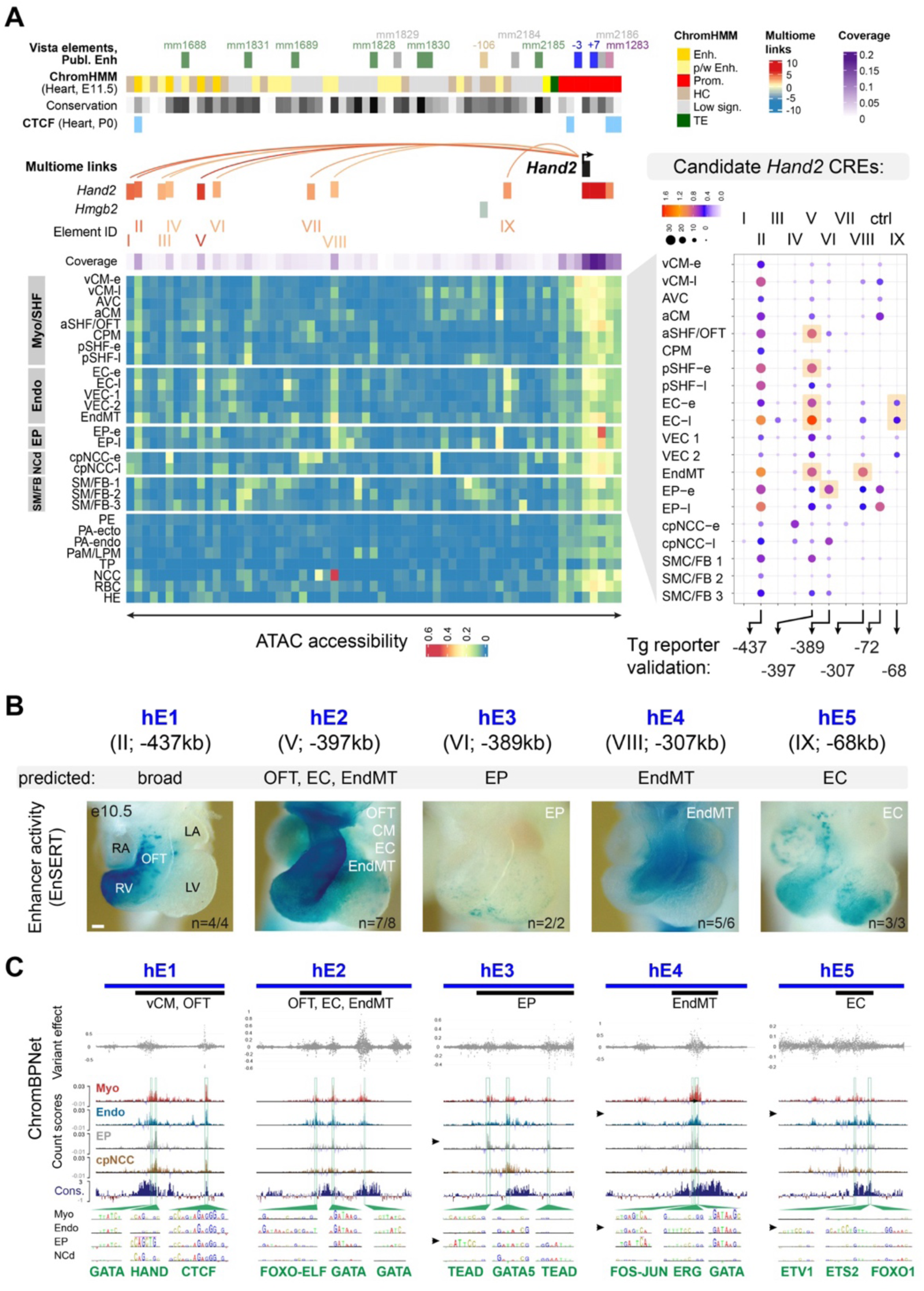
Multiome-guided discovery of a repertoire of cardiac compartment- and lineage-specific *Hand2* enhancers. (**A**) Multiomic prediction of cardiac cell type enhancers within the extensive (430kb) *Hand2* upstream domain. All putative regulatory elements accessible in at least one multiome cluster are listed from left to right (**Table S11**). Per element, ATAC coverage across all clusters is shown in purple shade. Top: Color-coding in matrix indicates previously validated elements by tissue-specific activity (blue, heart; green, limb; purple, craniofacial; light brown, stomach; gray, no activity), ChromHMM cis-regulatory element (CRE) annotations^97^, conservation (black, high; white, low) and CTCF enrichment (peaks) in hearts^70^. P0, postnatal day 0. Gene-enhancer links were computationally inferred from multiome data by considering all genes within the corresponding TAD and *Hand2*-correlated regulatory elements (I-IX, distance from the *Hand2* TSS indicated in brackets) exhibiting strong chromatin accessibility (highlighted in yellow) served as a basis for prediction of cardiac compartment or cell type -specific enhancer activities (**Table S11**). Ctrl, control region (*H2*-72kb; **Table S12**). Dot color and size represent ATAC accessibility and percentage of cells across clusters showing accessibility, respectively. CREs, cis-regulatory elements. (**B**) Site-directed LacZ reporter transgenesis^69^ validates cardiac in vivo enhancer activities consistent with multiomic predictions. Reproducibility numbers (n) are indicated on the bottom right of each representative embryo shown (reproducible tissue-specific staining vs. number of transgenic embryos with any LacZ signal). Scale bar, 100μm. hE, heart enhancer. (**C**) ChromBPNet predictions of variant effects and count contribution scores for identification of relevant TF motif instances^103^. Count contribution scores are shown for overarching cluster groups (Myo, Endo, EP, cpNCC) as defined in (A), with strongest predictive motif instances in cell types marked by enhancer activity (black arrows) and corresponding TFs (green) highlighted. Enhancer IDs are assigned based on their upstream distance (in kb) from the *Hand2* TSS or according to the VISTA Enhancer database (mm, Mus musculus). OFT, outflow tract. RV, right ventricle. LV, left ventricle. RA, right atrium. LA, left atrium. a/vCM, atrial/ventricular cardiomyocytes. AVC, atrioventricular canal. a/pSHF, anterior/posterior second heart field-derived cells. CPM, cardiopharyngeal mesoderm. EC, endocardial cells. VEC, vascular-endothelial cells. EndMT, endocardial-to-mesenchymal transition/transformed cells of the cushions. EP, epicardial cells. cpNCC, cardiopharyngeal neural crest-derived cells/mesenchyme. SMC/FB, smooth muscle/fibroblast-like cells. PE, pulmonary endoderm. PA, pharyngeal arch. PaM, paraxial mesoderm-derived cells (skeletal muscle). TP, thyroid progenitor-like cells. RBC, red blood cells. HE, hematopoietic (progenitor) cells. -e, early. -l, late. E, embryonic day. -e, early. -l, late.

To pinpoint cardiac lineage regulators in control of the identified *Hand2* upstream enhancers, we applied ChromBPNet, a deep learning model able to predict base-resolution chromatin accessibility revealing relevant nucleotide-encoded TF motifs and for prioritization of enhancer-associated variants^103^. All enhancer elements showed direct sequence conservation of core regions, with hE1, hE2, and hE4 exhibiting a remarkable degree of conservation down to *Coelacanth* (**Fig. S9A**), while hE3 and hE5 showed conservation restricted to mammals (**Fig. S9A**). Application of ChromBPNet across aggregates of major cell lineages revealed distinct, cardiac lineage-specific features predominantly within these conserved enhancer cores, highlighting sensitivity to variant-driven modulation (**Fig. 4C**). These sensitivity intervals overlapped with count scores, which quantify the contribution of each base sequence to the predicted overall accessibility, thereby marking the positions of putative critical TF motifs^103^ (**Fig. 4C**). The ChromBPNet-predicted functional motifs within these high-sensitivity nucleotide regions indeed corresponded to cardiac lineage- or cell type TF regulators, largely consistent with their respective roles in the assigned cardiac cell clusters (**Fig. 4C**). For example, HAND and GATA motifs were discovered within the hE1 enhancer with predominant OFT/RV activity, while GATA5, FOS-JUN/ERG and an ETV1/ETS2/FOXO1 signatures were detected in EP (hE3), EndMT (hE4) and EC (hE5) -specific enhancers (**Fig. 4C**). The pronounced enrichment of GATA motifs at key positions within the analyzed enhancers likely reflects the broad role of GATA factors in initiating and shaping myocardial and endocardial lineage-specific enhancer landscapes^51,74^ and may reflect the regulatory wiring underlying the upstream requirement of *Gata4* for *Hand2* expression in *Nkx2-5*^+^ myocardial progenitors^27^. In line with the distal CTCF peak flanking the hE1 conserved enhancer core (**Fig. S9A**), ChromBPNet analysis also identified a corresponding CTCF motif (**Fig. 4C**). Given that the entire *Hand2* upstream non-coding interval lacks additional CTCF enrichment (**Fig. S7A**), this far-upstream element may act as a distal domain anchor insulating interactions between *Hand2* and its upstream enhancers.

To evaluate the ChromBPNet predictions through physical presence of respective cardiac TFs at these enhancers, we reprocessed available ChIP-seq and ATAC-seq datasets from embryonic hearts^51,72,74,75^ (**Fig. S9, Table S13**). In accordance with the presence of GATA4 motifs within the conserved cores of hE1, hE2 and hE4 elements (**Fig. 4C**), we identified highest level of GATA4 occupancy precisely at these enhancers (**Fig. S9A**). Moreover, in agreement with the detection of an active HAND motif within the hE1 core (**Fig. 4C**), our analysis confirmed the presence of HAND2 in the same location in hearts at E10.5^72^ (**Fig. S9A**), indicating potential feedback activity. TBX5, a key regulator of myocardial activity was further present exclusively at hE1 (**Fig. S9A**). In accordance with the endocardial specificity of the hE5 enhancer from our multiome predictions and the results from EnSERT *in vivo* reporter analysis (**Figs. 4A, B; S8**), hE5 was showing chromatin accessibility in ECs but not the myocardial population at E12.5^51^ (**Fig. S9B**). Interestingly, absence of *Gata4* in hearts at E12.5^51^ did not significantly alter chromatin accessibility at any of the enhancer elements (**Fig. S9C**), suggesting pioneering activity at an early stage or alternative factors involved. Finally, GATA4 continues to physically interact with these enhancers in adult mouse ventricles and atria^104^ (**Fig. S9D**), suggesting a putative role in *Hand2* regulation beyond developmental stages. Collectively, our results indicate that distinct combinations of cardiac lineage-specific TFs, such as ETS/ETV or HAND factors, in conjunction with broadly expressed cardiac identity factors such as GATA4, act cooperatively on *Hand2* heart enhancers to establish or maintain the functional properties defining cardiac cell types.

### Polymer modelling defines cardiac-specific regulatory architecture

To characterize the 3D physiological context encompassing the *Hand2* cardiac cell type enhancer landscape and to investigate chromatin features of cardiac specificity, we conducted Capture Hi-C (C-HiC)^53,105^ on a 3.5 Mb genomic region encompassing the *Hand2*-containing and adjacent TADs in mouse embryonic hearts (HT), forelimbs (FL), and mandibular processes (MD) at E11.5, tissues predominantly affected by *Hand2* LOF^83,106^ (**Fig. 5A**). Analysis of insulation scores^107^ from HT, FL, and MD datasets revealed that while the *Hand2* gene body resides within a sub-TAD boundary in all three tissues, a distinct TAD boundary located approximately 435 kb upstream of Hand2 was detected exclusively in HT, but not in FL or MD tissues (**Fig. 5A**). To pinpoint elements that strongly interact with the *Hand2* promoter, we computed normalized virtual 4C (nv4C) profiles and sub-TAD boundaries for each tissue type, using the *Hand2* genomic region as the viewpoint (**Tables S14, S15)** (see Methods). This analysis identified a region located approximately 435kb upstream of the *Hand2* TSS (*H2*-435) harboring the hE1 OFT/RV enhancer and an adjacent CTCF-interacting element (**Figs. 4, 5A, S10A**). We thus hypothesized that this region representing a pronounced *Hand2* interaction site might act as a principal domain anchor in the heart (designated *H2*-HT) (**Figs. 5A, S10A**). In contrast, the most pronounced *Hand2* interaction site in both FL and MD tissues (*H2*-FL/MD) was located further upstream (-685kb) at an element enriched for H3K27ac in limb tissue and flanked by multiple CTCF peaks present in limb but not heart tissue (**Figs. 5A, S10A**). Prominent interactions between *Hand2* and the *H2*-HT element were also observed in MD/FL tissues, likely associated with the activity of a tissue-invariant CTCF site (**Fig. S10A**). To test reciprocal interactions of the *Hand2* promoter, we generated nv4C tracks using the identified upstream heart enhancers (hE1-hE5) as viewpoints (**Figs. 4, 10B, Table S14**). A viewpoint centered on the hE1 (*H2*-437) module indeed corroborated strongest interaction between this element with the *Hand2* promoter in the heart, with higher nv4C interaction counts than in FL or MD tissues (**Fig S10B**). Similarly, other far-upstream cardiac enhancers (hE2-hE4) engaged in pronounced contacts with *Hand2*, with interaction frequencies markedly enriched in heart tissue relative to FL and MD samples (**S10B**). In contrast, the more proximally located hE5 enhancer (*H2*-68) exhibited most pronounced contacts predominantly with a region upstream of the *H2*-HT anchor (**Fig. S10B**). Taken together, these analyses imply that the *Hand2* upstream regulatory domain exhibits distinct regulatory interaction dynamics in heart versus FL and MD tissues. In support, the previously identified mm1687 strong limb enhancer^100,101^ (*H2*-488) is located outside of the putative *H2*-HT domain interval but included in the *H2*-FL/MD domain anchored at 685kb upstream of *Hand2* (**Fig. 5A, S10A**).

**Figure 5:**
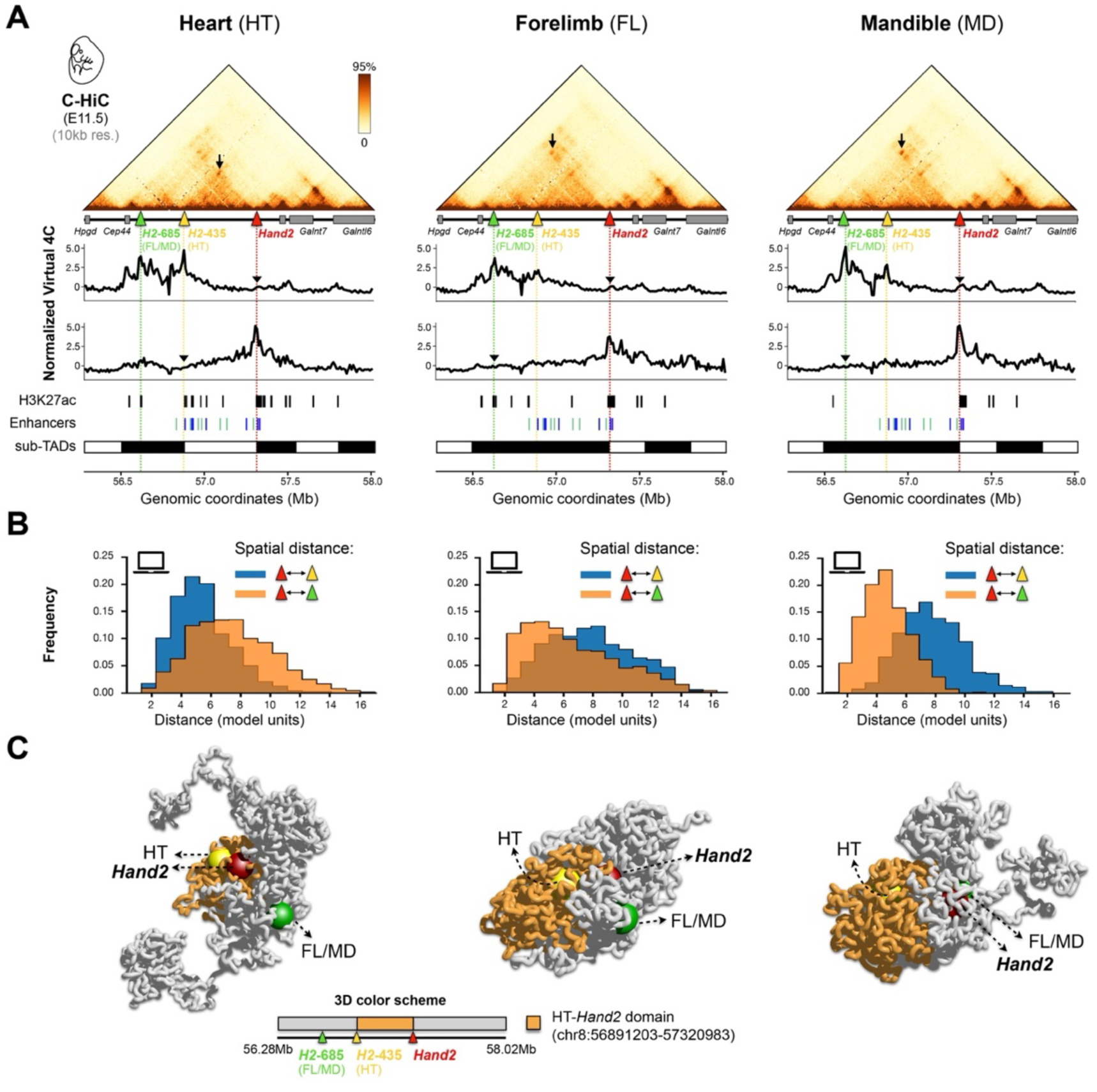
Chromatin polymer modeling uncovers cardiac regulatory domain organization at the *Hand2* locus. (**A**) Top: region-capture HiC (CHi-C) from heart (HT), forelimb (FL) and mandible (MD) tissues at E11.5 across a genomic interval containing the *Hand2* TAD (mm10, chr8:56281203-58021203; 10kb resolution data). Black arrow indicates the strongest anchor point of the prominent interaction domain upstream of the *Hand2* gene (red triangle) in all three tissues. Middle: normalized virtual 4C tracks (see Methods) using either *Hand2* (upper panels) or the strongest anchor element (lower panels) as viewpoint (black arrowhead). Bottom: Comparison to tissue-specific H3K27ac enrichment profiles (ENCODE)^70^, newly identified (Fig. 4) or previously validated enhancers active in heart (blue), limb (green), and craniofacial (purple) tissues^60,63,68,100,101^ (VISTA Enhancer Browser), and sub-TAD domains derived from HT, FL, and MD C-HiC datasets (**Table S15**). (**B**) Spatial distances between *Hand2* and its most prominent interaction sites in HT, FL and MD tissues, as computed from single molecule 3D structures using the Strings and Binders (SBS) model^168^ (see Methods). (**C**) Tissue-specific single-molecule SBS 3D-reconstructions revealing closest spatial association of *Hand2* (red) with the *H2*-435 element (yellow) in the heart, and with *H2*-685 (green) in FL and MD tissues.

To further explore the 3D spatial organization at the *Hand2* locus across tissues, we applied the Strings and Binders (SBS) model^108^, a polymer physics-based framework which has been previously shown to explain chromatin contact (e.g., Hi-C and GAM) and single-cell microscopy data both at specific genomic regions^109–111^ and genome-wide^112^. To construct SBS polymer models for each selected tissue, we applied the PRISMR method, which infers the minimal set of distinct binding site types and their spatial arrangement along the polymer chain using tissue-specific C-HiC interaction data as input^112,113^ (Methods). This analysis returned a polymer model with nine different types of binding sites (model binding domains) for each of the tissues analyzed (**Fig. S10C**). Notably, many identified binding domains displayed specific spatial distribution patterns across tissues, with heart tissue showing a distinct central domain extending from the *Hand2* gene body to the *H2*-435/hE1 domain anchor (**Fig. S10C**). In contrast, the central domains in FL and MD showed a greater extension, with the signal reaching to the more distant *H2*-685 element (**Fig. S10C**). Together, these results indicated dynamic tissue-specific compartmentalization of chromatin structure at the *Hand2* locus. To refine these tissue-specific 3D configurations predicted by the SBS model, we computed the distributions of pairwise spatial distances between the *Hand2* gene and its most prominent interaction sites in heart (*H2*-435) or FL and MD (*H2*-685) tissues (**Fig. 5B**). Analysis of distance distribution models in FL and MD revealed that the *Hand2* gene body was consistently positioned closer to the *H2*-685 interaction site than to the *H2*-435/hE1 element, whereas in heart the opposite scenario was identified (**Fig. 5B**). Consistently, single-molecule conformations of the model revealed close 3D proximity of *Hand2* and the *H2*-435 element in cardiac tissues, further supporting a role as loop anchors stabilizing a regulatory domain (**Fig. 5C**). In contrast, *H2*-685 was positioned farther from *Hand2* in the heart but showed closer proximity in FL and MD models (**Fig. 5C**). Hence, consistent with our nv4C analysis, our polymer model suggests confined interactions between elements within the noncoding upstream domain and *Hand2*, supported by a preferential 3D chromatin configuration anchored through strong contacts between the *H2*-435 site and the *Hand2* promoter region (HT domain). These findings support the existence of tissue-specific differences in the 3D genomic architecture within the *Hand2* TAD, which in the heart may facilitate *Hand2* communication with the distant upstream heart enhancers (hE1-hE5).

### The *Hand2*-upstream regulatory interval controls endocardial development and embryonic survival

Given the cardiac compartment- and cell type-specific, yet partially overlapping, activities of the identified upstream *Hand2* heart enhancers, we hypothesized that these regulatory modules are essential for controlling the dynamic, cell type-specific expression of *Hand2* and, collectively, are critical for heart development. To evaluate whether deletion of the non-coding *Hand2* upstream regulatory interval (termed *H2*-URI), delimited by the CTCF-containing *H2*-435 domain anchor at the 5′ end and the *Uph* lncRNA adjacent to the *Hand2* TSS at the 3′ end^98^, alters chromatin domain architecture, we performed Molecular Dynamics simulations using SBS polymer models of the *Hand*2 locus in heart, FL, and MD tissues (**Figs. 6A, S11A**). Thereby, the deletion was incorporated into the wild-type models to recompute the three-dimensional spatial organization (see Methods). The resulting model-predicted 3D conformations indicated that, in the absence of the 373 kb-spanning *H2*-URI, the basic 3D chromatin structure of the *Hand2* regulatory locus was maintained (**Fig. S11A**). Therefore, considering the predicted integrity of putative domain anchors in the absence of the *H2*-URI, we used CRISPR/Cas9 to generate a mouse model harboring the respective 373 kb deletion of endogenous non-coding sequence (*H2*-URI^Δ^) to assess the collective contribution of the distant *Hand2* upstream cardiac enhancers to heart development (**Figs. 6A, S11B**). The deletion included hE2, hE3, hE4, and hE5, each of which drove activity in specific cardiac compartments and/or cell-type populations known for critical *Hand2* functions, such as RV/OFT progenitors (hE2), the endocardium (hE5), EndMT-derived mesenchymal cells of the OFT and AVC cushions (hE2 and hE4), and EPs (hE3)^25,30,32,72,92^ (**Figs. 4, 6A, S8**). Following the establishment of founders, *H2*-URI^Δ/+^ mice were born at expected Mendelian ratios showing neither impaired viability nor fertility (**Fig. S11B**). However, when *H2*-URI^Δ/+^ mice were intercrossed to produce embryos lacking the *H2*-URI, all homozygous (*H2*-URI^Δ/Δ^) littermates (n = 5/5) exhibited consistent embryonic lethality and developmental arrest around embryonic day E10–E10.5 (**Figs. 6B, S11B**). In comparison, *Hand2* deletion in isogenic (FVB) background resulted in earlier embryonic lethality at E9.25 (n = 4/4) (**Fig. 6B**), consistent with previous studies linking the cause of lethality to RV hypoplasia^25,28,32^. In contrast, molecular analysis comparing *H2*-URI^Δ/Δ^ embryos and WT controls at E9.5 uncovered neither morphological abnormalities at the whole mount level nor evidence of excessive apoptosis in the heart (**Fig. S12A, B**). Instead, spatial transcript analysis by ISH revealed a clear reduction of *Hand2* in atrial and ventricular chambers, as well as the more caudal part of the OFT, in *H2*-URI^Δ/Δ^ embryos at E9.5 compared with WT controls (**Fig. 6C, S12A**). Accordingly, qPCR analysis confirmed significant *Hand2* gene dosage reduction (∼70%) in mutant embryos at the same stage (**Fig. 6D**). *Hand2* transcripts persisted in the upper part of the OFT as well as the branchial arches, but with marked reduction in the mandibular arch (MA) (**Fig. 6C, S12A**). Analysis of HAND2 protein distribution at E9.5 further revealed absence of HAND2 from the endocardial (EC) population in atrial and ventricular chambers, as well as the OFT in *H2*-URI^Δ/Δ^ embryos (**Fig. 6E, S12C**). We further confirmed the absence of HAND2 from the endocardial lining in cardiac compartments by co-localization with the endocardial/endothelial marker CD31 (PECAM1) (**Fig. S12D**). In addition, HAND2 was markedly reduced in portions of the RV myocardium and posterior OFT (**Fig. 6E, S12C**), which may contribute to the embryonic lethality observed in *H2*-URI^Δ/Δ^ littermates. GATA4, which has been described to act upstream of HAND2 in SHF-related GRNs^114,115^, was not markedly changed in the HAND2-deficient endocardium, EC-derived cushions or OFT cells at E9.5 (**Fig. 6E, S12C**). Together, these results align with the observed downregulation of *Hand2* in cardiac chambers and indicate an essential role for the *H2*-URI-encoded heart enhancers, particularly those with activities in the endocardium and EC cushion mesenchyme (hE2, hE4, and hE5). Given that GATA4 remained present in ECs even in the absence of HAND2 (**Fig. 6E, S12C**), and that it physically occupied these enhancers (**Fig. S9**), our findings support an upstream role of GATA4 in regulating endocardial *Hand2* through the *H2*-URI.

**Figure 6:**
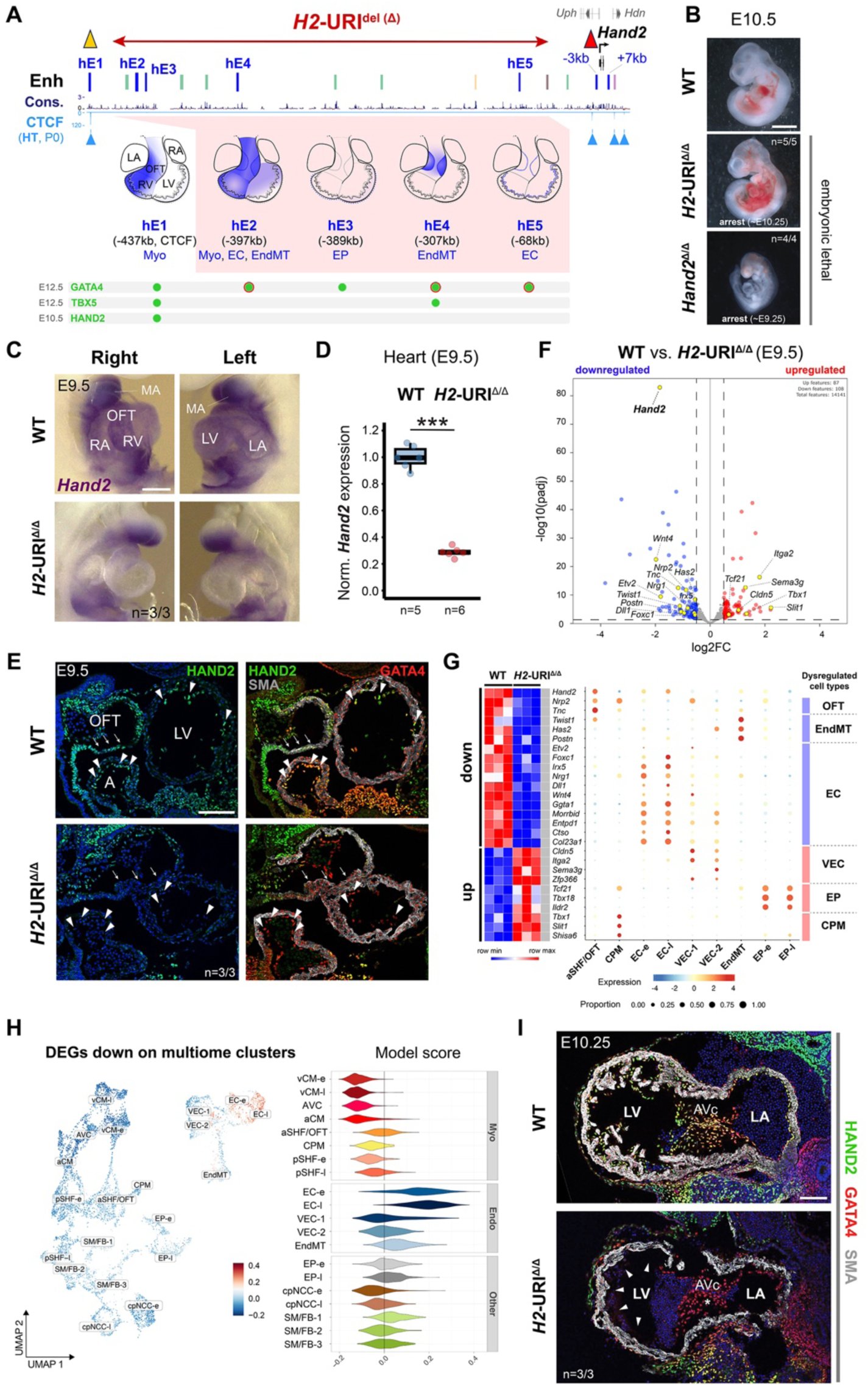
The *Hand2* upstream regulatory interval (URI) integrates cardiac gene networks essential for endocardial development. (**A**) Engineering of a 373kb-spanning endogenous deletion (chr8:56903578-57277013, mm10) in mice (*H2*-URI^Δ^ allele) to remove the regulatory interval within the designated upstream chromatin domain of *Hand2* (Fig. 5). hE1-hE5: identified upstream *Hand2* heart enhancers driving cardiac compartment- and cell type-specific activities (Fig. 4). Previously validated enhancers with activity in the heart (blue), limb (green)^100,101^, stomach (yellow)^102^, midbrain (brown) (mm2187) or craniofacial tissues (purple) (mm1283) are listed. CTCF peaks in hearts at P0 (ENCODE)^70^ are marked by light blue triangles. The interaction of cardiac TFs in hearts (green dots) and endocardium (red outline) is indicated (see **Fig. S9**). Cons: vertebrate conservation track by PhyloP. Distance to *Hand2* TSS (+/-; in kb) is indicated for heart enhancers. (**B**) *H2*-URI^Δ/Δ^ embryos exhibit developmental arrest followed by embryonic lethality around E10.5. Hand2^Δ/Δ^ embryos in isogenic FVB strain background arrest around E9.25 due to right ventricle (RV) hypoplasia confirming previous results^25,83^. Scale bar, 500 μm. (**C**) ISH reveals reduction of *Hand2* transcripts in *H2*-URI^Δ/Δ^ embryonic hearts at E9.5 compared to WT controls. MA, mandibular arch. Scale bar, 250 μm. (**D**) Significantly reduced *Hand2* mRNA in hearts of *H2*-URI^Δ/Δ^ embryos compared to wildtype (WT) controls at E9.5. Box plots indicate interquartile range, median, maximum/minimum values (bars) and individual biological replicates (n). P-value is shown, with ***P < 0.001 (two-tailed, unpaired t-test). (**E**) Co-localization of HAND2 (green) and GATA4 (red) in hearts of *H2*-URI^Δ/Δ^ compared to WT embryos at E9.5. Arrowheads point to loss of HAND2 in endocardial cells (ECs) in mutants, while GATA4 is retained. Arrows point to reduction of HAND2 in the outflow tract (OFT). Scale bar, 100 μm. (**F**, **G**) Bulk RNA-seq from *H2*-URI^Δ/Δ^ embryonic hearts at E9.5 compared to WT controls (n=3 replicates per genotype). Differentially expressed genes (DEGs) with FDR < 0.05 and log2FC > 0.5 (F) and affected cardiac cell type markers/regulators with multiome snRNA-seq profiles (see **Fig. S13** for top 50 down/up-regulated DEGs). (**H**) Left: UMAP projection of downregulated bulk RNA-seq DEGs from *H2*-URI^Δ/Δ^ vs. WT mouse embryonic hearts at E9.5 onto aggregated cardiac cell type clusters from single-cell multiome analysis (Fig. 3). Color shades indicate over- (red) and under-represented (blue) expression of the downregulated DEGs, in each cell. Right: violin plots showing the distribution of these values, per cluster. See **Fig. S13** for projection of upregulated DEGs. (**I**) Co-localization of HAND2 (green) and GATA4 (red) in hearts of *H2*-URI^Δ/Δ^ and WT embryos at E10.25. Arrowheads denote the markedly diminished SMA⁺ (gray) myocardial trabeculae, whereas the arrow indicates the hypoplastic atrioventricular cushion (AVc) comprising HAND2⁻ EndMT-derived cells in *H2*-URI^Δ/Δ^ hearts. Scale bar, 100 μm. “n” indicates number of biological replicates analyzed, with similar results. LV, left ventricle. RA, right atrium. LA, left atrium. a/vCM, atrial/ventricular cardiomyocytes. AVC, atrioventricular canal. a/pSHF, anterior/posterior second heart field-derived cells. SM/FB, smooth muscle/fibroblast-like cells. EP, epicardial cells. cpNCC, cardiopharyngeal neural crest-derived cells/mesenchyme. EC, endocardial cells. VEC, vascular-endothelial cells. EndMT, endocardial-to-mesenchymal transition/transformed cells of the cushions. -e, early (E9.5 or E10.5). -l, late (E10.5 or E11.5).

To explore the necessity of the *H2*-URI at the transcriptional level and to define its downstream effect on cardiac cell type GRNs, we performed whole-mount RNA-seq on hearts from *H2*-URI^Δ/Δ^ and wild-type embryos at E9.5. Computational analysis revealed a total of 87 up- and 108 down-regulated genes, with *Hand2* being the most significantly down-regulated one (**Fig. 6F, S13A; Table S16**). Remarkably, the top 50 down-regulated differentially expressed genes (dDEGs) were found predominantly enriched for regulators of endocardial (EC) development, EC cushion (EndMT) cells and to a lesser degree OFT markers (**Fig. 6G, S13B, C; Table S16**). As corroborated by our multiome RNA-seq profiles, the complement of dDEG genes included *Etv2*, *Foxc1* and *Irx5* EC regulators as well as *Nrg1* (*Neuregulin-1*) acting downstream of *Hand2* to mediate endocardial-to-myocardial interactions essential for trabeculation initiation^30,116^ (**Fig. 6G, S13B**). In addition, several other genes with EC-specific expression profiles (e.g., *Dll1*, *Ggta1*, *Morrbid*, *Ctso* and *Col23a1*) were identified among dDEGs (**Fig. 6G, S13B**). Among the most significantly downregulated EndMT genes were *Twist1*, *Has2* and *Postn* which were previously shown to be dependent on *Hand2* in AVC development and harbored *HAND2* occupied regions in their regulatory domains^72^ (**Fig. 6G, S13B**). These results indicate that the absence of the hE5 EC-enhancer encoded in the *H2*-URI is likely a major determinant of *Hand2* downregulation in ECs, while EndMT genes are regulated likely through overlapping functions of hE2 and hE4 enhancers both driving activity in the OFT or AVC cushion mesenchyme (**Figs. 5, S8**). In addition, downregulation of myocardial OFT genes *Nrp2* and *Tnc* (**Fig. 6G, S13B**) aligned with the previously observed reduction of *Hand2*/HAND2 in the lower portion of the OFT (**Fig. 6D, E**), indicating functional contributions of the hE2 enhancer through its activity in the OFT myocardium (**Fig. S8**). RNA-seq analysis further revealed a specific complement of genes upregulated in the absence of the *H2*-URI (**Fig. 6F, S13A**). These upregulated DEGs (uDEGs) showed predominance of markers enriched in the clusters for vascular endothelial cells (VECs), including *Cldn5* and *Itga2*, as well as regulators of epicardial cells (EPs) such as *Tcf21* or *Tbx18*, and genes specifically expressed in the cardiopharyngeal mesoderm (CPM) population, such as *Tbx1* or *Slit1* (**Fig. 6G, S13C**). In line with the upregulated VEC signatures (**Fig. 6G**), a previous study observed upregulation of related coronary development genes in hearts deficient for endocardial *Hand2*, leading to a hypervascularized myocardium at E13.5^30^. In summary, our findings further suggest that regulatory elements within the *H2*-URI are necessary for the transcriptional control of *Hand2* in VECs, EP, and CPM populations. These elements may function as either enhancers or silencers, depending on whether *Hand2* deploys activating or repressive downstream function in a respective cell type.

To better visualize the cell type-specific affiliation of DEGs dependent on *H2*-URI function, we mapped dDEGs and uDEGs onto the cardiac multiome clusters previously generated (**Figs. 3**; **6H; S13D, E**). Consistent with the observed loss of HAND2 from the EC population, this revealed a marked overrepresentation of dDEGs in the EC-e and EC-l, as well as the EndMT cluster (**Fig. 6H**). In addition, comparison of average normalized expression of dDEGs projected to multiome clusters also indicated reduced dosage of genes belonging to cpNCC-e and aSHF/OFT clusters (**Fig. 13E**). Instead, projection of uDEGs showed an overrepresentation of genes not only in VECs, EPs, and CPM clusters, but also within ventricular/atrial and AVC myocardial populations, indicating additional perturbations across CM subtypes (**Fig. S13D**). In summary, these results corroborate the cardiac cell type-specific functions of the heart enhancers encoded within *H2*-URI and suggest a role in the transcriptional control of GRNs and gene sets that define the properties of distinct cardiac cell types^28,30,59^.

Finally, to elucidate the functional relevance of the *H2*-URI and its downstream GRNs on structural heart morphogenesis, we performed IF analysis of *H2*-URI^Δ/Δ^ and WT embryos at E10.25, a stage at which *H2*-URI^Δ/Δ^ embryos show embryonic arrest but the heart is not yet affected by the presence of apoptotic cells (**Figs. 6I; S12B**). In agreement with the loss of *Hand2*/HAND2 in ventricular ECs accompanied by downregulation of key EC mediators such as *Nrg1*^30,116^, we observed near absence of *Nrg1*-dependent myocardial trabeculae in the ventricular chambers of *H2*-URI^Δ/Δ^ embryonic hearts (**Fig. 6I**). In addition, consistent with the downregulation of EndMT regulators (e.g., *Twist1*)^117^ and genes required for proper expansion of EC cushions (e.g., *Has2* and *Postn*)^118,119^, we observed loss of HAND2 also in the *H2*-URI^Δ/Δ^ cushion mesenchyme which appeared hypoplastic compared to counterparts in WT embryos (**Figs. 6I**). Thus, these findings demonstrate that the *H2*-URI plays a central role in regulating *Hand2* transcription dynamics within EC and EndMT-derived cushion populations and suggest that the collective EC-related enhancer activities (hEC2, hEC4 and hEC5) are functionally required for proper trabeculation of cardiac chambers and for the expansion of EC cushions in the AVC. In this context, the identified heart enhancers shape a genomic hub that integrates cardiac lineage- and cell type-specific TFs acting within EC and EndMT GRNs associated with hypotrabeculation and tricuspid atresia CHD phenotypes^30^. Overall, and integrated with results from limb development^101,120^, our findings suggest that the *H2*-URI functions as an insulated, modular regulatory hub of ancient vertebrate origin, orchestrating heart and limb morphogenesis through distinct enhancers with partially overlapping compartment- and cell type-specific activities.

## Discussion

In this study, we examined the functional necessity of five individual cardiac enhancer modules associated with *Gata4* and *Hand2*, two core regulators of heart development that act in a coordinated manner across myocardial, endocardial, and epicardial lineages^5,14,115^. Although each factor plays a critical role in myocardial and endocardial lineages essential for RV/OFT morphogenesis^25,27,28,32^, as well as in trabeculation and cushion-mediated heart valve development, respectively^29,30^, homozygous deletion of individual heart enhancers associated with these functions did not alter cardiac gene dosage, cause cardiac malformations, or reduce embryonic viability. These findings are consistent with previous studies of mammalian TF enhancer landscapes in limb and brain development demonstrating that enhancers with partially overlapping activities often provide transcriptional robustness^52,65,121^. Our results also confirm previous predictions based on integration of transcriptomic and epigenomic data from embryonic heart tissues that projected a generally larger number of putatively redundant heart enhancers near TF genes, as compared to other tissue-specific or housekeeping genes^48,52,122^. Similar regulatory architectures have been observed for other developmental processes in which multi-partite enhancer landscapes were shown to orchestrate the dynamic spatiotemporal expression patterns of developmental regulator genes^76–78,123^. In several instances, even the combined deletion of tissue-specific enhancers with overlapping activities led to only partial or even insignificant reduction of target gene expression, demonstrating that certain loci of key developmental regulators exhibit exceptionally high regulatory resilience^52,78,124^. The enhancers involved can act additively, super-additively, cooperatively or in an interdependent manner to deploy robust compartment- or cell type-specific gene dosage^123,125–127^. Pioneering studies in *Drosophila* previously demonstrated that locus-specific enhancers with overlapping transcriptional activities, termed shadow enhancers, are critical for normal development under genetic or environmental stress, while they act largely redundant under optimal conditions^129–131^. Similarly, studies of mammalian eye lense, tooth or limb development uncovered that normally redundant enhancers confer robustness to deleterious genetic perturbations^52,132,133^. Consistent with these findings, we show that *Gata4* enhancers appearing nonessential under normal conditions become essential for embryonic survival under genetic constraint, likely related to dose-dependent cell type-specific *Gata4* functions promoting RV integrity^27^, AV cushion formation^29,62^ or conduction system related defects^134,135^. As the heart is the first organ to attain functional indispensability during embryogenesis, a substantial proportion of cardiac enhancers is thus likely to be implicated in the genetic mechanisms underlying heart defects or fetal demise, consistent with a role as cardiac disease modifiers^33,37^.

While a large number of cardiac cell populations contributing to four-chambered heart morphogenesis has been identified using single-cell transcriptome profiling^18,32,84,85^, integration with matched chromatin accessibility data provided more detailed insights into the gene regulatory programs active in both embryonic and adult hearts^49,50,136^. Here, we performed multiome joint profiling of the transcriptome and accessible chromatin in single nuclei from developing mouse hearts across midgestation, enabling direct association of gene regulatory architecture with transcriptional output, as recently applied to delineate a MEF2C-dependent, segment-specific GRN directing heart tube morphogenesis^11^. The identified clusters showed strong correspondence with the genetic signatures of cardiac cell types reported in previous single-cell studies of the developing heart at comparable stages^18,32,84,85^. The related regulatory signatures, including correlated gene-enhancer associations across multiple timepoints, enabled functional characterization of the *Hand2* cardiac enhancer landscape. Thus, our approach provides a robust framework for discovering functionally relevant heart enhancers and their cell type specificities as key nodes within cardiac cell type GRNs^7,13,137^. Analysis of cardiac TF cistromes previously identified prevalence of linked ETS1/2 and GATA4 motifs in enhancers of the EC lineage, while NKX2-5, TBX5 and MEF2C preferentially occupied CM-selective enhancers bound by GATA4^51,74^. While we observed similar signatures, the binding motifs of GATA4 and HAND2 were among the TFs showing the broadest enrichment across the identified cell type clusters, consistent with their critical functional roles in the respective cardiac cell types and lineages^29,30,59,91^. By integrating transcriptomic and chromatin signatures, we identified tens of thousands of correlative cardiac gene-enhancer connections associated with cluster-specific genes. We provide this resource as an accessible and comprehensive framework for exploring genome-wide candidate heart enhancer profiles linked to target genes across cardiac cell types, extending upon existing resources for genome-wide mammalian heart enhancer prediction^48,70,138^.

The utility of our multiome-based framework for accurate *in vivo* cardiac enhancer prediction was validated through comparison with the VISTA Enhancer resource, which includes nearly 400 enhancer elements driving reporter activity in the mouse embryonic heart^94^. We therefore anticipate that, along with other recent studies integrating multi-modal single cell profiles underlying mouse or human heart development^11,49,136^, our multiome-based map of target gene and cell type-associated heart enhancer predictions will contribute to advancing more accurate reconstruction of cardiac cell-type GRNs based on underlying enhancer modules, with the potential to unveil novel non-coding regulatory mechanisms and nucleotide variation linked to congenital heart pathologies^1,37^. However, despite recent mapping of causal atrial fibrillation variants to CM-specific *cis*-regulatory elements^49^ and the demonstrated value of multiomic studies in identifying risk variants associated, for example with neurological diseases^139,140^, our analysis of *de novo* variants identified in sizeable cohorts of individuals with CHD^39,50^ did not reveal a major specific cell type enrichment for these variants. While this observation was rather unexpected given the established lineage- and compartment-specific defects underlying several CHD manifestations, our results align in part with previous snATAC-seq findings from human hearts^50^. This pattern may therefore reflect attenuation of effects across cells computationally assigned to distinct clusters or indicate that a considerable fraction of individual de novo CHD variants exert only limited influence on cardiac cell type–specific regulatory programs.

Consistent with the existence of CTCF-dependent pre-formed chromatin topology^141,142^, our chromatin polymer modeling of C-HiC data identifies the *H2*-URI as a regulatory unit with cardiac-specific conformational properties. In heart tissue the *Hand2* TSS appears to show a more stable steady-state association with its distal anchor point (*H2*-435), which contains a strong tissue-invariant CTCF site flanking a deeply conserved OFT/RV enhancer (hE1/*H2*-437)^68^. This spatial configuration is less pronounced in limb or craniofacial tissues, despite the presence of multiple limb enhancers within the *H2*-URI^100,101,120^. Consequently, these findings indicate that the proposed chromatin domain in the developing heart fosters an environment to stabilize interactions between *Hand2* and cardiac enhancers in the *Hand2* upstream domain. Our results emphasize an essential requirement of the EC lineage enhancers contained in the *H2*-URI in directing *Hand2*-mediated transcriptional control of downstream effectors and thereby enable trabecular morphogenesis and AV cushion patterning^30,72^. Regulatory variation and loss-of-function effects in these EC enhancers might thus underly or contribute to CHD involving hypoplastic ventricles or ventricular non-compaction, tricuspid atresia or interventricular septal defects^30,72,143^. Given that HAND2 was only partially reduced in the RV and OFT, the enhancer collective controlling SHF-derived myocardial expression of *Hand2* in the RV/OFT essential for progression of heart morphogenesis beyond E9.5^25,28^ appears distributed across a larger distance, likely involving also *Hand2*-proximal and downstream elements^60,144^.

The identification of a complement of upstream *Hand2* heart enhancers is also relevant for further refinement of the mechanisms mediated by lncRNAs such as *Upperhand* (*Uph/Hand2os1*) and *Handsdown* (*Hdn*), which both through active lncRNA transcription have been implicated in either activating or suppressing cardiac *Hand2* transcription, respectively^98,99,145^. While active transcription of *Uph* has been shown to be required for RV/OFT-specific *Hand2* expression and related embryonic viability through a mechanism that permits GATA4-induced activation of the *H2*-3kb cardiac enhancer^98^, *Hdn* is required to suppress *Hand2* in the developing myocardium, thereby preventing dose-dependent RV hyperplasia in adult mice^145^. This repressive role of *Hdn* has been proposed to occur through its physical interaction with upstream *Hand2* heart enhancers^145^.

In contrast to the generally weaker, predominantly mammal-restricted, sequence conservation reported for *in vivo* heart enhancers^61^, several cardiac enhancers within the *H2*-URI locus exhibit deep sequence conservation including a subset with core regions conserved in *Coelacanth*. While this pattern points to integration into ancient cardiac GRNs that may have emerged during the fish-to-tetrapod transition^146,147^, a recent study demonstrated widespread ‘indirect conservation’ of enhancer orthologs in embryonic hearts, driven by shuffling of lineage-specific TF motifs, revealing hidden functional conservation despite sequence divergence^68^. ChromBPNet deep learning trained on our multiome dataset identified motifs of cardiac lineage TFs that align with the cardiac cell type specificities of the *Hand2* heart enhancer identified, providing a foundation for functional dissection of relevant upstream TF interactions and enhancer grammar^10,148^. Interestingly, GATA motifs were among the most highly predicted functional motifs across heart enhancers identified within the *H2*-URI, reinforcing the role of GATA4 as a pioneer TF that, in coordination with lineage-selective non-pioneer factors like ETS1, facilitates chromatin opening and enhancer priming^51,74^. Such multiome-based deep learning applications offer particular promise for refining nucleotide grammar implicated in CHD and advancing the synthetic design of cell type-specific enhancers holding significant therapeutic potential^58,149,150^.

## Methods

### Ethics statement, animal studies and experimental design

The study complies with all relevant ethical regulations and involved analysis of mouse embryos at days (E) 9.5-13.5 and inspection of animals of both sexes. All genetically modified mouse lines were housed and bred at the CAF animal facilities (Mu28/50) of the University of Bern under licenses BE96/2020 and BE90/2023, with approval by the Regional Commission on animal experimentation and the Cantonal Veterinary Office of the city of Bern (Switzerland). All founder mice from genomic deletion lines as well as mouse embryos for enSERT transgenesis at E10.5 were generated at the Centre for Transgenic Models (CTM) of the University of Basel under licenses 1023G/H. Validation of transgenic enhancer-reporter activity in E11.5 embryos was conducted at Lawrence Berkeley National Laboratory (LBNL), using mice maintained in the Animal Care Facility, which is fully accredited by AAALAC International. All animal work at LBNL was reviewed and approved by the LBNL Animal Welfare Committee under protocol numbers 290003 and 290008. Zebrafish lines were produced and held at the animal facility of the Institute of Anatomy of the University of Bern with national license number 35. Zebrafish experiments were conducted according to legal premises regulated by the Cantonal Veterinary Office of the city of Bern (Switzerland).

All mice were maintained with water supply on a 12:12 light-dark cycle, with relative humidity set at 30–70% and a temperature of 22 °C + /– 2 °C (Bern/CTM) or 20–26.2 °C (LBNL). Mice at the University of Bern were housed in standard IVC cages GM500 Tecniplast, on Safe® Aspen wood granulate bedding with enrichment consisting of Pura® crinkle brown kraft paper nesting material, Pura® Brick Aspen Chew Block, and red polycarbonate mouse house and fed on Kliba Nafag standard breeding (3800) and maintenance diet (3430). Mice at the CTM (University of Basel) were housed in standard IVC cages GM500 Tecniplast, on ABEDD classic 4604 bedding with enrichment consisting of tissues and a red polycarbonate mouse house or a transparent tunnel, that was used for tunnel handling as well. Mice were fed on Kliba Nafag irradiated feed (3432 extrudate) and received acidified water with pH 2.8. Mice at LBNL were housed in standard micro-isolator cages on hard-wood bedding with enrichment consisting of crinkle-cut naturalistic paper strands and fed on ad libitum PicoLab Rodent Diet 20 (5053).

Embryos of zebrafish (Danio rerio) were obtained from adult fish aged 3-12 months, raised at maximal 5 fish/l. Fish were maintained under the same environmental conditions: 27.5-28°C, with 14 hours of light and 10 hours of dark, 650-700μs/cm, pH 7.5 and 10% of water exchange daily. Embryos at 2-5 dpf were obtained for experiments from family crosses of 2-3 females and 4-5 males. After collection embryos were raised in petri dishes at a density of 50 embryos per plate in 50 ml of E3 embryo medium, and kept in an incubator at 28 °C.

All animals were health checked and monitored daily for food and water intake by trained personnel. Euthanasia of mice was performed in the home cage using CO2 asphyxiation while ensuring gradual fill and displacement rate. Selection of sample size and randomization was performed as follows:

#### Transgenic reporter assays in mouse embryos

The sample size selection and scoring criteria were informed by prior experience conducting transgenic mouse assays for over 3,000 candidate enhancers, as documented in the VISTA Enhancer Browser (http://enhancer.lbl.gov). Mouse embryos were excluded from analysis if they lacked the reporter transgene or corresponded to an incorrect developmental stage. For site-directed enhancer-reporter assays (enSERT), transgenic outcomes were validated in at least two independent biological replicates, following the established standards of the VISTA Enhancer Browser^94,151^.

#### Genomic knockout embryos

Sample sizes for experiments based on genomic enhancer/gene deletions in mouse embryos were determined empirically, following the strategy described in our previous studies^52,53^. The minimal number of biological replicates analyzed for each experiment is specified in the corresponding experimental sections below. All phenotypic analyses of knockout (KO) mice were conducted using a matched littermate design. *H2*-URI^Δ/Δ^ and enhancer KO embryos and mice analyzed in this study were generated by crossing heterozygous animals for the respective deletions, enabling direct comparison of littermates with different genotypes. Tissue collection and processing were performed blinded to genotype whenever applicable.

### CRISPR/Cas9 deletion mouse lines

Enhancer KO and *H2*-URI^Δ^ mouse strains were engineered at the Centre for Transgenic Models (CTM) of the University of Basel using CRISPR-EZ based on electroporation of CRISPR/Cas9 reagents into fertilized mouse eggs^73^. Single-guide RNAs (sgRNAs) targeting sequences flanking the genomic regions to be deleted were designed using CRISPOR^152^ (http://crispor.tefor.net/). To induce deletions, HiFi Cas9 Nuclease (IDT, #1081061) was complexed with cr:tracrRNA (IDT) at equimolar concentrations (final concentration of each component: 8 μM) in Hepes-KCl buffer, following previously described procedures^53^. FVB/NRj (Janvier Labs) strain mouse oocytes originating from the oviducts of super-ovulated FVB/NRj females (8 weeks) mated to FVB/NRj males (8 weeks or older) were incubated in KSOM (Sigma/Merck MR-106) supplemented with amino acids at 37 °C, 5% CO2 and then electroporated with the RNP mix. Following repeated culture in KSOM medium, embryos were transferred into the oviduct of pseudo-pregnant Swiss Albino (Janvier Labs; strain name: RjOrl:SWISS) females on the same day. Genotyping of mouse alleles was carried out by PCR using either High-Fidelity Platinum Taq Polymerase (Fisher Scientific) or standard Taq polymerase. For F0 and F1 mice, deletion breakpoints generated by non-homologous end joining (NHEJ) were identified through Sanger sequencing of the corresponding PCR-amplified fragments (**Figs. S2, S4** and **S11**). Genomic deletion coordinates and sgRNA sequences used for genome editing are listed in **Table S2**. Primer pairs specific for the deleted genomic interval or WT allele were used for PCR-genotyping of mice and embryos (**Table S3**).

### Site-directed enhancer-reporter transgenesis

Predicted enhancer elements were PCR-amplified from mouse genomic DNA and cloned into the PCR4-Shh::lacZ-H11 vector (Addgene #139098) for site-directed integration at the safe-harbor locus H11 (enSERT)^69^ using Gibson assembly. Genomic elements were amplified with the proofreading Platinum™ SuperFi II DNA Polymerase (Invitrogen). Sanger sequencing was used to corroborate sequence integrity of cloned constructs. EnSERT transgenic enhancer-reporter embryos at E10.5 were generated at the Centre for Transgenic Models (CTM) at the University of Basel following a modified published protocol^69,151^. Briefly, the enSERT procedure was carried out by microinjecting a mixture containing CRISPR cr:trRNA (20 ng/μl), sgRNAs generated using the CRISPOR tool^152^, HiFi Cas9 Nuclease (IDT #1081061; 200 ng/μl), and enSERT donor vector harboring the enhancer element of interest (400 ng/μl) in microinjection buffer (1 mM Tris, pH 8.0; 0.1 mM EDTA) into the pronuclei of single-cell stage fertilized B6CBAF1/JRj hybrid embryos. Following culture in KSOM medium supplemented with amino acids, embryos were transferred into the oviduct of pseudo-pregnant Swiss Albino females on the same day and subsequently collected and stained with X-gal following standard techniques^151^, as described below. EnSERT transgenic enhancer-reporter embryos at E11.5 were generated at LBNL following the published approach^69,151^. Briefly, single-guide RNAs (sgRNAs; 50 ng/μl) targeting the H11 locus and Cas9 protein (Integrated DNA Technologies, catalog no. 1081058; final concentration: 20 ng/μl) were prepared in microinjection buffer (10 mM Tris, pH 7.5; 0.1 mM EDTA). The mixture was injected into the pronuclei of fertilized one-cell stage FVB/NJ embryos (Jackson Laboratory; strain no. 001800) collected from oviducts of super-ovulated 7–8-week-old FVB/NJ females mated with 7–8-week-old FVB/NJ males. Injected embryos were cultured in M16 medium supplemented with amino acids at 37 °C in 5% CO₂ for approximately 2 hours and subsequently transferred into the uteri of pseudo-pregnant CD-1 surrogate females (Charles River Laboratories; strain code: 022).

### Analysis of LacZ reporter activity

To visualize LacZ reporter activity, transgenic embryos generated by enSERT were collected at E10.5 or E11.5 and fixed in 4% paraformaldehyde (PFA) for 15 or 20 minutes, respectively. Fixed embryos were incubated overnight at 4 °C in X-gal staining solution with gentle rolling. Following staining, embryos were genotyped to confirm integration of the transgenic construct. All transgenic results were validated in at least two independent biological replicates, following enSERT criteria consistent with those applied in the VISTA Enhancer Browser^94^. Embryos positive for the transgene were imaged using a Leica S9i microscope (E10.5) or a Leica MZ16 microscope (E11.5). For each positive embryo, close-up images of the heart were captured, and staining patterns were categorized based on X-gal signal distribution across major anatomical regions. For section analysis, selected embryos were embedded in gelatin/sucrose according to standard protocols. Sagittal sections (10 μm thick) were prepared and counterstained with eosin to visualize embryonic structures under bright-field microscopy. Section images were acquired using a Nikon Eclipse Ti-E microscope.

### Generation of transgenic zebrafish lines

An established *Tol2*-based fluorescent reporter vector framework (pCK036) was used to assess enhancer activity in zebrafish embryos^80^. This system incorporates a dual-reporter cassette comprising the transgenic mouse *β-globin* minimal promoter (mbG) driving *mCerulean* followed by an SV40 polyadenylation signal, and the *α-crystallin* promoter (cryaa) driving a Venus cassette including a bovine growth hormone polyadenylation (bghpA) signal. Cryaa:*Venus* expression provides constitutive fluorescence in the eye lens, serving as a transgenic control while allowing simultaneous detection of enhancer-driven *mCerulean* activity^80^. A vector version containing a desmin regulatory element upstream of the *mCerulean* unit (desma-mbG:*mCerulean*-SV40pA, cryaa:*Venus*-bghpA)^80^ was used as a template for EcoRI digestion-mediated replacement of the desmin fragment with individual enhancer elements (*G4*-124, *G4*-93, *G4*+177, or *G4*+182) (**Fig. 1**). These enhancer sequences were PCR-amplified from mouse genomic DNA. The resulting pCK036 enhancer-reporter constructs were co-injected with *Tol2* mRNA into one-cell stage zebrafish embryos to enable genomic integration.

### Quantitative real-time PCR (qPCR)

Mouse embryonic hearts were micro-dissected at E9.5, E10.5 or E11.5 in ice-cold PBS, transferred to RNA-later (Sigma-Aldrich), and stored at -20 °C until further processing. Total RNA was extracted from micro-dissected tissues using the ReliaPrep RNA Tissue Miniprep Kit (Promega) according to the manufacturer’s instructions. Complementary DNA (cDNA) was synthesized using the GoScript Reverse Transcription System (Promega) with poly-dT primers, following the manufacturer’s protocol. QPCR was performed on a ViiA 7 Real-Time PCR System using PowerTrack SYBR Green Master Mix (Applied Biosystems). Relative gene expression levels were determined using the 2^-ΔΔC^ method, normalized to the *Rpl19* housekeeping gene, with the mean value of wild-type (WT) control samples set to 1, as described previously^53^. For each genotype, hearts from at least five embryos (biological replicates) were analyzed. Primer sequences are listed in **Table S4**.

### Whole-mount *in situ* hybridization (ISH)

Whole-mount in situ hybridization (ISH) using digoxigenin-labeled antisense riboprobes was performed as previously applied^153^. Embryos were fixed overnight at 4 °C in 4% paraformaldehyde (PFA) prepared in PBS, gradually dehydrated through a 25%, 50%, and 75% methanol/PBT series, and stored in 100% methanol at -20 °C until further use. For ISH, tissues were rehydrated through a reverse methanol/PBT series, bleached in 6% hydrogen peroxide (in PBT) for 15 minutes, and permeabilized with 10 µg/ml proteinase K. Protease activity was subsequently quenched with 2 mg/ml freshly prepared glycine in PBT for 5 minutes, after which embryos were post-fixed in 0.2% glutaraldehyde and 4% PFA in PBT for 20 minutes. Embryos were incubated in pre-hybridization buffer (50% deionized formamide, 5x SSC pH 4.5, 2% Roche Blocking Reagent, 0.1% Tween-20, 0.5% CHAPS, 50 mg/mL yeast RNA, 5 mM EDTA, 50 mg/mL heparin) at 65 °C for ≥3 h, followed by overnight hybridization in 1 mL solution containing 1 µg/mL DIG-labeled *Hand2* or *Gata4* riboprobe at 70 °C. On the following day, embryos were washed extensively and digested with 20 µg/mL RNase at 37 °C for 45 min. After additional washes and pre-blocking, samples were incubated overnight at 4 °C with anti-digoxigenin antibody (1:5000; Roche, cat. no. 11093274910). Excess antibody was removed by serial washes. Signal detection was performed after equilibration in NTMT using BM Purple (Roche, cat. no. 11442074001) and terminated prior to saturation by repeated PBT washes. Whole-mount ISH was assessed qualitatively to evaluate spatial expression patterns. For each genotype, ≥3 embryos were analyzed. Images were acquired using a Leica S9i stereomicroscope equipped with a Leica digital camera. Brightness and contrast were adjusted uniformly using Photoshop (CS5).

### Live-fluorescence analysis in zebrafish

Imaging of whole-mount live zebrafish embryos was conducted with a fluorescent stereomicroscope (Nikon SMZ25) coupled to a DS-Fi3 camera. Briefly, embryos were anesthetized with 0.016% Tricaine in E3 media and kept in a 100mm Petri dish. For image acquisition, larvae were mounted in 1% low melting agarose in a MatTek petri dish. Images were acquired with a 20x water immersion objective and z-stack size was 1.08 µm. Images were then processed with FiJI software. 3D projection images were first treated with a mean filter and a radius of 1.0 pixels. Interpolation was also applied when rendering the 3D projections.

### Immunofluorescence (IF)

Immunofluorescence on mouse embryonic tissues was carried out as previously described^53^. Mouse embryos were isolated in cold PBS, fixed in 4% paraformaldehyde (2-3 h), cryoprotected in sucrose, and embedded in a 1:1 mixture of 30% sucrose and OCT, followed by storage at -80 °C. Cryosections (10 µm) were blocked with BSA and incubated overnight with primary antibodies against HAND2 (R&D, AF3876-SP, 1:500), GATA4 (Santa Cruz, sc-25310, 1:300), or GFP (Invitrogen, A10262, 1:300). Secondary antibodies included donkey anti-mouse Alexa Fluor 647, donkey anti-goat Alexa Fluor 488, or goat anti-chicken Alexa Fluor 488 (all Invitrogen, 1:1000). Anti-SMA-Cy3 (1:250, Merck/Sigma, C6198) was applied additionally when indicated. Nuclei were counterstained with Hoechst 33258. TUNEL staining was performed using the In Situ Cell Death Detection Kit (Roche, Cat. No. 11684795910). Cryosections were treated with MeOH/glycine, permeabilized with Triton X-100/sodium citrate, incubated with TUNEL reagent (1 h, 37 °C) and mounted in DAPI-containing medium for imaging. Whole-mount IF in zebrafish embryos was performed as follows: embryos were fixed overnight at 4 °C in 4% paraformaldehyde (PFA; EMS, 15710), washed in 0.1% PBS-Tween20, and permeabilized for 60 min in 0.5% PBS-TritonX100. After blocking for 2 h in *histoblock* solution (5% BSA, 5% goat serum, 20 mM MgCl₂ in PBS), embryos were incubated overnight at 4 °C with primary antibodies against eGFP (Aves, eGFP-1010; 1:300) and dsRed (Takara, 632496; 1:200) diluted in 5% BSA. The following day, embryos were thoroughly washed with 0.1% PBS-Tween20 at least 6 times. Embryos were incubated overnight at 4 °C with Alexa Fluor 488/568 secondary antibodies (Life Technologies, 1:250) and DAPI (Merck, 1246530100; 1:1000) in 5% BSA. Specimens were washed with 0.1% PBS-Tween20 several times, mounted in 1% low-melting agarose. All fluorescent section images were acquired using a Leica TCS SP8 DLS fluorescence microscope. Brightness and contrast were adjusted uniformly in Photoshop (CS5) following image acquisition.

### Nuclei extraction for single-cell multiome analysis

All procedures were conducted on ice unless otherwise indicated. Hearts, with the pericardium removed, were dissected from wild-type mouse embryos (CD-1 strain) at embryonic days E9.5 (n=23), E10.5 (n=8), and E11.5 (n=5), then pooled in cold PBS. To ensure comprehensive capture of cardiac structures and the aortic sac, the dissection included tissues adjacent to both the arterial and venous poles, such as pharyngeal arch (PA) tissue and pulmonary endoderm (PE). Collected hearts were washed three times with cold PBS (w/o Ca/Mg) before tissue dissociation in 1 ml PBS supplemented with 0.4% BSA (Invitrogen, AM2618) using a glass douncer (pestle A, 6–10 strokes). The resulting tissue suspension was transferred to a 1.5 ml Eppendorf tube and centrifuged at 300 g for 3.5 minutes at 4°C. Following removal of the supernatant, the cell pellet was gently resuspended in lysis buffer (10 mM Tris-HCl pH 7.4, 10 mM NaCl, 3 mM MgCl₂, 1% BSA [Invitrogen AM2618], 0.1% Tween-20, 0.1% IGEPAL, 0.01% Digitonin, 1 mM DTT, and 1 U/µl RNase inhibitor [Roche]) by pipetting up and down 10–15 times with wide-bore tips. Samples from E9.5 embryos were incubated in lysis buffer for 1 minute, whereas E10.5/E11.5 samples were incubated for 1.5 minutes. Lysis was stopped by adding wash buffer containing 10 mM Tris-HCl pH 7.4, 10 mM NaCl, 1% BSA (Invitrogen AM2618), 3 mM MgCl₂, 0.1% Tween-20, 1 mM DTT, and 1 U/µl RNase inhibitor (Roche). The nuclei suspension was first passed through a pre-wetted 70 µm cell strainer and centrifuged at 300 g for 5 minutes at 4°C. After discarding the supernatant, the nuclei pellet was resuspended in wash buffer and filtered through a pre-wetted 45 µm strainer. The suspension was then centrifuged again under the same conditions, the supernatant removed, and the pellet resuspended in 80 µl (E9.5) or 200 µl (E10.5 and E11.5) of nuclei dilution buffer (10x Genomics). A final filtration step was performed using a pre-wetted 45 µm strainer. Nuclei number and quality were evaluated using a cell counter and DAPI staining, with visual inspection under a microscope confirming nuclei integrity and the absence of debris or aggregates.

### Multiome RNA-seq and ATAC-seq processing

The Chromium Next GEM Single Cell Multiome ATAC + Gene Expression Reagent Bundle (10x Genomics, PN-1000285) was used with a targeted recovery of 10’000 nuclei per sample. Nuclei processing was performed as described in the user manual. Briefly, nuclei were incubated in a transposition mix for 1 hour at 37 °C, after which the transposed nuclei were loaded into the Chromium Controller for GEM generation, followed by barcoding through incubation at 37 °C for 45 minutes and at 25 °C for 30 minutes. After addition of Quenching Agent to stop the reaction, samples were cleaned using MyONE SILANE and SPRIselect Dynabeads, after which the library was pre-amplified by PCR and re-purified with SPRIselect beads. The sample was subsequently divided, with 40 µL allocated for ATAC library preparation and 35 µL used for cDNA amplification. For ATAC-seq, P7 and a sample index were added to pre-amplified transposed DNA during ATAC library construction via PCR. The final ATAC libraries contain the P5 and P7 sequences used in Illumina bridge amplification. The cDNA library was amplified by PCR and enzymatically fragmented before the sequencing adaptors were added via end repair, A-tailing, adaptor ligation, and PCR. Both types of libraries were sequenced on a DNBSEQ-G400 system (BGI), using paired-end (PE) 100/100/10/10 (cDNA) or PE 50/50/8/16 (ATAC) settings. Sequencing data were processed using the Cell Ranger Arc workflow (version 2.0.0). A total of 7,198 cells were obtained at E9.5 (cDNA: 43,676 mean raw reads per cell; ATAC: 57,963 mean raw reads per cell), 3,760 cells at E10.5 (cDNA: 87,013 mean raw reads per cell; ATAC: 108,771 mean raw reads per cell), and 5,614 cells at E11.5 (cDNA: 55,344 mean raw reads per cell; ATAC: 73,931 mean raw reads per cell). In total, 16,572 cells were recovered, with a median of 2,882 genes detected per cell.

### Multiome RNA-seq and ATAC-seq Library Preprocessing

RNA-seq count matrices were preprocessed independently for each time point using Seurat v4.2.1^154^. Count matrices were initially filtered to exclude cells based on the following thresholds: Library size (nCounts_RNA) > 25,000 and < 500, percent of mitochondrial reads (percent.mt) < 20%, number of genes detected per cell (nFeature_RNA) < 200 and > 7500. Prior to dimensionality reduction, data were normalized and scaled using the SCTransform function, and 3,000 variable genes were identified. Principal component analysis (PCA) with 50 components was performed using variable genes, followed by UMAP dimensionality reduction based on all 50 principal components. Each dataset was visualized on UMAP space and cells were colored by average hemoglobin gene expression. Distinct clusters of cells with high hemoglobin expression were excluded. Subsequently, individual datasets were merged and the merged RNAseq object was preprocessed as described above. ATAC-seq count matrices were processed using the ArchR package from R (version 1.0.2)^155^. Count matrices from all three time points were concatenated and analyzed jointly. Cell filtering followed the default ArchR criteria, retaining only cells with a transcription start site (TSS) enrichment score greater than 4 and more than 1,000 unique nuclear fragments. Doublets were detected using the function *addDoubletScores* and removed. ATAC-seq peaks were called using MACS3 [https://github.com/macs3-project/MACS] settings *shift* to -100 and *extsize* argument to 200. Peaks were filtered to exclude elements overlapping with ENCODE mm10 blacklisted regions [https://github.com/Boyle-Lab/Blacklist/tree/master/lists]. MACS3 peaks were added to the ATAC-seq object using the *CreateChromatinAssay* function from the Signac package v1.9.0^156^ and *addPeakSet* within ArchR. Prior to dimensionality reduction, top variable features were identified using the *FindTopFeatures* function setting the *min.cutoff* argument to “q0”. Subsequently, TF-IDF normalization and singular value decomposition was applied to the MACS peak matrix. Dimensionality reduction was performed using Signac’s LSI method using the function *RunUMAP* setting the *reduction* argument to “lsi” and considering all 50 dimensions except the first.

### Multiome data integration and clustering

RNA-seq and ATAC-seq matrices were filtered to include cells that passed filtering criteria from Seurat and ArchR pre-processing steps. Multimodal neighbors were identified using the *FindMultiModalNeighbors* function from Seurat using the PCA and LSI reductions from SCT RNA matrix and ATACseq matrix, respectively. Clustering was performed using the Seurat *FindClusters* function at different resolutions (0.05, 0.4, 0.75, 1.2) with 0.75 selected as the optimal clustering resolution resulting in 28 clusters. Differentially expressed genes and differentially accessible peaks for each cluster were identified using the Seurat *FindAllMarkers* function setting the *min.pct* to 0.25 and *logfc.threshold* to 0.58 for genes and 1 for peaks. Clusters were labeled after manual curation based on their gene expression signature. Motif analysis in the MACS3-defined peak regions was performed in ArchR using the EncodeTFBS motif database. Motif enrichment was performed using the *peakAnnoEnrichment* function setting FDR ≤ 0.01 and Log2FC ≥ 0.58 cutoffs. Motif enrichment results were then filtered based on the mRNA expression of the cognate TF-gene, in published bulk-RNA-seq profiles of whole hearts (ENCODE; E10.5-E12.5; identifiers: ENCFF770SOB, ENCFF159DWP, ENCFF484QWQ)^70^. Motifs whose cognate TF-gene showed an expression equal or higher than 1 FPKM in at least one of the considered stages were retained. Considering only the twenty cardiac cell clusters, Spearman’s correlation distance was then calculated across all the remaining motifs, and highly correlated features were removed using the *findCorrelation* function of the package caret (v6.0.91). The resulting matrix was log-transformed and used as input for PCA (using the *prcomp* function in R). Elbow plot analysis revealed four PCs as the optimal number for downstream analyses, including further non-linear dimensionality reduction (to two components) via UMAP (performed via the *umap* function in the umap package in R, v0.2.8.0). As a rough proxy of pioneering activity, pervasiveness scores were calculated based on the relative strength of the enrichment for each TF motif, separately in each cell cluster. First, the values from the matrix obtained above were ranked separately for each cell cluster (using the *rank* function in R). After that, these ranks were subtracted one, and a score was calculated for each TF motif, aggregating the ranks across the cell clusters (Borda Count). A relative sum of these scores was also obtained by dividing these numbers by the best possible value (the maximum).

### Multiome chromatin accessibility of *in vivo* validated enhancers

The *getGroupBW* function from ArchR^155^ (v1.0.2) was used to compute genome-wide chromatin accessibility tracks per lineage. Human and mouse *in vivo* enhancer elements (3’869 elements in total) were downloaded from the VISTA enhancer browser^94^ (March 2024). The elements were grouped by their tissue activity using activity annotations provided by the VISTA enhancer browser. The tool *bigWigAverageOverBed* (from UCSC tools) was used to compute the average chromatin accessibility of VISTA elements per lineage.

### Differentially accessible peak fraction per multiome cluster

Cluster-specific peaks and cluster-specific differentially accessible peaks (1 vs. all other clusters) were computed (**Table S7**). Peak calling was performed using MACS3 (v3.0.2) with the callpeak function on paired-end aligned BAM files (-f BAMPE), previously filtered using a mapping quality threshold of MAPQ ≥10. The mouse genome size was specified (-g mm), and to avoid empirical model building, the --nomodel option was applied. Fragment size was manually defined using a shift of -100 bp and an extension size of 200 bp (--shift -100 --extsize 200), enabling accurate representation of the fragment center for peak detection. Differential enrichment (DE) peak calling analysis was performed by applying MACS3 (v3.0.2) *callpeak* function, using parameters (-f BAMPE, -g mm, --nomodel, --shift -100, --extsize 200)^157^. To identify regions showing differential enrichment, each BAM file was designated as the treatment (-t) sample, while the remaining BAM files were used as controls (-c). Peaks were classified based on their chromatin state (enhancer, promoter, CTCF, other) with ChromHMM^158^. Publicly available H3K4me3, H3K4me1, H3K27ac, H3K27me3, H3K36me3, H3K9me3 (E11.5) and CTCF (postnatal) mouse heart datasets^70^ were used to establish a 12-state ChromHMM model using the LearnModel command^97^.

### Overlap of cluster-specific peaks with GWAS and CHD variants

GWAS summary statistics for multiple traits were obtained from the GWAS Catalog^159^ as of May 2024. *De novo* non-coding variants previously identified in patients with CHD and healthy controls were collected from Ameen et al.^50^ and Xiao et al.^39^. Variants were filtered to include only single-nucleotide substitutions with experimentally annotated functional effects, classified as increasing, decreasing, or showing no change in gene activity. To evaluate potential regulatory relevance, overlaps between cluster-specific chromatin peaks and each variant set were computed using *bedtools intersect* (v2.31.1)^160^. The number of overlapping regions was normalized by both the total number of variants and the total base pair length of the chromatin peaks.

### Computation of multiome gene-enhancer links

Gene-enhancer associations were calculated using the Signac package (v1.10.0) in R. The gene set for analysis was assembled by integrating three sources: (i) curated candidate genes, (ii) marker genes identified from cell type/cluster annotations, and (iii) the top 1,000 genes with the highest expression levels across all cells. Peak-level statistics were first computed using the *RegionStats* function, followed by gene-enhancer link inference with the *LinkPeaks* function. Links were computed within a ±2.5 Mb window around each gene’s transcription start site (TSS). Links were inferred separately for all cells or cells from the 20 cardiac cell clusters.

### ChromBPNet deep learning variant and motif predictions

ChromBPNet (v1.1.0) was used to train sequence-to-accessibility models for nine cardiac cell type clusters aggregated based on multiome profiling (CM, aSHF/OFT, pSHF, CPM, EC, VEC, EndMT, EP, and cpNCC; **Fig. 3**). Cluster-specific fragment subsets were generated by filtering with respective cell barcodes. Preprocessing steps were performed using the default pipeline of ChromBPNet, which included peak-calling with MACS2, filtering of ENCODE blacklist regions, and generation of negative (non-peak) background regions. Train/validation/test chromosome splits were generated with ChromBPNet prep splits, designating chr1/chr3/chr6 as test, chr8/chr18 as validation, and the remaining chromosomes for training. A pre-trained bias model from the authors was used, which was trained on an ATAC-seq dataset from the K562 human cell line (ENCSR868FGK). After ChromBPNet model training, the model quality was checked by ensuring that the bias model and ChromBPNet model performance metrics reached the recommended thresholds. The trained models were further applied for the generation of contribution score tracks and for the prediction of variant effects on chromatin accessibility using the variant-scorer package.

### Web-interface for interrogation of multiome data (R ShinyApp)

An interactive web interface (https://barozzilab.shinyapps.io/locus_heatmap/) was developed to provide genome-wide predictions of cardiac compartment or cell type-specific regulatory elements derived from our multiome data. For each chromatin accessibility peak, average accessibility per cluster was calculated using the *getGroupBW* and *bigWigAverageOverBed* methods. The interface allows users to visualize either raw (unnormalized) accessibility values or Z-score–normalized accessibility across peaks. ChromHMM chromatin state annotations^158^ and multiome-based predictions of gene-enhancer interactions for cardiac-expressed genes (+/-2.5 kb around the TSS) were generated as described above. Experimentally validated enhancer elements were retrieved from the VISTA Enhancer Browser^94^. Regions with significant CTCF binding at P0 and 8 weeks^70^, as well as binding of selected cardiac TFs in mouse embryonic hearts^72,74,75^, were annotated. Sequence conservation was assessed using phastCons60way scores obtained from the UCSC Genome Browser. Promoter annotations were downloaded from EPDnew (https://epd.expasy.org/epd/mouse/mouse_database.php?db=mouse). Distances between ATAC peaks and CTCF or promoter sites were calculated using *bedtools closest*. Overlaps between ATAC peaks and VISTA enhancer elements were identified with *bedtools intersect*.

### ATAC and ChIP-seq data processing

Previously published ATAC-seq and ChIP-seq datasets used in this study (**Table S13**) were processed using Trim Galore (v0.6.6), which integrates Cutadapt for adaptor removal. Sequencing adaptors were trimmed with default parameters (‘-j 1 -e 0.1 -q 20 -O 1’ for single-end reads and ‘--paired -j 1 -e 0.1 -q 20 -O 1’ for paired-end reads) and reads shorter than 20 bp were discarded. For read alignment, Bowtie2^161^ (v2.4.2) was used to map sequencing reads to the GRCm38/mm10 reference genome utilizing pre-built Bowtie2 indexes from Illumina iGenomes. ATAC-seq reads were aligned using the parameters ‘-q --no-unal -p 8 -X2000’, while ChIP-seq reads were aligned with ‘-q --no-unal -p 2’. Low-quality alignments (MAPQ=255) and duplicate reads were removed with SAMtools^162^ (v1.12) using the ‘markdup -r’ command and filtering options ‘-bh -q10’. Peak calling was performed using MACS2 (v2.1.0) with p-value < 0.01 and parameters ‘-t -n -f BAM -g mm --nolambda --nomodel --shift 50 --extsize 100’ for single-end and ‘-t -n -f BAMPE -g mm --nolambda --nomodel --shift 50 --extsize 100’ for ATAC data, while ChIP-seq peak calling used ‘-t -c -n -f BAM -g mm’ parameters^157,163^. Genomic profiles were visualized using the UCSC Genome Browser (https://genome.ucsc.edu).

### Region Capture Hi-C (C-HiC)

C-HiC probes were generated using the GOPHER Java desktop application (v0.5.7) with the “tiling design” option to span the *Hand2* genomic locus and flanking TADs (mm10, chr8:55524922-59121583). Custom-designed probes (SureSelectXT Custom 0.5–2.9 Mb library; Agilent Technologies) were used for target enrichment. Forelimbs (FL) and mandibular processes (MD) were micro-dissected from n=10 wildtype (WT) FVB/NRj strain embryos at E11.5, while hearts (HT) were collected from n=20 WT FVB/NRj embryos at E11.5. All dissections were performed in cold 1x PBS to preserve tissue integrity. Samples were homogenized with a Dounce tissue grinder, and the resulting cell suspensions were resuspended in 10% FCS in PBS. Formaldehyde (37%) was then added to each preparation to achieve a final concentration of 2% in a total volume of 1 mL, and cells were fixed for 10 minutes following a previously described approach^53,105^. The fixation process was quenched with 1.25M Glycine and pellets were snap-frozen in liquid nitrogen and stored at 80°C. Pellets were resuspended in fresh lysis buffer (10mM Tris, pH7.5, 10mM NaCl, 5mM MgCl2, 0.1 mM EGTA complemented with Protease Inhibitor) for nuclei isolation. Samples were washed with 1xPBS after 10 min incubation on ice and frozen in liquid nitrogen. 3C libraries were generated from thawed nuclei digested with DpnII (NEB, R0543M), followed by religation with T4 DNA ligase (Fisher Scientific) and subsequent de-crosslinking, as described previously^53,105^. A total of 500 ng of the 3C library sample, along with digested and undigested control samples, was evaluated by agarose gel electrophoresis (1.5% gel) to assess 3C library quality. Re-ligated DNA products were subsequently sheared using a Covaris ultrasonicator under the following settings: duty cycle, 10%; intensity, 5; cycles per burst, 200; duration, two cycles of 60 seconds each. After adaptor ligation and amplification of sheared DNA fragments, libraries were hybridized to the custom-designed SureSelect beads and indexed according to Agilent’s instructions at the iGE3 sequencing facility (University of Geneva). Multiplexed libraries were sequenced using 50 bp paired-end reads on a HiSeq 4000 platform at the iGE3 facility.

### C-HiC processing, normalized virtual-4C and TAD boundary calling

C-HiC data were processed following a previously described workflow^164^ using the HiCUP pipeline v0.8.1^165^. Sequenced reads were aligned to the GRCm38/mm10 reference genome with Bowtie2 v2.4.5^161^. Subsequent processing with HiCUP included filtering with parameters set to *no size selection*, *Nofill: 1*, and *format*: *Sanger*, followed by duplicate removal to retain valid Hi-C di-tag pairs (*.bam* files). Juicer command line tools v1.9.9^166^ were used for the generation of binned contact maps in .hic format from unique read pairs (MAPQ ≥ 30). To identify reciprocal interactions of *Hand2* and target enhancers, normalized virtual-4C (nv4C) profiles were generated from *.hic* output at 10kb resolution using designated genomic viewpoints (**Table S14**). In brief, for contact frequencies associated with the viewpoint (corresponding to the respective row and column of the contact matrix), the average contact frequency at the same genomic distance was subtracted and subsequently normalized by dividing by that average value. The nv4C computation was performed in Python using the standard *NumPy* library, following previously established procedures^113^. TAD boundaries in HT, FL, and MD datasets were identified using the insulation score method implemented in the *cooltools* software package^167^. In brief, insulation profiles for each tissue were calculated independently from the corresponding balanced C-HiC contact matrices at 10 kb resolution, using a 100 kb genomic window. Domain boundaries were identified as local minima of the log2-transformed insulation score signal based on the default *cooltools* parameters^167^.

### Strings and Binders (SBS) polymer model

In the SBS model^108^, chromatin is modeled as a self-avoiding polymer chain composed of beads, which contain specific binding sites for diffusing molecular binders. These binders facilitate interactions by bridging their corresponding binding sites along the polymer, thereby driving chromatin folding through thermodynamically governed phase transition mechanisms^110,168^. A genomic region encompassing the *Hand2* TAD (mm10, chr8:56281203-58021203) was used for modeling of the *Hand2*-related 3D chromatin organization in heart, forelimb and mandible tissues. The PRISMR method^113^, based on a standard Monte Carlo recursive optimization procedure, was used to derive tissue-specific SBS models^112^ of the *Hand2* locus by identifying the minimal number and optimal arrangement of distinct binding site types (visually represented by different colors) along the polymer chain; thereby revealing the binding sites that best reproduce the tissue-specific C-HiC contact matrices^112,113^. Polymer models comprising nine distinct binding site types for each tissue were identified from Knight-Ruiz (KR) normalized^169^ C-HiC datasets at 10kb resolution.

### Computation of spatial distances and contacts in the SBS polymer model

Distributions of pairwise spatial distances between the *Hand2* gene and its most prominent nv4C interaction sites in HT (*H2*-437) and FL/MD tissues (*H2*-685) were computed. Built-in functions from the Python *SciPy* library (version 1.11) were employed to efficiently compute the distributions of Euclidean distances. Histograms were generated using the open-source Matplotlib library (version 3.7). Average pairwise contact matrices for the *Hand2* locus across the three tissues were calculated following a standard approach, defining two polymer sites as being in contact if their relative spatial distance was below a specified contact threshold (μ)^109,113,170^. Molecular Dynamics simulations were based on μ=5σ^110^ and verified similar results were obtained using values in between a range of μ=3 σ and μ=10σ. Significant Pearson correlation coefficients (r) between model contact matrices and corresponding C-HiC datasets (r=0.85, r=0.90 and r=0.87 for heart, FL and MD, respectively) and distance-corrected Pearson correlation coefficients (r’)^112^ (r’=0.52, r’=0.59 and r’=0.58 for HT, FL and MD, respectively) were identified in models.

### Molecular Dynamics simulations

An advanced Molecular Dynamics (MD) simulation framework was employed to model polymers and their associated binders, both represented as beads undergoing Brownian motion and described by stochastic Langevin dynamics^109,110,168^. The equations of motion were numerically integrated using the standard Velocity-Verlet algorithm implemented in the open-source *LAMMPS* package^171^ optimized for parallel computing^172^. In the initial phase of each simulation, the polymer was modeled as a self-avoiding random chain influenced only by thermal fluctuations, while binders diffused freely within a simulation box with periodic boundary conditions. By increasing either the binder concentration or their interaction affinity, the system exhibited a micro-phase separation transition, from the initially randomly folded polymer to an equilibrium configuration in which DNA globules self-assemble along the chain corresponding to local domains of enhanced self-interaction, identified as TADs and sub-TADs in the simulated contact maps^109,110,168^. The number of different types and genomic arrangement of SBS binding sites on the chain was inferred by the PRISMR method (detailed above). The polymer model employed standard interaction potentials^173^: (i) consecutive beads were linked by a finitely extensible non-linear elastic (FENE) potential with standard parameters R0=1.6σ and K= 30kBT/σ^2^; (ii) excluded volume effects were modeled using a purely repulsive Weeks-Chandler-Andersen potential; and (iii) short-range Lennard-Jones attractions captured attractive homotypic interactions between binding sites and cognate bridging binders with previously applied bead-binder energy affinities^110,174^. Simulations with varied energies and concentrations (up to 10KBT) produced comparable outcomes, consistent with Statistical Mechanics expectations^175^. To generate a statistically robust ensemble of equilibrium single-molecule 3D polymer conformations, MD simulations were performed for up to 10^8^ MD time iteration steps, ensuring the system had reached full stationarity^109,110^. For each analyzed tissue, up to 10^3^ independent *Hand2* model conformations were sampled from the steady state. 3D-snapshots of individual polymer configurations were rendered using POV-Ray software (version 3.7).

### Bulk RNA-seq

Hearts from *H2*-URI^Δ/Δ^ and WT embryos at E9.5 were micro-dissected and collected in RNAlater (Sigma, R0901), incubated overnight at 4°C, and stored at -20°C. Following genotyping, three stage-matched hearts (n = 3) from *H2*-URI^Δ/Δ^ and WT embryos were selected for comparative analysis and processed for total RNA extraction using the Qiagen RNeasy Micro Kit (Cat. No. 74004). Per replicate, RNA quantity and integrity were evaluated using a Qubit 4.0 Fluorometer with the Qubit RNA HS Assay Kit (Thermo Fisher Scientific, Q32855) and a Fragment Analyzer System with the Fragment Analyzer RNA Kit (Agilent, DNF-471), respectively. Sequencing libraries were generated from 300 ng of total RNA using the TruSeq Stranded mRNA Library Prep Kit (Illumina, 20020595) in combination with TruSeq RNA UD Indexes (Illumina, 20022371), according to the manufacturer’s protocol. Pooled libraries were quantified using a Qubit 4.0 fluorometer with the Qubit dsDNA HS Assay Kit (Fisher Scientific, Q32854) and assessed for fragment size distribution on an Agilent Fragment Analyzer with the HS NGS Fragment Kit (Agilent, DNF-474). Equimolar amounts of each library were then pooled and sequenced in paired-end mode on an Illumina NovaSeq 6000 platform using the NovaSeq 6000 SP Reagent Kit v1.5 (300 cycles; Illumina, 20028400). Sequencing quality was monitored with the Illumina Sequencing Analysis Viewer (version 2.4.7), and base call files were demultiplexed and converted to FASTQ format using bcl2fastq v2.20. All library preparation, quality control, and sequencing were carried out by the NGS Platform at the University of Bern. RNA-seq reads were aligned to the mouse reference genome assembly GRCm39.110 (mm39) using STAR (version 2.7.9a)^176^. Gene-level read counts were obtained with the featureCounts function from the Subread package (version 2.0.1)^177^. Analysis of differentially expressed genes (DEGs) was performed using the DESeq2 package (version 1.30.1) within Bioconductor^178^. Genes with an adjusted p-value < 0.05 were considered significantly differentially expressed. Data exploration and visualization were conducted using the iBET package (version 0.0.0.9) in R (version 4.0.5). Scaled RNA-seq read count matrices of DEGs were visualized using the Morpheus web tool (https://software.broadinstitute.org/morpheus).

### Integration of bulk RNA-seq DEGs with single-cell multiome cluster profiles

For assignment, each multiome cluster, the mean expression of all cluster-specific multiome marker genes was computed across the bulk RNA-seq samples form the *H2*-URI^Δ/Δ^ versus WT comparison (n=3 per genotype). The expression of each marker gene was normalized by the average expression across all respective replicate samples. The average expression of up- and down-regulated DEGs in the *H2*-URI^Δ/Δ^ vs. WT bulk RNA-seq dataset was quantified and projected on datapoints of mouse multiome clusters using the *AddModuleScore* function from Seurat^179^. This function computes a module score by aggregating the expression levels of specified target genes and subtracting the aggregated expression of a control gene set to correct for background signal. The resulting module scores were mapped onto the multiome UMAP for spatial visualization and further displayed using violin plots grouped by cell cluster.

## DATA AVAILABILITY

Raw and processed files of next-generation sequencing (NGS) datasets generated as part of this study are available in the NCBI GEO database with the accession codes GSExxxxxx. Reference genomes Mouse GRCm38/mm10 or GRCm39/mm39 were used for alignment and comparisons. Correspondence and requests for reagents or materials should be addressed to M.O.

## ACKNOWLEDGEMENTS

We are grateful to P. Pelczar and his team at the University of Basel Center for Transgenic Models (CTM) for their excellent services including the generation of all genome-edited knockout mouse lines in this study, the establishment of the enSERT workflow and production of transgenic LacZ reporter embryos. We also thank the micro-injection team at LBNL for their excellent support in additional enhancer-reporter analysis. We are grateful to S. Fuochi, C. Detotto, Lois Zitzow and Alessandra Bergadano and their teams at the Experimental Animal Center (EAC) and Central Animal Facilities (CAF) of the University of Bern for excellent mouse care and strain coordination. Valuable initial bioinformatics support for multiome analysis was provided by Grigorios Georgolopoulos (Genevia Technologies Ltd). We are indebted to R. Zeller and A. Zuniga (DBM, University of Basel) for their generous support regarding the establishment of enSERT at the CTM and transgenic mouse embryo collections and we thank Evgeny Kvon (University of Irvine) for expert guidance on the enSERT workflow and providing the ZRS construct. We thank C. Mosimann and A. Burger (University of Colorado) for providing the *pCK036* plasmid. We are grateful to P. Nicholson and her team at the NGS platform of the University of Bern for RNA-seq library preparation and sequencing. We thank M. Docquier and her team from the iGE3 facility for preparation and sequencing of C-HiC libraries, and Fabian Blank and his team at the live cell imaging facility (LCI) at the DBMR for their support. This study was supported by grants from the Swiss National Science Foundation (SNSF) PCEFP3_186993 (to M.O.), 407940_206520 (NRP79) (to M.O. and I.B.) and Novartis Foundation for Medical-Biological Research (#21C183) (to M.O.). J.L-R. is supported by the MICINN grant PID2023-148267NB-I00. Research in the Andrey laboratory is supported by Swiss National Science Foundation grants PP00P3_210996 and 320030-231203 (to G.A). A.V. was supported by National Institutes of Health grants R01HL162304, R01HD114353 and R01HG003988. Research conducted at the E.O. Lawrence Berkeley National Laboratory was performed under US Department of Energy contract DE-AC02-05CH11231, University of California.

## AUTHOR CONTRIBUTIONS

M.O. and V.R. conceived the project and wrote the manuscript with input from the other authors. V.R. performed experimental and molecular analyses of mouse knockout and transgenic enhancer-reporter alleles with support of E.W., V.T., J.G. and V.Ra. J.T. performed all multiome-related computational analyses under the supervision of I.B. M.Z. conducted the (re)processing of C-HiC, ATAC-seq, and ChIP-seq datasets and contributed to the initial multiome analysis. A.E. and M.C. performed all C-HiC analyses including vn4C, polymer modeling and molecular dynamics simulations under the supervision of M.N. J.L.R, G.N. and J.G. conducted the multiome experiment including embryo collection, sample processing and library preparation. R.R.G. and V.R. performed experimental C-HiC in the G.A. lab. I.M. conducted zebrafish experiments in the N.M. lab. A.A., P.A. and H.G. carried out RNA-seq analysis. B.J.M. and J.A.A. conducted transgenic reporter analysis in mouse embryos in the A.V. lab. H.W. and B.A.F. performed LacZ embryo section analysis in the lab of A.B.F.

## CODE AVAILABILITY

This study made use of current community-accepted and benchmarked bioinformatic analysis methods which are cited in the main text or Methods section. No previously unreported custom computer code, mathematical or software algorithms were employed for data analysis.

## Supplementary Figures

**Figure S1.**
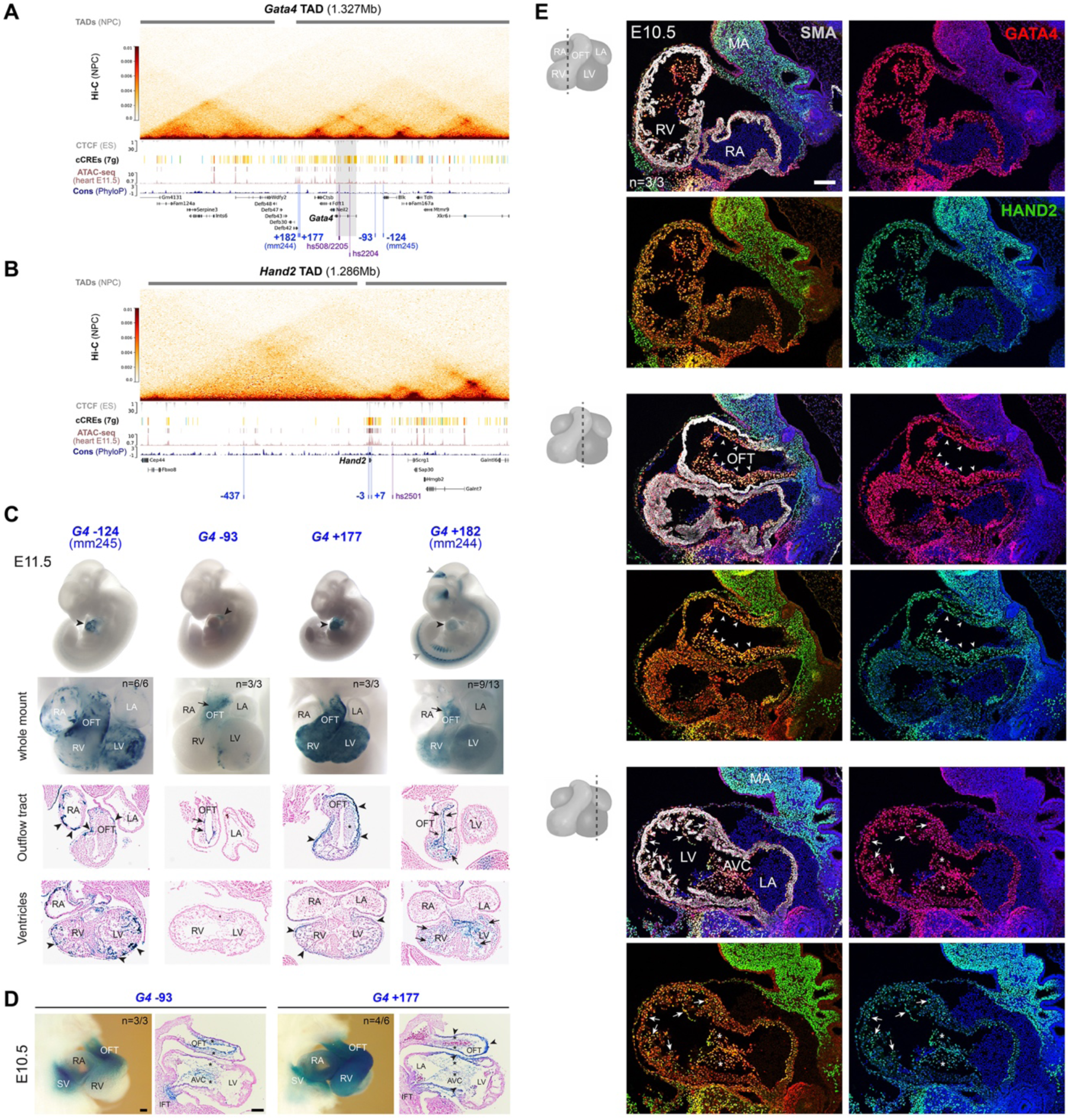
*Gata4* and *Hand2* genomic landscapes and protein distribution in mouse embryonic hearts. (**A, B**) Hi-C interaction counts of elements within *Gata4* (mm10, chr14:62494001-63821000) or *Hand2* (mm10, chr8:56523001-57809000) mouse topologically associated domains (TADs) in cortical neurons (CN) and illustrating TAD extension in neural progenitor cells (NPCs)^180^. Below, 7-group (7g) cCRE track from ENCODE (https://screen.encodeproject.org) listing promoter-like elements (red), enhancer-like sequences (ELS, yellow), CTCF-only sequences (blue) and DNase-only sequences (green) across tissues. CTCF track from mouse embryonic stem cells (ES)^180^, ATAC-seq profiles from mouse embryonic hearts at E11.5^53^ and placental mammal basewise conservation track (by PhyloP) are shown along with protein coding genes and previously identified heart enhancer elements (purple, tested human sequence; blue, tested mouse sequence). VISTA enhancer IDs are indicated (mm, mus musculus; hs, homo sapiens). Mouse enhancer elements are re-defined based on upstream (-) or downstream (+) distance (in kb) from associated *Gata4* (*G4*) or *Hand2* (*H2*) target genes: *G4*-124 (mm245)^61^, *G4*-93 (G9)^62^, *G4*+177 (identified in this study), *G4*+182 (mm244)^61^, *H2*-437^68^, *H2*-3 (*Hand2* -4.2 to -2.7 element)^60^ and *H2*+7 (mm1284)^63^. (**C**) Analysis of mouse *Gata4* heart enhancer activities in mouse embryos at E11.5. Black arrowheads in whole embryo panels point to reproducible staining in the heart. Gray arrowheads demarcate regions of additional reproducible reporter activity. Whole-mount panels: frontal view. Lower panels: frontal sections. Arrows point to reproducible staining in endothelial/endocardial cells. Arrowheads mark myocardial cells. (**D**) Whole mount (left) and Eosin-counterstained sagittal section (right) images demonstrating LacZ enhancer-reporter activity (blue) of *G4*-93 and *G4*+177 elements in the sinus venosus (SV) and OFT at E10.5, as resulting from enSERT analysis (see also Fig. 1B). Asterisks mark signal in mesenchymal cushions of the atrioventricular canal (AVC). Arrowheads indicate staining in the myocardial layer. (**E**) Sagittal sections showing co-localization of nuclear GATA4 (red) and HAND2 (green) in mouse wildtype (WT) hearts at E10.5. Schemes on the left indicate section plane. Yellow nuclei indicate GATA4/HAND2 co-expression. Nuclei are stained blue. “n” denotes fraction of biological replicates with reproducible results (for LacZ reporter transgenesis: reproducible tissue-specific staining vs. number of transgenic embryos with any LacZ signal). Scale bars, 100 μm. Coordinates and primer sequences of mouse elements analyzed via enSERT are listed in **Table S1**. Reproducible results from VISTA enhancers (mm244, mm245) are available at the VISTA Enhancer Browser (https://enhancer.lbl.gov/vista/). RA, right atrium. LA, left atrium. RV, right ventricle. LV, left ventricle.

**Figure S2.**
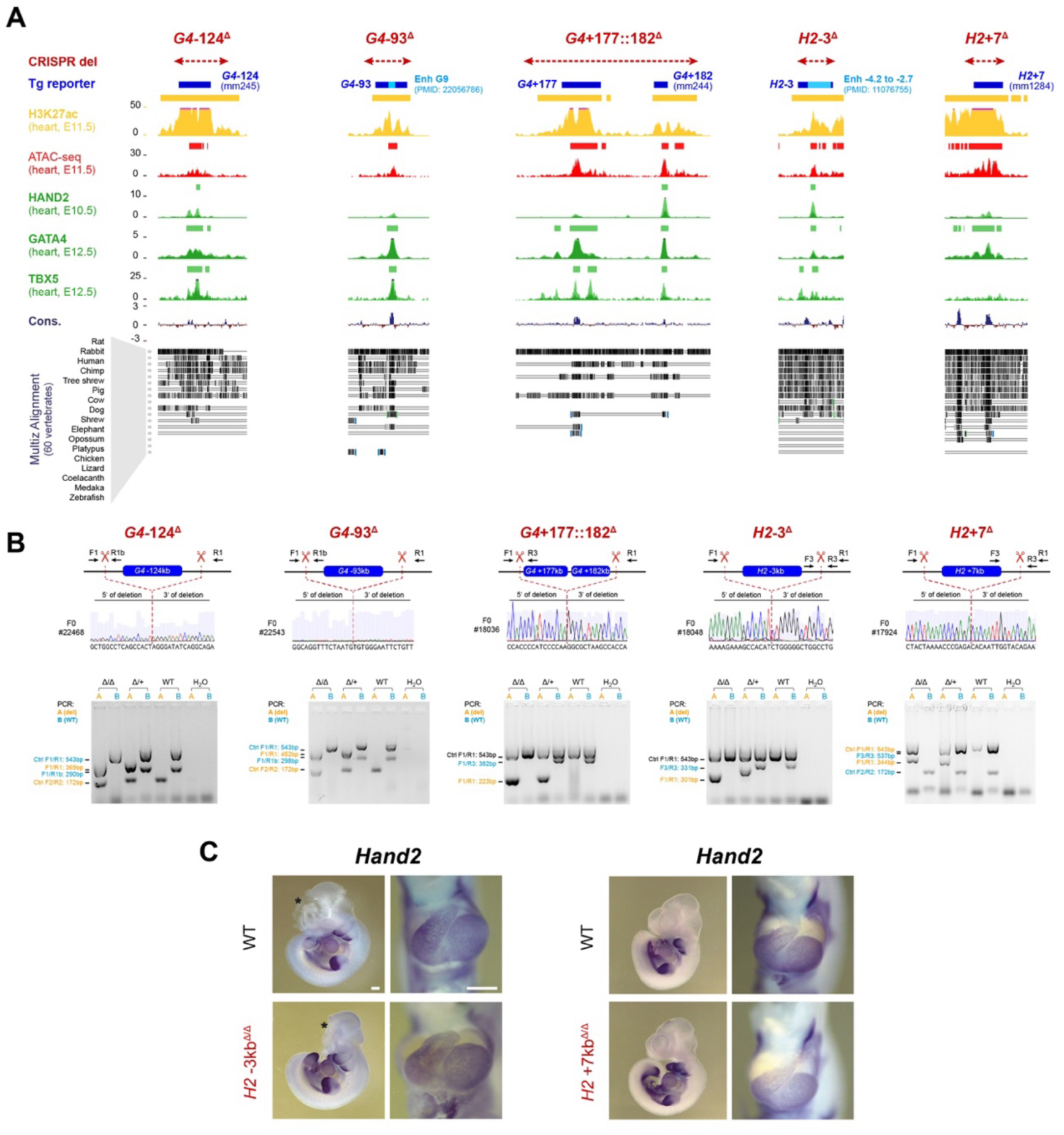
Epigenomic profiles of *Gata4*- and *Hand2*-associated heart enhancers and generation of endogenous deletion alleles in mice. (**A**) UCSC browser windows (mm10) of *G4*-124 (mm245)^61^, *G4*-93 (containing the G9 core)^62^, *G4*+177::182 (mm244)^61^, *H2*-3 (*Hand2* -4.2 to -2.7 element)^60^ and *H2*+7 (mm1284)^63^ heart enhancer regions revealing H3K27ac ChIP-seq (yellow)^70^, ATAC-seq^53^ (red) and available ChIP-seq profiles of cardiac core TFs (green) in embryonic hearts at E11.5 (H3K27ac, ATAC-seq), E12.5 (GATA4^74^, TBX5^75^) or E10.5 (HAND2^72^). Peak calls across replicates are indicated as bars on top of each track. The dark blue bar on top indicates the enhancer region used for transgenic analysis in previous studies via conventional transgenesis at E11.5 (*G4* -124, *G4*+182, *H2*+7) or analysis at E10.5 via enSERT in this study (*G4*-124, *G4*-93, *G4*+177). The light blue bar indicates the extension of corresponding shorter G9^62^ and *Hand2* -4.2 to -2.7^60^ elements analyzed by conventional transgenesis in previous studies. The red bar above indicates the extension of the genomic deletions (del) generated as shown in (B). Cons., conservation track by PhyloP. Corresponding VISTA Enhancer IDs (mm, mus musculus) are listed. +/-: downstream (+) or upstream (-) distance (in kb) from target gene TSS. (**B**) Generation of five mouse cardiac enhancer deletion alleles (*G4*-124^Δ^, *G4*-93^Δ^, *G4*+177::182^Δ^, *H2*-3kb^Δ^ and *H2*+7kb^Δ^) using CRISPR-EZ^73^. Upper panels: red scissors indicate representative CRISPR-mediated deletion breakpoints verified by Sanger sequencing (dashed line) in F0 and F1 animals per deletion line. Lower panels: a specific PCR genotyping strategy was designed for detection of wild-type (+) and enhancer deletion (Δ) alleles in F0 and subsequent generations (amplicon sizes indicated on the left). Primers amplifying an unrelated genomic region were included as positive controls. SgRNA and primer sequences are listed in **Tables S2 and S3**, respectively. (**C**) ISH revealing unchanged spatial *Gata4* or *Hand2* expression in hearts of *G4*+177::182^Δ/Δ^, *H2*-3kb^Δ/Δ^ and *H2*+7kb^Δ/Δ^ embryos compared to wildtype (WT) controls at E10.5. For each genotype, n=3 embryos were analyzed, with similar results. Scale bars, 500 μm (embryos) and 100 μm (heart close-ups). “n” denotes fraction of biological replicates with reproducible results. *G4*, *Gata4*. *H2*, *Hand2*.

**Figure S3.**
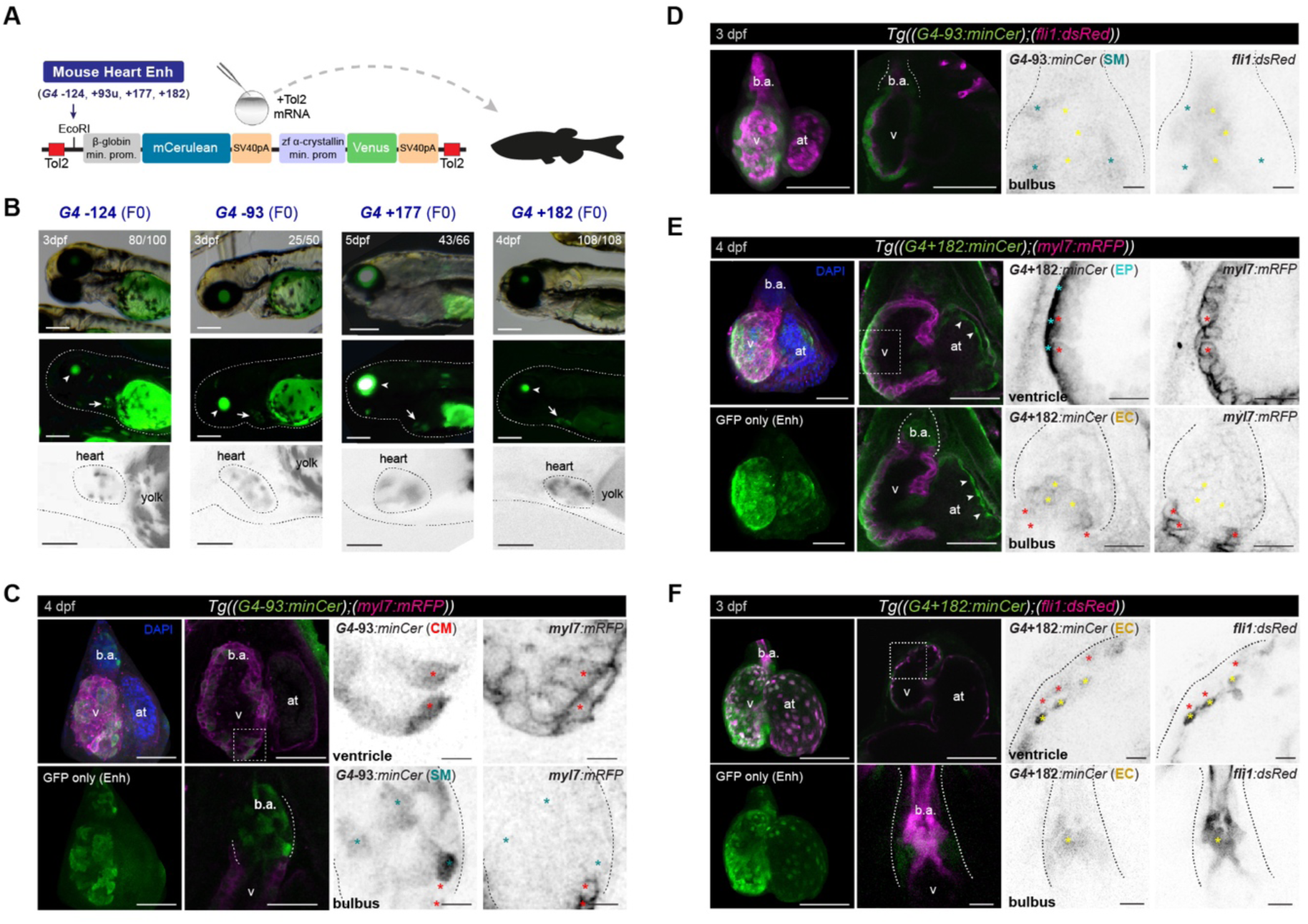
Partially conserved cell type specificities of mouse *Gata4* heart enhancers in zebrafish. (**A**) Schematic representation of enhancer reporter constructs injected into 1-cell stage zebrafish zygotes for generation of F0 transgenics. A mouse beta-globin minimal promoter-mCerulean (minCER) reporter cassette (pCK036)^80^ is used for detection of enhancer-mediated activity via the GFP channel. The secondary α-crystallin (cryaa) promoter-driven Venus reporter unit drives specific signal in the lense and allows identification of animals with an integrated transgene^80^. (**B**) Detection of enhancer-mediated cardiac activities in transgenic founder (F0) zebrafish embryos (**Table S5**). The fraction of embryos with GFP signal in the heart (arrow) compared to the number of total transgenic embryos based on GFP in the lense (arrowhead) is indicated (scale bars, 200 μm). Close-up images of the heart with inverted grayscale for the fluorescence signal are shown (scale bars, 100μm). (**C-F**) Representative confocal images of double transgenic immune labelled hearts showing G4-93 or G4+182 minCer enhancer-reporter activities along with staining for cardiomyocytes (CMs) marked by myl7:mRFP (magenta) (C, E) or endocardial/endothelial cells (EC) marked by the fli1:dsRed transgene (magenta) (D, F). Left-most panels display 3D projections of the fluorescent channels. The middle-left panels present cross-sections of ventricle (v) or bulbus arteriosus (b.a.) cardiac compartments. The panels on the right show single-channel images of the zoomed-in regions indicated by dashed lines with cell types marked by reporter activity indicated in brackets. Asterisks mark CMs (red), ECs (yellow), epicardial cells (EP) (cyan) and SMs of the b.a. (teal). Arrowheads in (E) mark activity in atrial (at) ECs. Fluorescent signals represent reproducible patterns detected in n=11/12 (founder 1) and n=11/16 (founder 2) embryos from stable G4-93kb-minCER;myl7:mRFP transgenic lines (C), n=6/6 embryos from a stable G4-93kb-minCER;fli1:dsRed line (D), n=12/12 embryos from a stable G4+182kb-minCER;myl7:mRFP line (E) and n=3/6 from a stable G4+182kb-minCER;fli1:dsRed line (F). Images show ventral views on the heart, with the head oriented to the top. DAPI-stained nuclei (C, E) are blue. Scale bars: 100 µm (hearts, left panels), 10 µm (zoomed views, right panels). dpf, days post-fertilization.

**Figure S4.**
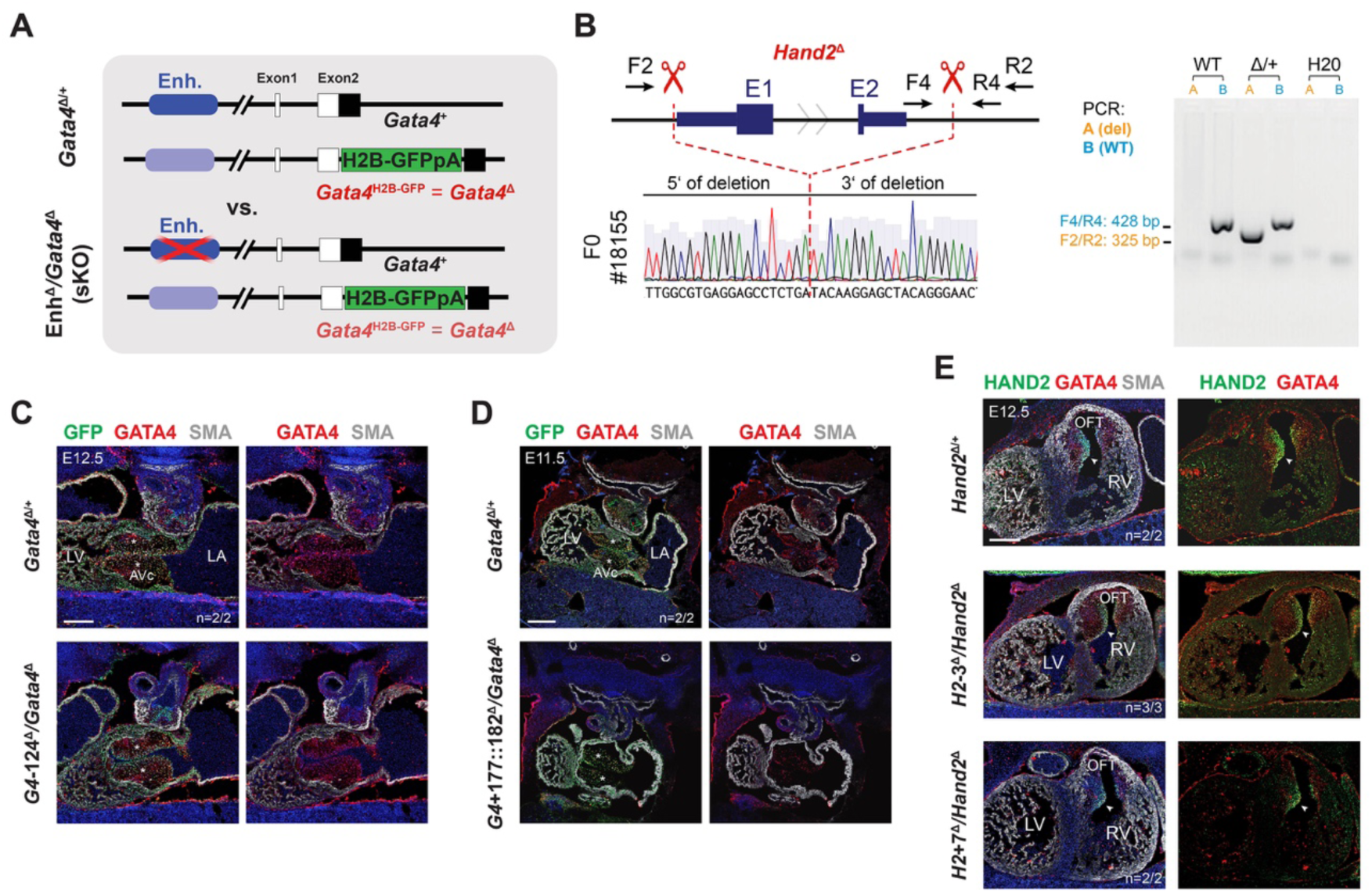
*Gata4* and *Hand2* loss-of-function alleles and molecular assessment of genetically sensitized cardiac enhancer deletions in mouse embryonic hearts. (**A**) Allelic framework enabling *Gata4* loss-of-function for comparison of sensitized *Gata4* enhancer knockouts (sKO) and *Gata4* heterozygous (het) controls. The *Gata4*^Δ(H2B-GFP)^ allele representing a *Gata4* Null allele that at the same time is reporting *Gata4* transcriptional activity via nuclear GFP^82^. (**B**) Generation of a *Hand2* knockout (*Hand2*^Δ^) allele in the FVB/NRj background isogenic to the enhancer deletion strains. Left: Sanger sequencing traces of the *Hand2*^Δ^ allele in F0 (founder) and F1 animals with deletion breakpoints marked by dashed red lines. Right: PCR genotyping with primers listed in the scheme on the left. Del, deletion. WT, wildtype. (**C, D**) Immunofluorescence (IF) co-localization of GATA4 protein (red), Gata4 transcriptional activity (green) mediated by the *Gata4*^Δ(H2B-GFP)^ allele^82^ and the myocardial marker smooth muscle actin (SMA, gray) in sagittal sections of hearts from *G4*-124^Δ^/*Gata4*^Δ(H2B-GFP)^ embryos at E12.5 (left) and *G4*+177::182^Δ^/*Gata4*^Δ(H2B-GFP)^ embryos at E11.5 (right), each compared to *Gata4*^Δ(H2B-GFP)/+^ controls. Asterisk marks AV cushion (AVc) mesenchyme. (**E**) IF staining of HAND2 (green), GATA4 (red), SMA (gray) on sagittal sections of the left ventricle (LV) and outflow tract (OFT) of hearts from H2-3^Δ^/*Hand2*^Δ^ (left) or *H2*+7^Δ^/*Hand2*^Δ^ (right) embryos compared to *Hand2*^Δ/+^ controls at E12.5. Arrowhead points to strong HAND2 domain in OFT cushions. Scale bars, 200μm.

**Figure S5.**
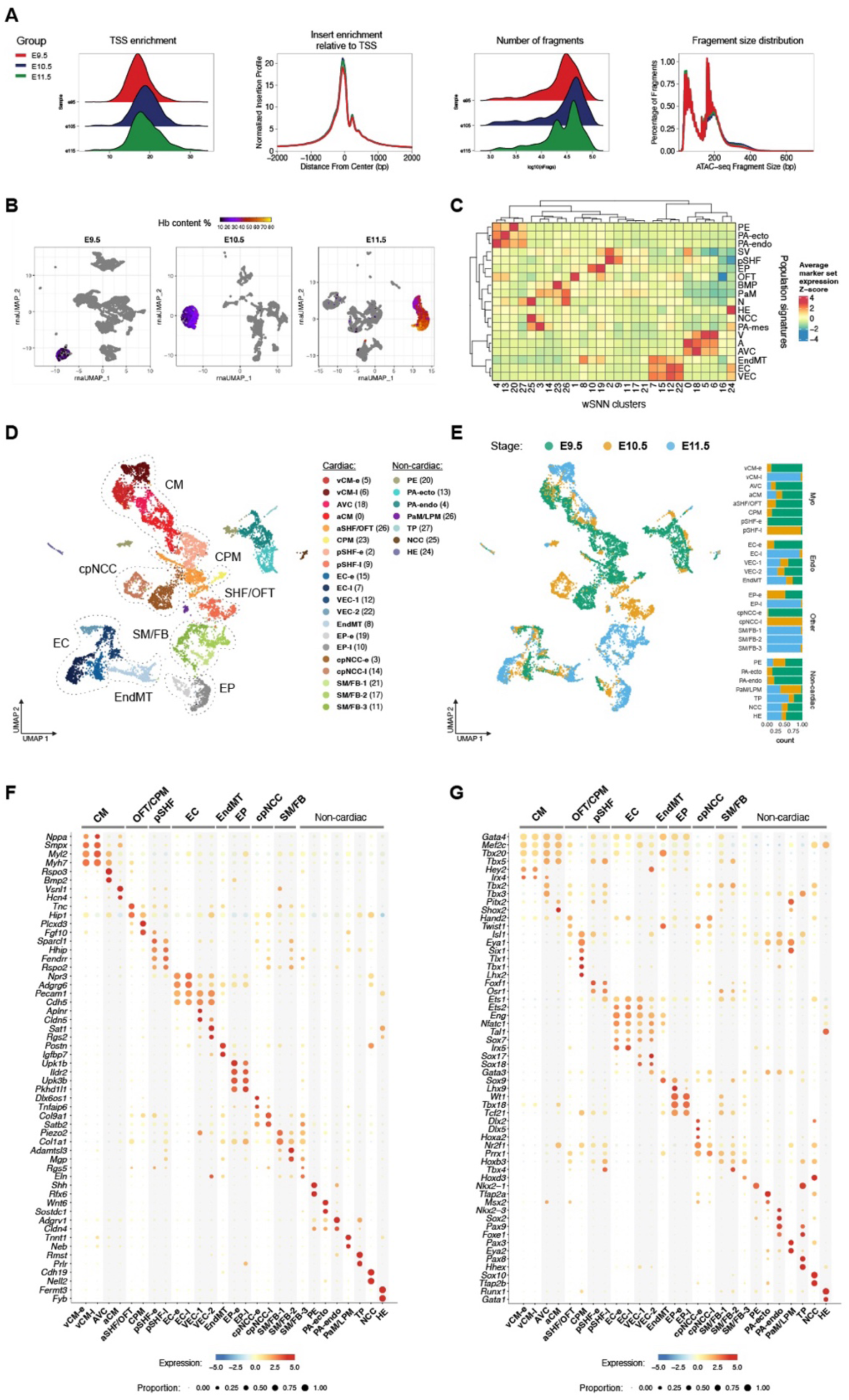
Single-cell multiome filtering and annotation of cell type identities. (**A**) SnATAC-seq quality control plots from embryonic hearts and associated tissues at different stages. (**B**) UMAP using the snRNA-seq signal alone, highlighting the cells with a high fraction of reads mapping to hemoglobin genes (Hb). Dashed lines indicate broader lineage definitions. (**C**) Heatmap showing the expression enrichment for the indicated gene sets (rows) across the indicated clusters (columns), based on sets of marker genes from previous studies of the early embryonic heart (E9.25/E10) and adjacent tissues (E10)^32,84^ (**Table S6**). SV, sinus venosus. BMP, branchiomeric muscle progenitors. N, neuron. Mes, mesoderm. V, ventricle. A, atria. (**D**) UMAP coordinates integrating all high-quality filtered cells from both RNA and ATAC datasets (28 clusters in total), after red blood cell (RBC) removal. Definitive cell type cluster annotations based on (C) and considering additional markers from previous studies (**Table S6**) as well as temporal signatures (E) are listed. Respective cluster numbers from (C) are listed in brackets. Dashed lines indicate broader cell type and lineage definitions. (**E**) Left: UMAP projection as in (D) but colored by embryonic stage. Right: Stacked bar plots showing the fraction of cells per stage in each multiome cluster (**Table S7**). (**F, G**) Bubble plots featuring expression signatures of non-TF (F) and TF (G) cell type/lineage markers used for annotation of cluster identities in line with previous single cell studies from embryonic mouse hearts including myocardial and endocadial cell types^32,84^ (**Table S6**). In addition, cells of the outflow tract (OFT) cluster are marked by a characteristic *Isl1*/*Hand2*/*Tnc* signature^32^. Cardiopharyngeal mesoderm progenitors (CPM) giving rise to cardiac and branchiomeric muscle are marked by a characteristic *Tbx1*/*Tlx1*/*Lhx2* signature^87,90^. Vascular endothelial cell (VEC) clusters are marked by high levels of *Sox17* and *Sox18*^50^. The EndMT cluster represents cells derived from endocardial-to-mesenchymal transition marked by *Twist1*, *Sox9* and *Tbx20*, as well as *Postn* known as an AV mesenchyme determinant^29,85^. The cardiopharyngeal neural crest-derived cell population (cpNCC) shows a previously reported *Tbx2/3* signature, along with OFT-cardiac NCC genes such as *Isl1*, *Gata3* and *Hand2*^18^, while the SM/FB-3 cluster displays a related *Gata3*, *Hoxb3* and *Rgs5* profile indicating (partial) neural crest origin^18^. In contrast, the SM/FB1 cluster exhibits *Wt1*, *Tbx18* and *Tcf21*, pointing to a population of epicardial-derived cells (EPDCs), likely developing into cardiac fibroblasts (FB) and vascular smooth muscle cells (vSM)^181^. a/vCM, atrial/ventricular cardiomyocytes. AVC, atrioventricular canal. a/pSHF, anterior/posterior second heart field-derived cells. SM/FB, smooth muscle/fibroblast-like cells. EP, epicardial cells. EC, endocardial cells. EndMT, endocardial-to-mesenchymal transition/transformed cells of the cushions. PE, pulmonary endoderm. PA-endo/ecto, pharyngeal arch endo-/ectoderm. PaM/LPM, paraxial/lateral plate mesoderm. TP, thyroid progenitor-like cells. NCC, neural crest cells. HE, hematopoietic lineage cells. -e, early (E9.5 or E10.5). -l, late (E10.5 or E11.5). E, embryonic day.

**Figure S6.**
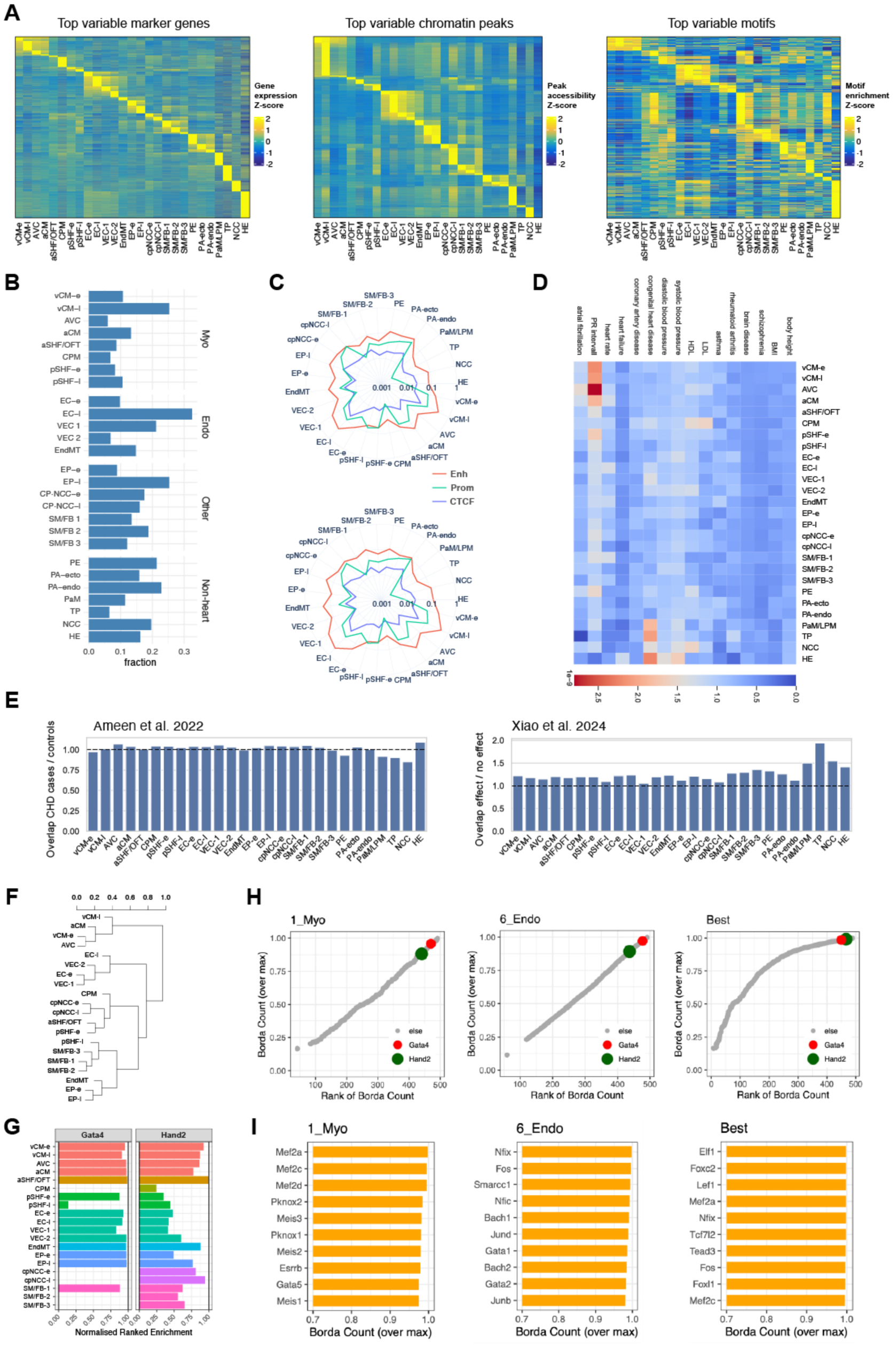
Characterization of the cardiac cistrome across cell types and identification of underlying regulators. (**A**) Heat maps showing the gene- and region-wise normalized expression or accessibility values, for the indicated top variable features (n=24333 genes, n=50000 peaks, n=1768 motifs), across the indicated clusters. (**B**) Bar plots showing the fraction of differentially accessible peaks per multiome cluster (**Table S7**). (**C**) Radar plots displaying the proportion of differentially accessible peaks for each cluster, categorized by regulatory identity, using ChromHMM annotations at E10.5 (top) and E11.5 (bottom) for enhancer, promoter, or insulating (CTCF-only) elements (see Methods) (**Table S7**). (**D**) Heatmap showing the normalized overlap of GWAS top associations with cluster-specific peaks (**Table S9**). (**E**) Bar plots showing the normalized enrichment of CHD variants in the regulatory regions in cardiac multiome clusters, using CHD variants from two studies^39,50^ (**Table S9**). Ameen at al. 2022: the plot shows the ratio between the overlap of CHD variants versus control mutations, within cluster-specific peaks. Xiao et al. 2024: the plot shows the ratio between the overlap of the variants with experimental effect versus variants without effect, within cluster-specific peaks. (**F**) Hierarchical clustering of the TF-binding sites enrichment across the cistrome of all cardiac cluster populations. (**G**) Barplot showing the ranked enrichment for the indicated TFBS models, per cluster. **(H, I**) Pioneer factor analysis, considering only the myocardial clusters (left), the endocardial clusters (middle), or all the clusters (right). Scatterplots (top) show the ranked TFs (candidates with higher scores to the right) based on their pervasiveness and strength of enrichment across the heart populations. Barplots (bottom) highlight the top 10 scoring TFs in each analysis. a/vCM, atrial/ventricular cardiomyocytes. AVC, atrioventricular canal. a/pSHF, anterior/posterior second heart field-derived cells. SM/FB, smooth muscle/fibroblast-like cells. EP, epicardial cells. EC, endocardial cells. VEC, vascular endothelial cells. EndMT, endocardial-to-mesenchymal transition/transformed cells of the cushions. PE, pulmonary endoderm. PA-endo/ecto, pharyngeal arch endo-/ectoderm. PaM/LPM, paraxial/lateral plate mesoderm. TP, thyroid progenitor-like cells. NCC, neural crest cells. HE, hematopoietic lineage cells. -e, early (E9.5 or E10.5). -l, late (E10.5 or E11.5). E, embryonic day.

**Figure S7.**
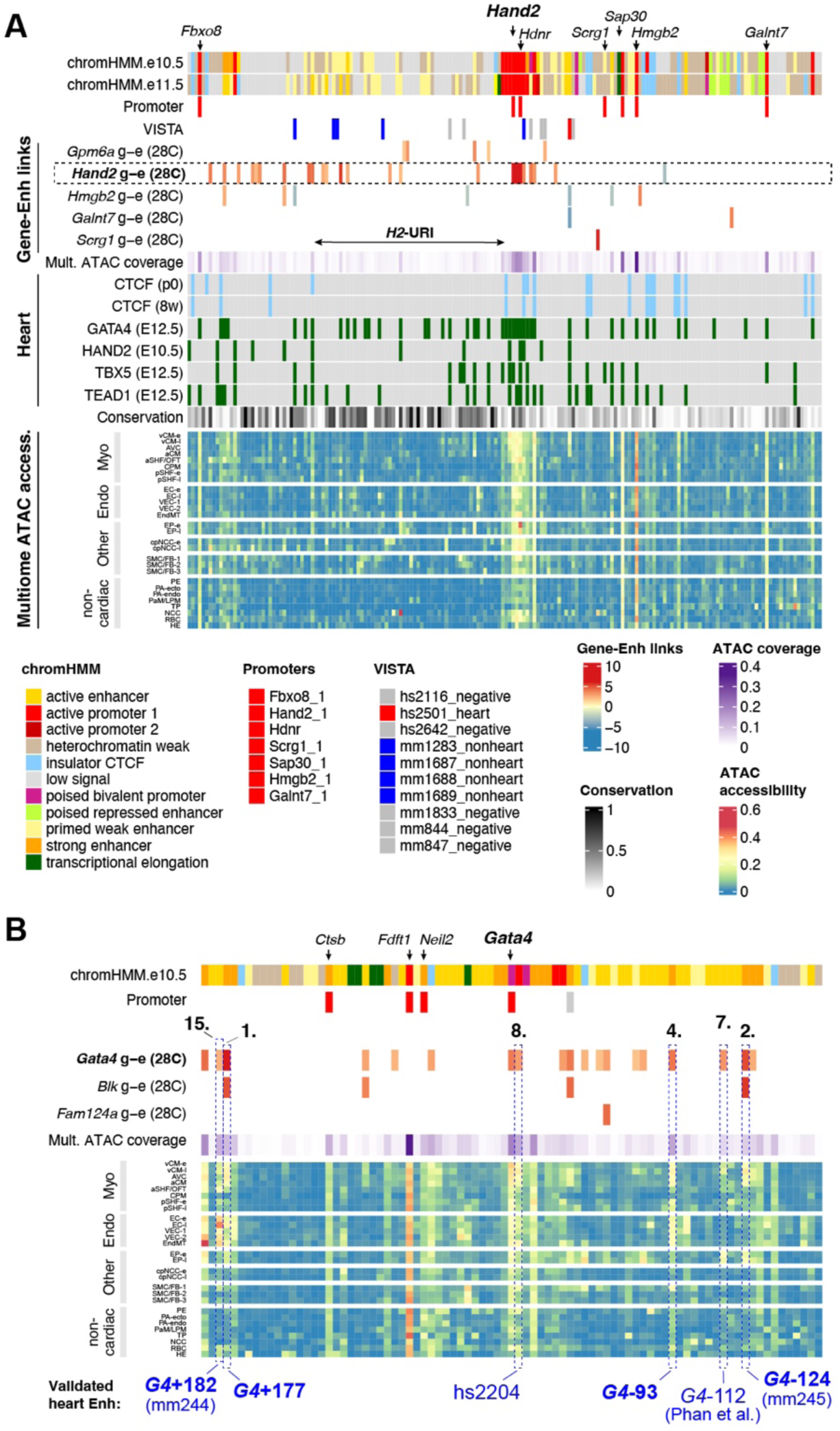
A user-accessible interface to interrogate multiome-based cardiac regulatory landscapes genome-wide. (**A**) As an example for any locus visualization genome-wide, the extended *Hand2* TAD (mm10; chr8:56,523,001-57,809,000) (**Fig. S1**) is shown. A comprehensive interface presents a heatmap of chromatin accessibility elements across cardiac cell type-specific multiome clusters (**Table S11**). Epigenomic ChromHMM profiles and *in vivo*-validated regulatory elements (VISTA Enhancer Browser) are listed for comparison. Gene promoter IDs and related multiome-computed gene-enhancer links (g-e) establish correlation of target gene expression and putative enhancer activity (see Methods). Average ATAC-signal across clusters (coverage) and conservation score (PhastCons) are shown for each element along with significant CTCF^70^ or TF occupancy from published ChIP-seq datasets^72,74,75^. *H2*-URI, *Hand2* upstream regulatory interval (see Fig. 4A). E, embryonic day. P, postnatal day. W, weeks. (**B**) Exploration of the *Gata4* locus (mm10; chr14:63,055,001-63,423,858) (**Table S11**). Previously validated *Gata4* heart enhancers^61,62,68^ are identified by top-ranked *Gata4* g-e links, with multiome accessibility profiles matching the cardiac cell type enhancer activities detected *in vivo* (**Figs. 1, S1**). Red-orange shading denoting levels of stringency based on correlation values. Computed g-e links, including profiles from all 28 clusters (28C) and all genes within the *Gata4* TAD, are shown.

**Figure S8.**
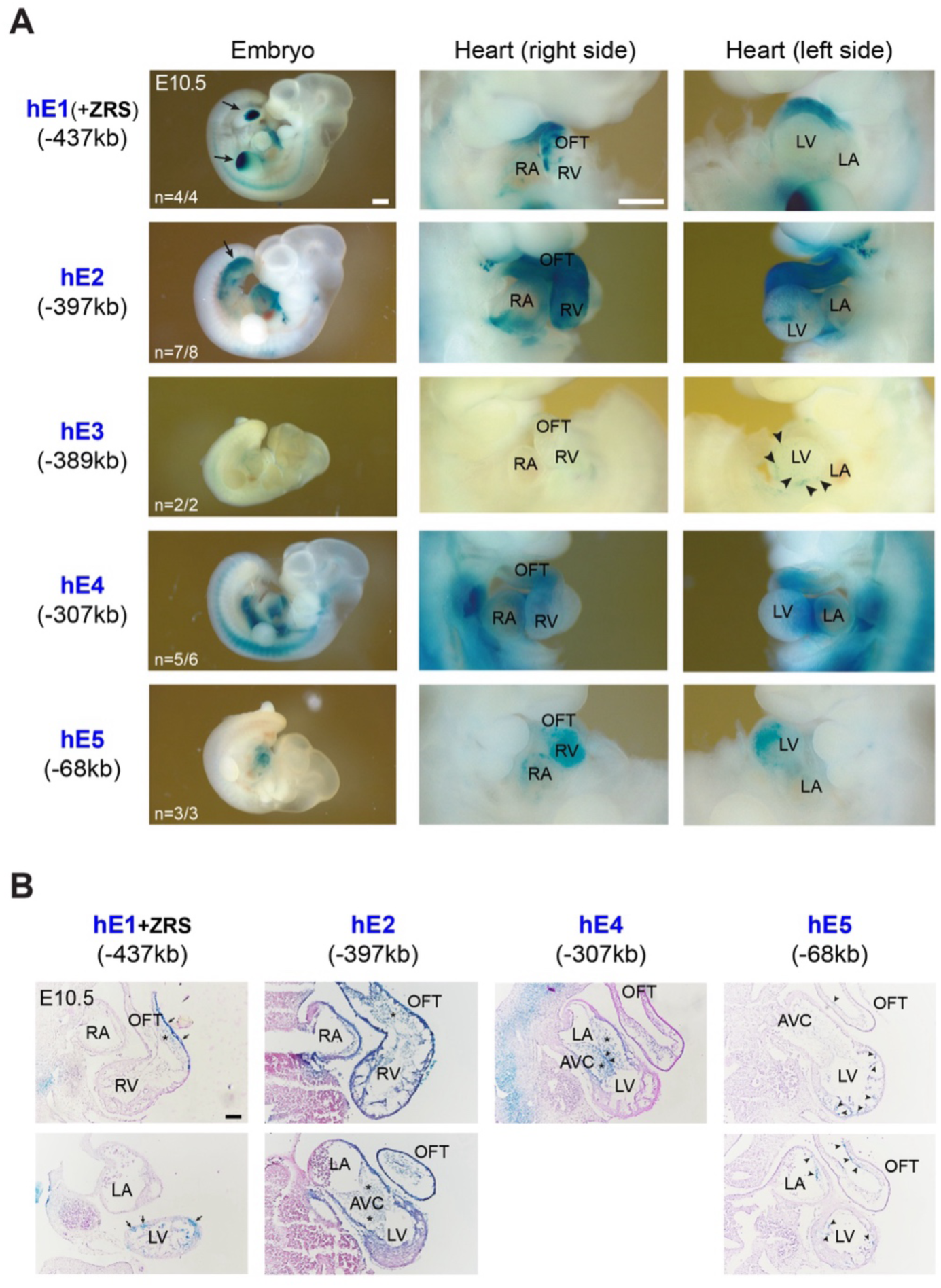
Compartment and cell type-specific activities of identified Hand2 heart enhancers. (**A**) Transgenic LacZ reporter validation of predicted enhancer elements within the *Hand2* upstream chromatin domain at E10.5 (Fig. 4; **Table S12**). Upstream distance (-) of each element from *Hand2* TSS is indicated (in kb). Per element, one embryo with representative LacZ staining is shown. Numbers in the bottom indicate the number of embryos with reproducible cardiac staining versus the number of transgenic embryos with any LacZ signal. HE1 was analyzed in tandem with the ZRS limb enhancer element^69^ driving enhancer activity exclusively in the posterior limb mesenchyme. Arrows point to reproducible staining in limbs. For hE3 only surface signal in single cells of the epicardial layer was detected (arrowheads). (**B**) Eosin-stained (red) sections of X-gal-treated embryos show enhancer-driven LacZ reporter activity (blue) localized to the myocardium, endocardium, or endocardial (EC) cushions, as identified by enSERT analysis. Arrows and arrowheads point to myocardial or EC staining, respectively. Asterisk marks enhancer activity in EC cushion mesenchyme. OFT, outflow tract. RA, right atrium. LA, left atrium. RV, right ventricle. LV, left ventricle. AVC, atrioventricular canal.

**Figure S9.**
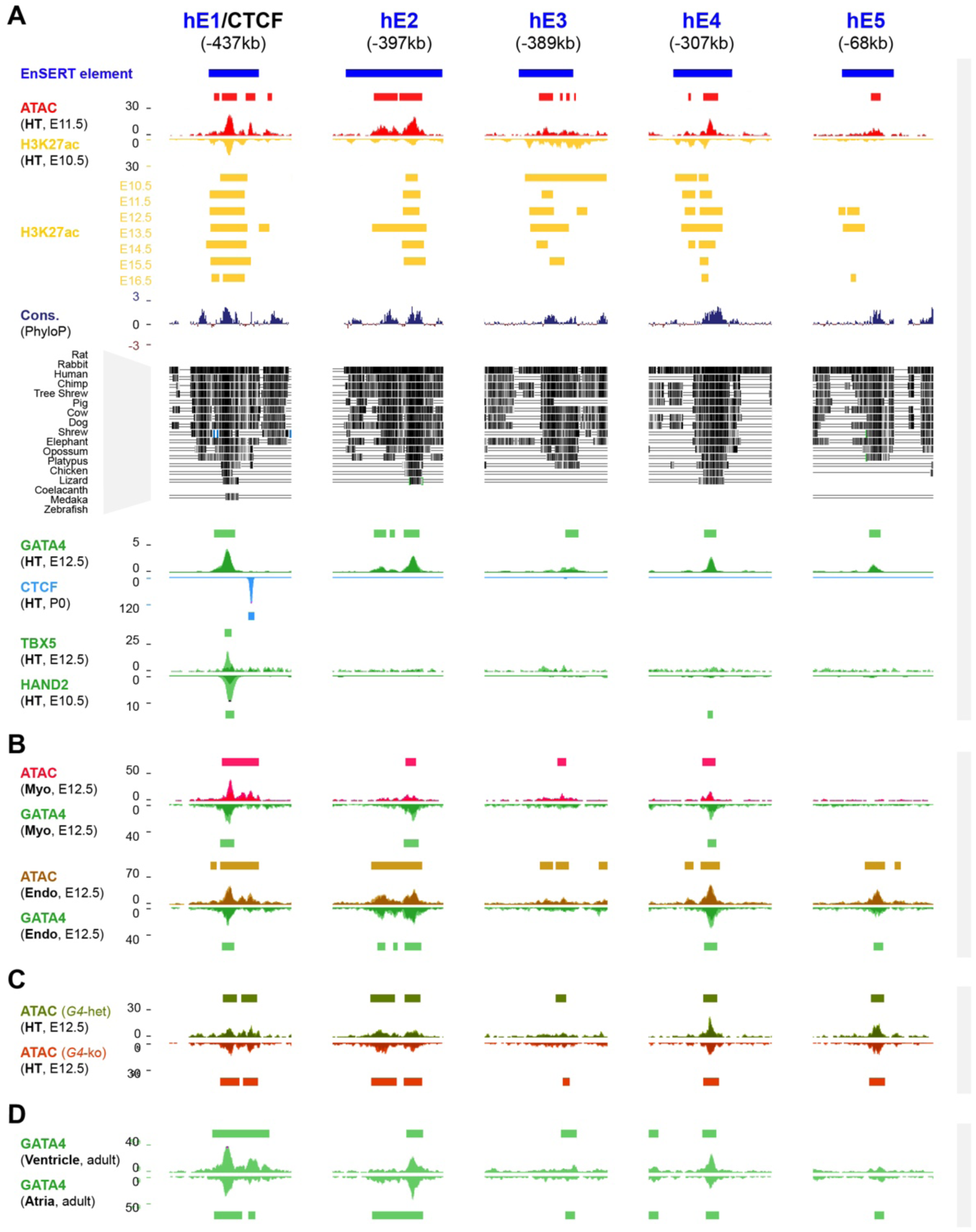
TF binding and chromatin features of the identified *Hand2* upstream heart enhancers. (**A**) Upper tracks: Genomic extension (blue bars) of the cardiac enhancer elements identified within the *Hand2* upstream region (Fig. 4**, Table S12**). Respective open chromatin (ATAC-seq, E11.5, red)^53^ (**Table S13**) and H3K27ac (E10.5-E16.5, yellow)^70^ profiles in mouse embryonic hearts are shown along with placental mammal basewise conservation (Cons) by PhyloP (UCSC Genome Browser). Lower tracks: Reprocessed ChIP-seq profiles of GATA4 and TBX5 in mouse hearts at E12.5^74,75^, and of HAND2 at E10.5^72^. CTCF from P0 hearts^70^ (light blue) is shown for comparison. (**B**) Accessible chromatin profiles from myocardial (red) and endocardial (dark yellow) cells from mouse embryonic hearts at E12.5 with respective GATA4 occupancy^51^ (**Table S13**). (**C**) Accessible endocardial chromatin signatures from ventricles of conditional *Cdh5*-CreERT2 *Gata4* mutant (ko) vs. control (het) hearts at E12.5^51^ (**Table S13**). (**D**) GATA4 ChIP-seq profiles in ventricles or atria of adult mouse hearts^104^ with selected conservation tracks (**Table S13**). Peak calls across replicates are indicated as bars on top of each track.

**Figure S10.**
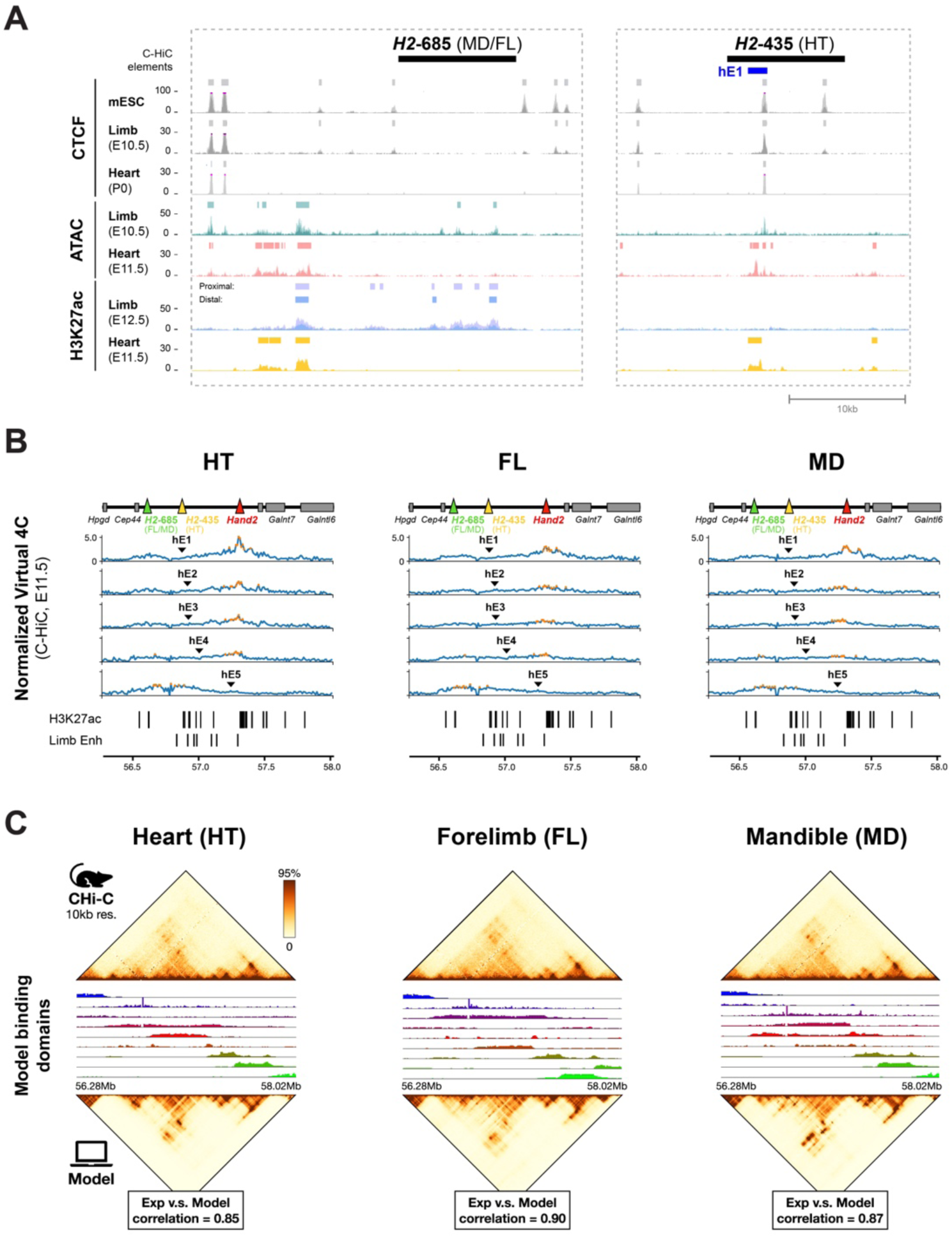
Heart enhancer interactions and tissue-specific SBS models for analysis of *Hand2*-associated chromatin dynamics. (**A**) UCSC Genome Browser tracks displaying CTCF, accessible chromatin (ATAC) and H3K27ac profiles across the most prominent *Hand2*-interacting regions (Fig. 5) in developing limbs and hearts, respectively^53,70,141,153,182^ (**Table S13**). CTCF enrichment in mouse embryonic stem cells (mESC)^180^ is shown for comparison. (**B**) Normalized C-HiC virtual 4C tracks from heart (HT), forelimb (FL), and mandible (MD) tissues (see Methods), with the identified heart enhancers (hE1-hE5) (Fig. 4) used as viewpoints. The position of the most predominant *Hand2*-interacting elements are denoted by green (FL, MD) and yellow (HT) triangles (Fig. 5**; Table S14**). Red triangle demarcates the position of the *Hand2* TSS. Outliers above the 95^th^ percentile are marked in orange. Respective tissue-specific H3K27ac peaks (ENCODE) and previously identified limb enhancers^100,101^ are listed for comparison. (**C**) C-HiC contact maps of the *Hand2* locus in E11.5 HT, FL and MD tissues (top, 10 kb resolution) and their corresponding SBS polymer models (bottom) show high similarity, with Pearson correlation coefficients of r = 0.85, 0.90, and 0.87, and distance-corrected Pearson coefficients (r′) of 0.52, 0.59, and 0.58, respectively. The bar plots illustrate the genomic locations and relative frequencies of nine distinct types of model binding sites across the analyzed tissues (see Methods). Each binding domain is represented by a unique color.

**Figure S11.**
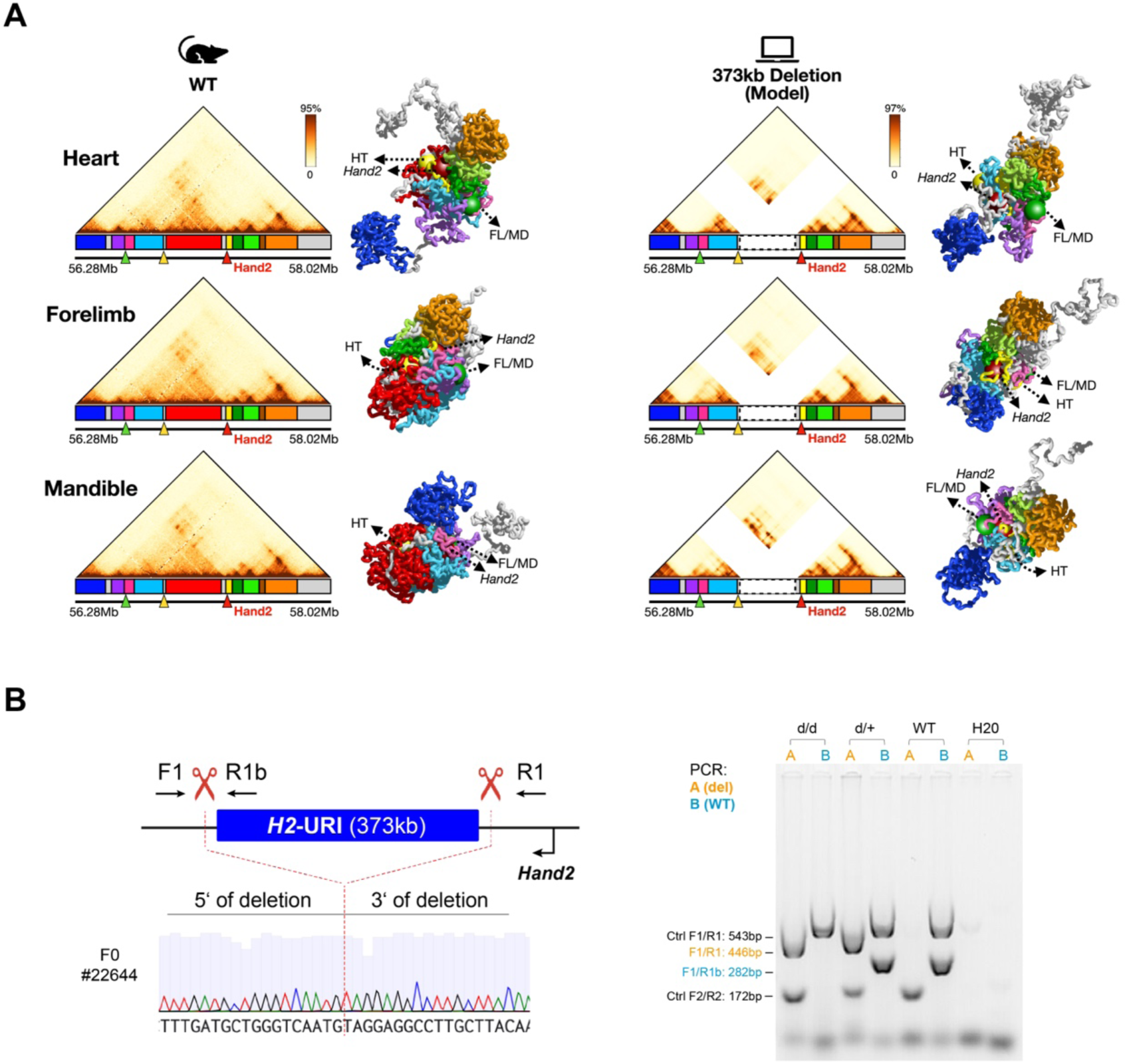
Genomic engineering of a mouse model lacking the *Hand2* upstream regulatory interval (*H2*-URI). (**A**) Predicted structural impact of a defined 373kb deletion within the non-coding upstream regulatory interval of *Hand2* (*H2*-URI^Δ^; chr8:56903578–57277013, mm10), as modeled using polymer simulations based on C-HiC datasets from heart (HT), forelimb (FL), and mandible (MD) tissues (**Figs. 5, S10**). Model-derived contact matrices and 3D chromatin conformations indicate that the locus retains its core structural organization despite the deletion. (**B**) CRISPR-engineering of the *H2*-URI^Δ^ allele in mice (**Table S2**, see Methods). Left: Sanger sequencing traces illustrating deletion breakpoints (dashed red lines), with genotyping primers (**Table S3**) indicated. Right: PCR genotyping to distinguish *H2*-URI^Δ/Δ^, *H2*-URI^Δ/+^ and WT control embryos.

**Figure S12.**
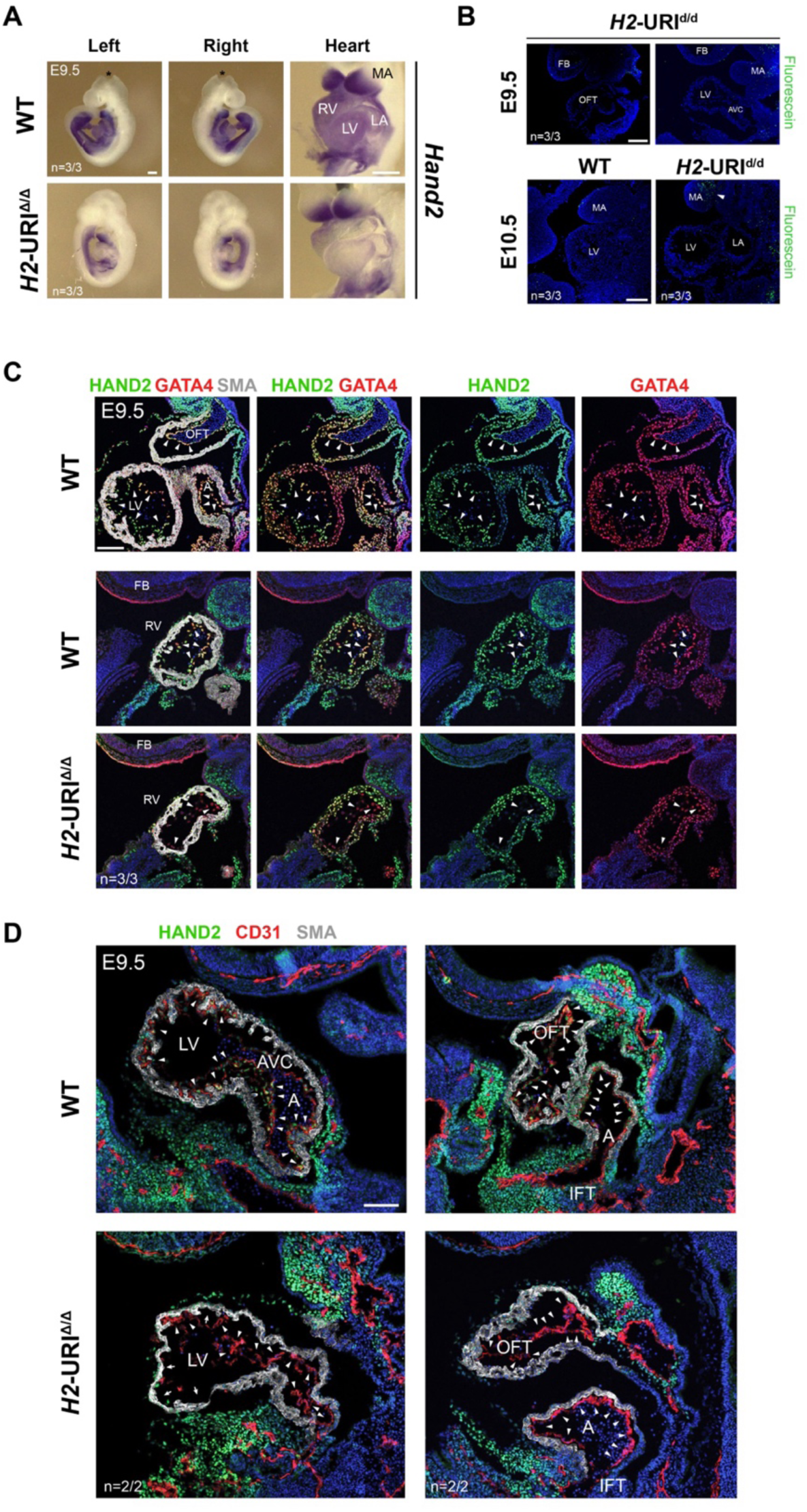
Endogenous loss of the *H2*-URI results in abolishment of HAND2 in the endocardial lineage. (**A**) ISH revealing downregulation of *Hand2* expression in the heart of *H2*-URI^Δ/Δ^ embryos at E9.5 (see also Fig. 6). *Hand2* expression in the trunk lateral plate mesenchyme, mandibular arch (MA) and outer portion of the outflow tract (OFT) of *H2*-URI^Δ/Δ^ embryos is partly retained. Asterisk indicates brain tissue removed for genotyping. Scale bars, 250 μm. (**B)** TUNEL analysis for detection of apoptotic cells. Sagittal sections of *H2*-URI^Δ/Δ^ vs. WT heart at E9.5 (top) and E10.5 (bottom), respectively. No enrichment of apoptotic cells was detected in hearts of *H2*-URI^Δ/Δ^ embryos compared to WT controls. White arrowhead points to apoptotic cells in the mandibular arch (MA). Scale bars, 200 μm. (**C**) Immunofluorescence (IF) co-localization of HAND2 (green) and GATA4 (red) in hearts of *H2*-URI^Δ/Δ^ embryos at E9.5. Similar to the endocardial cells (ECs) of the left ventricle (LV) and atrium (A) (Fig. 6E), ECs of the right ventricle (RV) in *H2*-URI^Δ/Δ^ embryos lack HAND2 (arrowheads), whereas the myocardium displays only a partial reduction. Smooth muscle actin (SMA) marks the myocardial layer (gray). Loss of ECM bubbles enclosing the forming trabeculae is also detected in mutants. Scale bar, 100 μm. (**D**) Co-localization of HAND2 (green) and CD31 (red) demonstrates absence of HAND2 in CD31-positive ECs (arrowheads) lining the inner wall of chamber and OFT myocardium (marked by SMA, gray) in *H2*-URI^Δ/Δ^ embryos. Scale bar, 100 μm.

**Figure S13.**
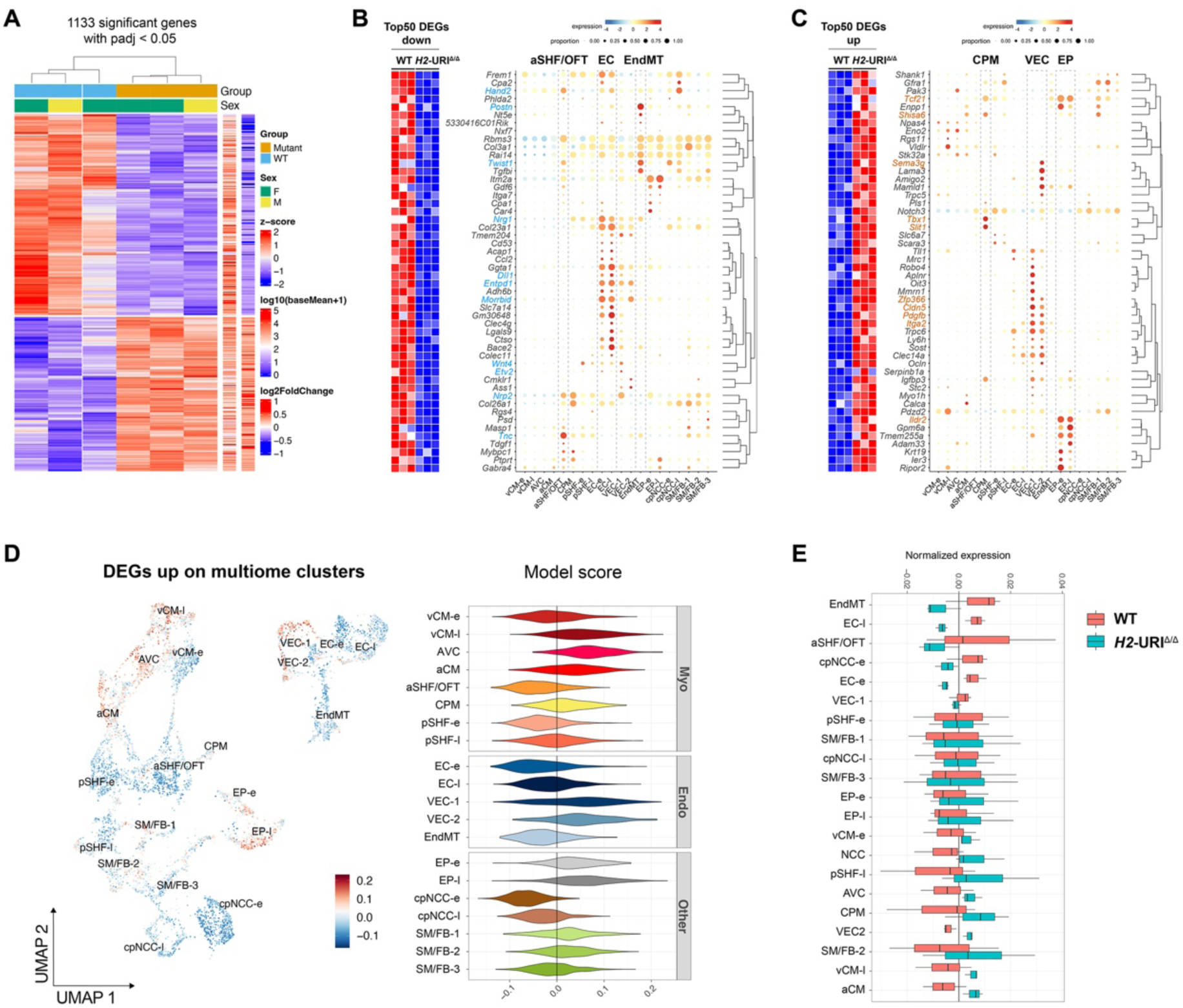
Transcriptomic alterations following loss of the *H2*-URI. (**A**) Heatmap revealing the total set of upregulated and downregulated differentially expressed genes (DEGs) resulting from bulk RNA-seq of embryonic hearts of *H2*-URI^Δ/Δ^ embryos compared to WT controls at E9.5 (see also Fig. 6). N=1133 significantly altered genes with padj. < 0.05 were identified (**Table S16**). (**B**) Top 50 downregulated genes (padj. < 0.05, log2FC > -0.5) clustered according to gene expression signatures are enriched for markers of outflow tract (OFT), endocardial (EC) and EC cushion mesenchymal transition (EndMT) cells. (**C**) Top 50 upregulated genes (padj. < 0.05, log2FC > 0.5) are enriched for markers of cardiopharyngeal progenitors (CPM) and cells with epicardial (EP) or vascular endothelial (VEC) signatures, in line with a role of *Hand2* in coronary endothelial development^30^. (**D**) Left: UMAP projection of upregulated bulk RNA-seq DEGs from *H2*-URI^Δ/Δ^ vs. WT mouse embryonic hearts at E9.5 onto aggregated cardiac cell type clusters (n=20) from multiome analysis (Fig. 3). Color shades indicate over- (red) and under-represented (blue) expression of DEGs, in each cell. Right: violin plots showing the distribution of these values per cluster. (**E**) Comparison of average normalized expression of downregulated DEGs mapped to cardiac multiome clusters in *H2*-URI^Δ/Δ^ and WT control hearts. a/vCM, atrial/ventricular cardiomyocytes. AVC, atrioventricular canal. a/pSHF, anterior/posterior second heart field-derived cells. SM/FB, smooth muscle/fibroblast-like cells. EP, epicardial cells. EC, endocardial cells. VEC, vascular endothelial cells. EndMT, endocardial-to-mesenchymal transition/transformed cells of the cushions. PE, pulmonary endoderm. PA-endo/ecto, pharyngeal arch endo-/ectoderm. PaM/LPM, paraxial/lateral plate mesoderm. TP, thyroid progenitor-like cells. NCC, neural crest cells. HE, hematopoietic lineage cells. -e, early (E9.5 or E10.5). -l, late (E10.5 or E11.5). E, embryonic day.

## Supplementary Tables

**Table S1.**
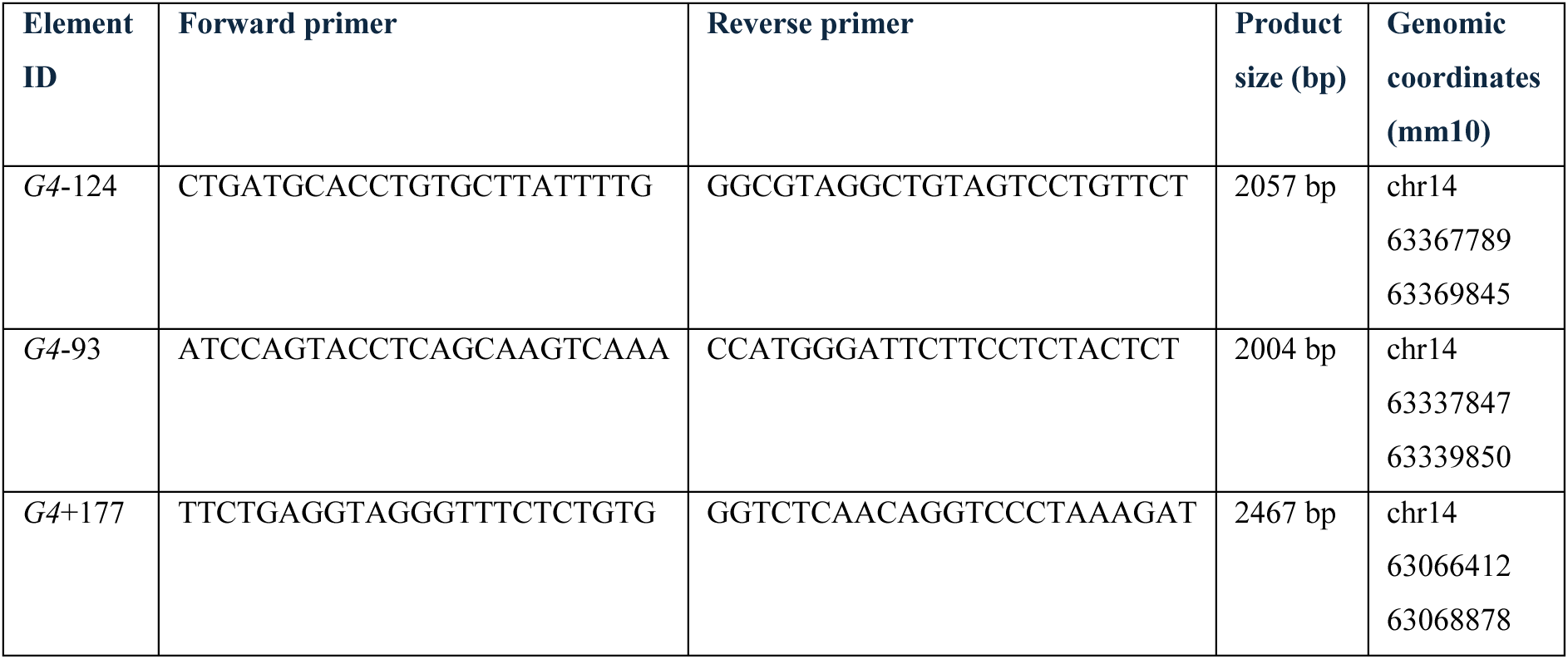
*Gata4* heart enhancer elements for site-directed reporter transgenesis. Evaluation of *Gata4* heart enhancer elements *G4*-124^61^, *G4*-93^62^ and *G4*+177 (new) using site-directed transgenesis at the *H11* safe-harbor locus^69,151^ (see Methods). Distance (in *kb*) from the *Hand2* TSS is indicated in brackets for each element (-, upstream; +, downstream).

**Table S2.**
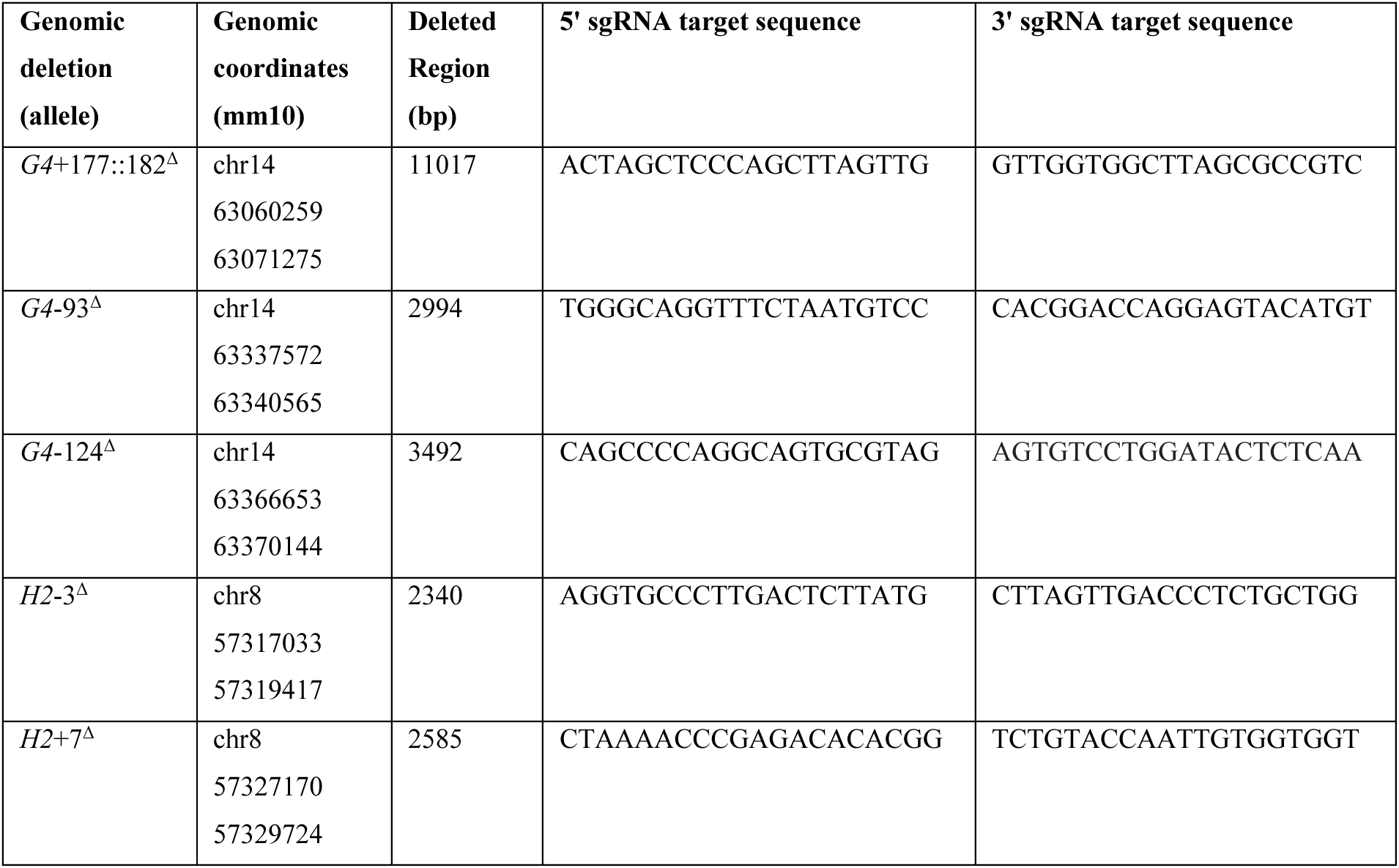
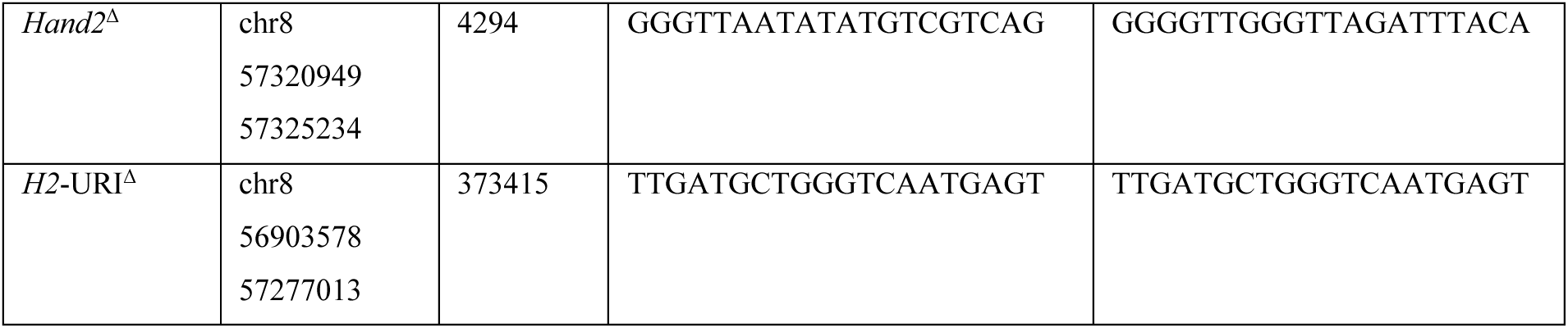
CRISPR deletions in mice. Genomic coordinates and sgRNA target sequences used for the generation of each founder mouse line via CRISPR/Cas9.

**Table S3.**
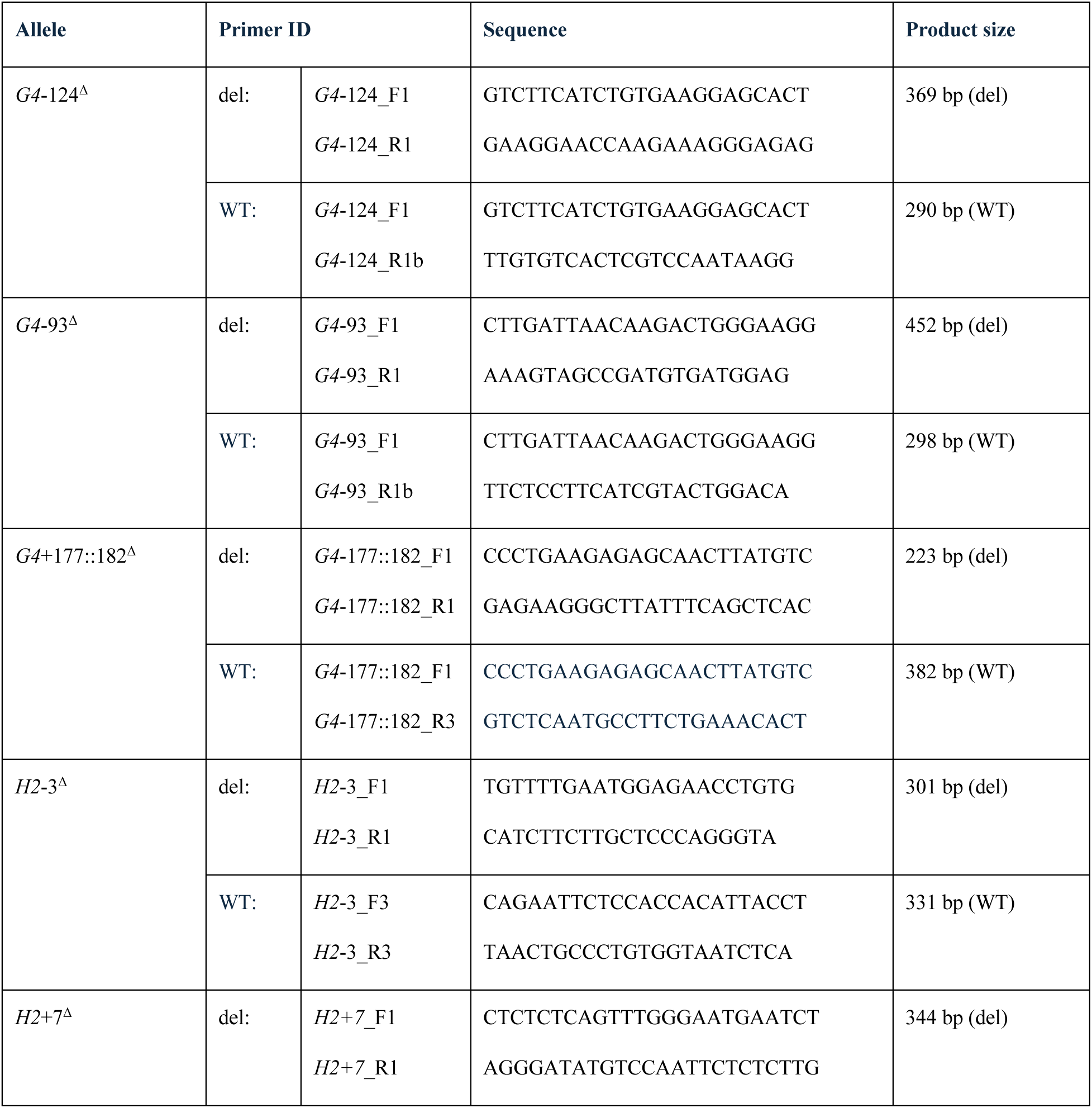
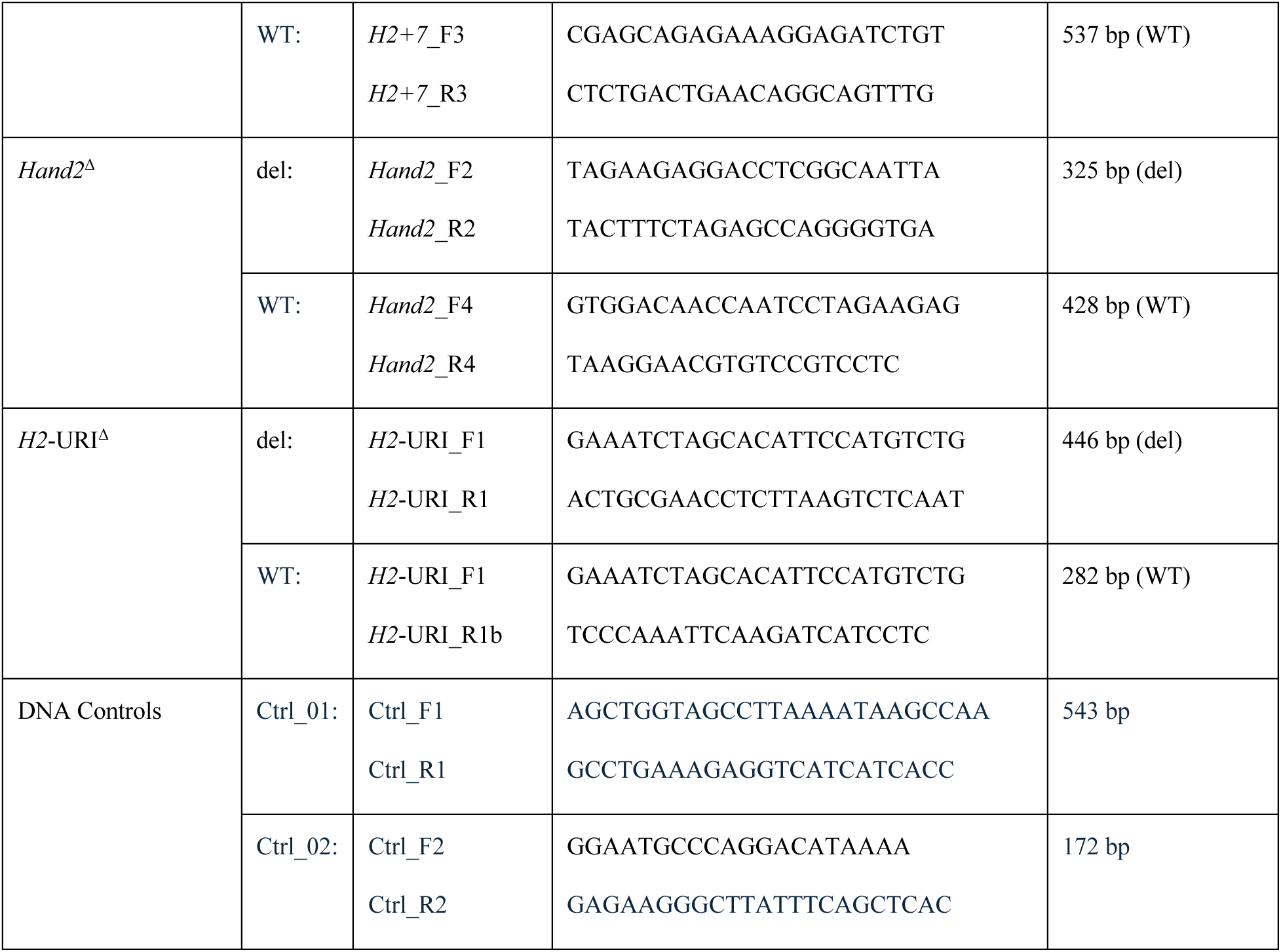
Primer pairs for screening and genotyping of mouse enhancer deletion alleles generated via CRISPR-Cas9. PCR genotyping strategies used to amplify genomic deletion (del) or wild-type (WT) allele-specific amplicons, along with the corresponding agarose gel electrophoresis results, are shown in **Figs. S2B**, **S4B** and **S11B**. A DNA control primer pair (Ctrl), designed to amplify an unrelated control region, was included in the reaction mixture together with the respective deletion (del) or wild-type (WT) primer pairs.

**Table S4.**
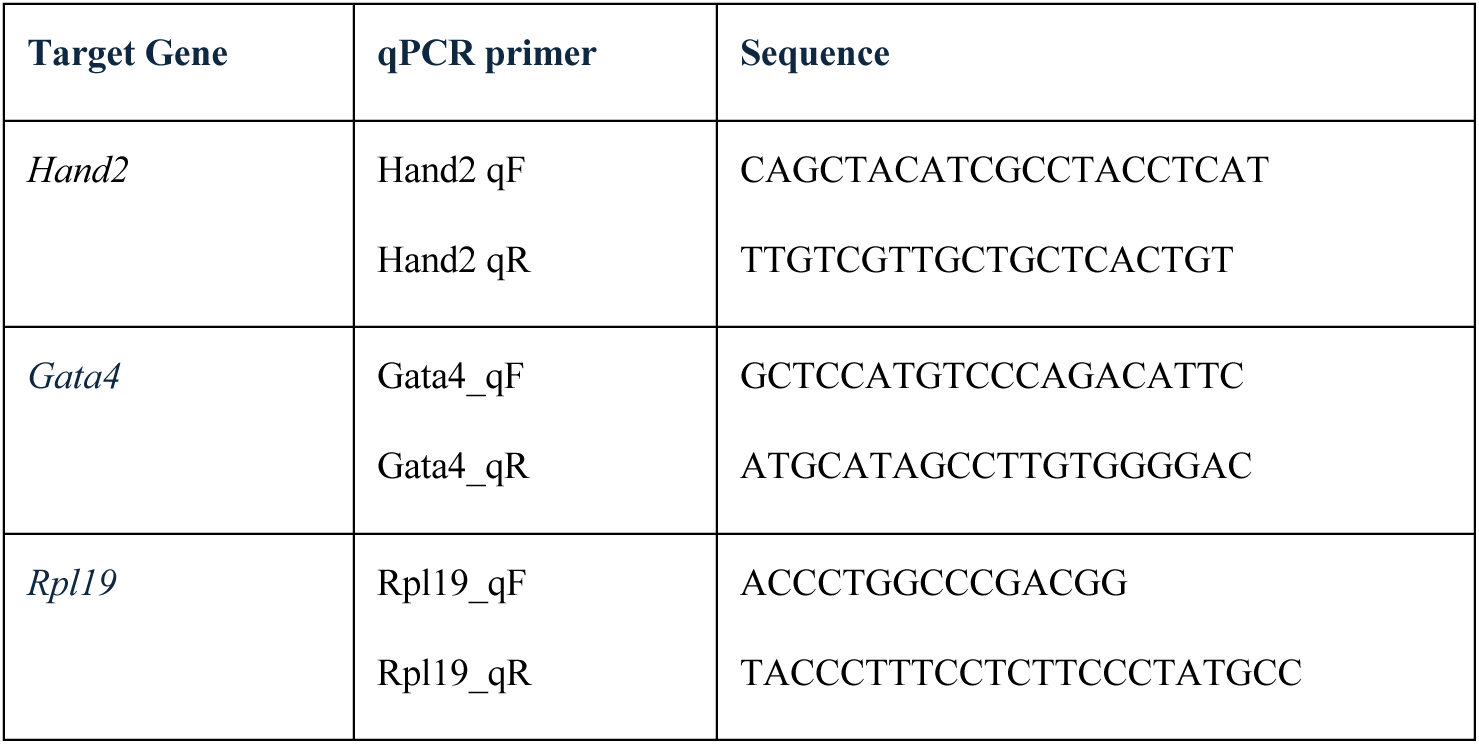
Primers used for qPCR analysis.

**Table S5.** Assessment of mouse heart enhancer-reporter activities in zebrafish. Number of transgenic F0 zebrafish embryos at 3-5 days post-fertilization (dpf) with tissue-specific fluorescent signals for the respective *Tol2*-integrated Enh-*mbG*-mCerulean; *cryaa*-Venus reporter constructs^80^ (**Fig. S3**). An integrated α-crystallin minimal promoter-driven reporter constitutively active in the eye was used to define transgene activity. IDs indicate the distance from the *Hand2* TSS (in kb; -, upstream; +, downstream).

| Enhancer ID | GFP+ |  | GFP- | Total |
| --- | --- | --- | --- | --- |
|  | Eye | Heart |  |  |
| G4-124 | 100 | 80 | 100 | 200 |
| G4-93 | 50 | 25 | 75 | 125 |
| G4+177 | 66 | 43 | 161 | 227 |
| G4+182 | 108 | 108 | 132 | 240 |

**Table S6 (provided as Excel file). Annotation of cell type identities to multiome clusters.** *<u>Sheet 1</u>*: List of published cardiac cell type markers based on differential gene expression in the midgestation mouse embryonic heart and adjacent populations^32,84^. *<u>Sheet 2</u>*: Cell-type cluster annotations for multiome pseudobulk clusters assigned based on marker genes listed in *Sheet1* and further validated with reference to published studies characterizing cardiac cell types^18,32,84–87,183^. The following parameters were used for cluster identification: WSNN, cluster0.75, min.pct.025, logfc.0.58 (see Methods). Myo, myocardial. Endo, endocardial/endothelial. NCC, neural crest cell lineage.

**Table S7 (provided as Excel file). Fraction of cells and differentially accessible peaks per multiome cluster.** Listed counts across developmental stages for each multiome lineage and cluster (*sheets 1-2*) (**Fig. S5E**), fractions of differentially accessible chromatin peaks for each multiome cluster across stages (*sheets 3-4*) (**Fig. S6B**), and fractions of differentially accessible peaks per cluster categorized by regulatory element identity (enhancer, promoter, or CTCF-only insulator) based on ChromHMM annotations (**Fig. S6C**).

**Table S8 (provided as Excel file). Chromatin accessibility of VISTA Enhancers across multiome clusters and cardiac lineages.** Table listing normalized average chromatin accessibility of n=3869 genomic elements previously validated for tissue-specific *in vivo* enhancer activity (VISTA Enhancer Browser)^94^ in multiome-defined cardiac lineages (*<u>sheet 1</u>*) and cell type clusters (*<u>sheet 2</u>*).

**Table S9 (provided as Excel file). Intersection of CHD-associated variants with cluster-specific accessible chromatin regions identified by multiome analysis.** *<u>Sheet 1</u>*: List of normalized overlap of GWAS CVD-associated top associations with cluster-specific peaks. *<u>Sheet 2</u>*: List of recently published CHD-associated variants and regions^39,50^ with and without overlap of multiome cluster-specific accessible chromatin signatures. *<u>Sheet 3</u>*: Overlap of CHD variants^39^ with experimental effect versus variants without effect, within cluster-specific peaks.

**Table S10 (provided as Excel file). List of TF motif enrichment across multiome clusters.** Table summarizing the enrichment values (–log10 of the p-value) for each TF motif (rows) within accessible chromatin regions (peaks) that exhibit significantly higher chromatin accessibility in each indicated cluster compared to all other clusters (columns).

**Table S11 (provided as Excel file). Metadata of predicted regulatory element maps across the *Hand2* and *Gata4* loci**. Table summarizing the datasets used to generate the panels in Figure S7. Data were retrieved from the Shiny App multiome interface (https://barozzilab.shinyapps.io/locus_heatmap/) and include cardiac cell type–specific chromatin accessibility, predicted multiomic correlation-based gene-enhancer interactions (“gene-enhancer links”), TF binding, evolutionary conservation and overlaps with predicted genomic features such as gene promoters, ChromHMM regulatory element state predictions (Ernst 2012) and previously validated enhancer elements (VISTA Enhancer Browser).

**Table S12.**
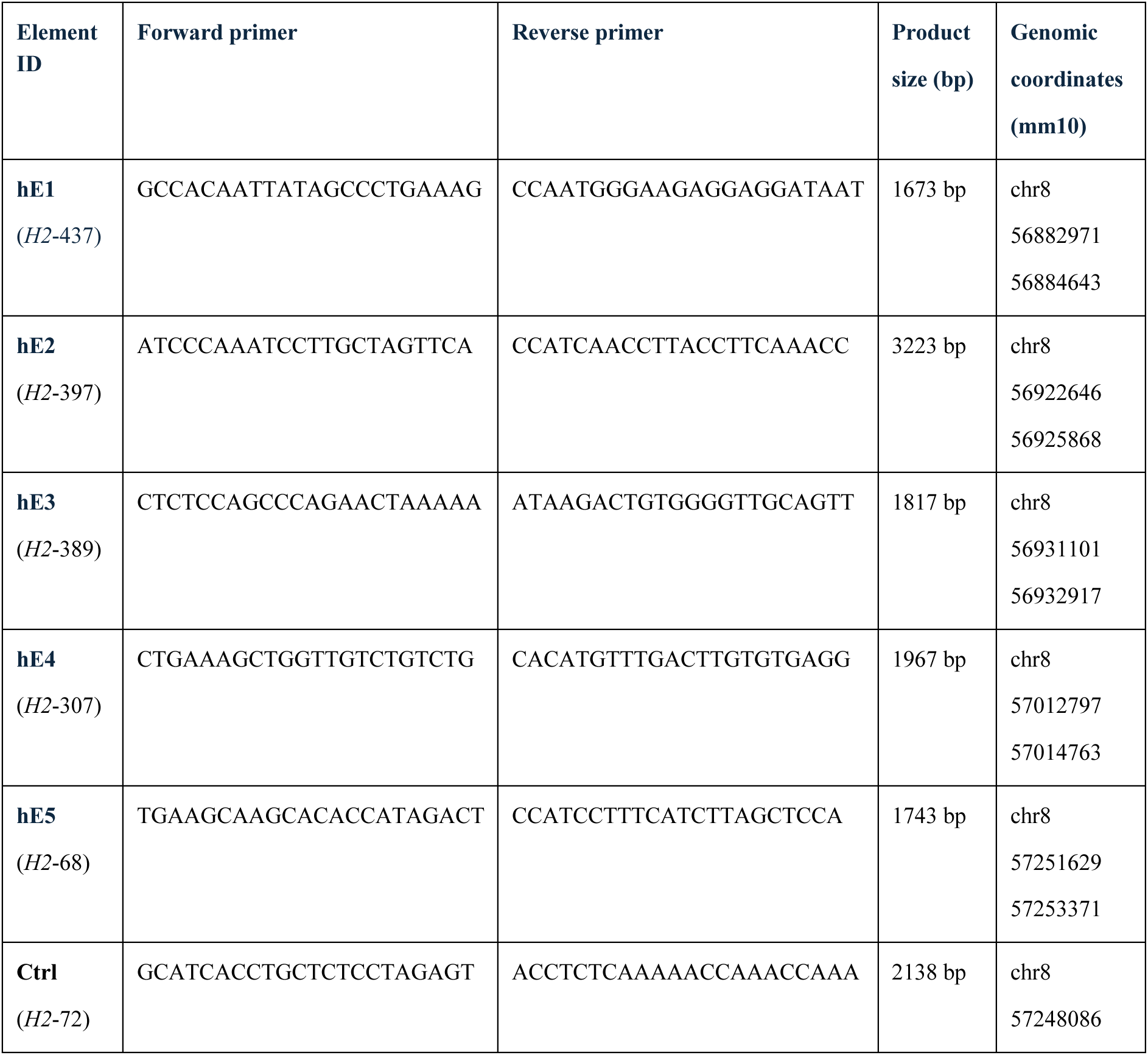
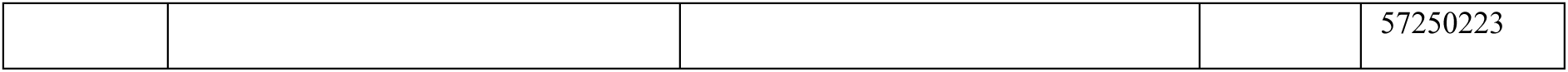
Predicted heart enhancers within the *H2*-URI validated using enSERT. Mouse genomic elements were PCR amplified and validated using enSERT site-directed transgenic enhancer-reporter analysis at the *H11* safe-harbor locus based on a targeting vector encoding a LacZ reporter unit with a minimal *Shh* promoter (enSERT)^69^. Element ID reflects distance (in *kb*) from the *Hand2* TSS (-, upstream; +, downstream).

**Table S13 (provided as Excel file). Reprocessed ATAC-seq and ChIP-seq datasets.** List of ENCODE datasets^70^ and previously published mouse ATAC-seq and ChIP-seq datasets from embryonic hearts^51,72,74,75,104^ re-processed as part of this study using uniform pipelines^53^ (see Methods). Resulting NarrowPeak (NP) and bigwig (BW) file types are indicated.

**Table S14.** Coordinates (mm10) of viewpoints used in normalized 4C analysis of the *Hand2* locus.

| Element ID | Genomic coordinates (mm10) |
| --- | --- |
| *Hand2* | chr8 57320983 57324633 |
| HT (*H2*-435) | chr8 56881203 56891203 |
| FL/MD (*H2*-685) | chr8 56631203 56641203 |
| hE1 (*H2*-437) | chr8 56882971 56884643 |
| hE2 (*H2*-397) | chr8 56922646 56925868 |
| hE3 (*H2*-389) | chr8 56931101 56932917 |
| hE4 (*H2*-307) | chr8 57012797 57014763 |
| hE5 (*H2*-68) | chr8 57251629 57253371 |

**Table S15.** Coordinates (mm10) of tissue-specific sub-TAD boundaries across the *Hand2* locus.

| Genomic Coordinates (mm10) | Tissue | Boundary ID |
| --- | --- | --- |
| chr8 56501203 56511203 | Heart | TAD-HT_01 |
| chr8 56871203 56881203 | Heart | TAD-HT_02 |
| chr8 57311203 57321203 | Heart | TAD-HT_03 |
| chr8 57541203 57551203 | Heart | TAD-HT_04 |
| chr8 57801203 57811203 | Heart | TAD-HT_05 |
| chr8 56491203 56501203 | Forelimb | TAD-FL_01 |
| chr8 57311203 57321203 | Forelimb | TAD-FL_02 |
| chr8 57531203 57541203 | Forelimb | TAD-FL_03 |
| chr8 57801203 57811203 | Forelimb | TAD-FL_04 |
| chr8 56491203 56501203 | Mandible | TAD-MD_01 |
| chr8 57301203 57311203 | Mandible | TAD-MD_02 |
| chr8 57531203 57541203 | Mandible | TAD-MD_03 |
| chr8 57801203 57811203 | Mandible | TAD-MD_04 |

**Table S16 (provided as Excel file). List of differentially expressed genes from RNA-seq comparison of WT and MUT (*H2*-URI^Δ/Δ^) embryos.** Table listing differentially expressed genes (padj > 0.05, log2FC > +/-0.5) (*sheets 1-3*) based on normalized read counts (*sheet 4*). Scaled read counts used for DEG matrix visualization via the Morpheus tool (https://software.broadinstitute.org/morpheus) are also listed (*sheet 5*).

## References

1. Morton, S. U., Quiat, D., Seidman, J. G. & Seidman, C. E. Genomic frontiers in congenital heart disease. Nat. Rev. Cardiol. 19, 26–42 (2022).

2. Bruneau, B. G. The developmental genetics of congenital heart disease. Nature 451, 943–948 (2008).

3. Jepson, B. M. et al. Pregnancy loss in major fetal congenital heart disease: incidence, risk factors and timing. Ultrasound Obstet. Gynecol. 62, 75–87 (2023).

4. Yasuhara, J. & Garg, V. Genetics of congenital heart disease: a narrative review of recent advances and clinical implications. Transl Pediatr 10, 2366–2386 (2021).

5. Olson, E. N. Gene regulatory networks in the evolution and development of the heart. Science 313, 1922–1927 (2006).

6. McCulley, D. J. & Black, B. L. Chapter nine - Transcription Factor Pathways and Congenital Heart Disease. in Current Topics in Developmental Biology (ed. Bruneau, B. G.) vol. 100 253–277 (Academic Press, 2012).

7. Waardenberg, A. J., Ramialison, M., Bouveret, R. & Harvey, R. P. Genetic networks governing heart development. Cold Spring Harb. Perspect. Med. 4, a013839 (2014).

8. Tolkin, T. & Christiaen, L. Development and evolution of the ascidian cardiogenic mesoderm. Curr. Top. Dev. Biol. 100, 107–142 (2012).

9. Meilhac, S. M. & Buckingham, M. E. The deployment of cell lineages that form the mammalian heart. Nat. Rev. Cardiol. 15, 705–724 (2018).

10. Luna-Zurita, L. et al. Complex Interdependence Regulates Heterotypic Transcription Factor Distribution and Coordinates Cardiogenesis. Cell 164, 999–1014 (2016).

11. Muncie-Vasic, J. M. et al. MEF2C controls segment-specific gene regulatory networks that direct heart tube morphogenesis. Genes Dev. (2025) doi:10.1101/gad.352889.125.

12. Gonzalez-Teran, B. et al. Transcription factor protein interactomes reveal genetic determinants in heart disease. Cell (2022) doi:10.1016/j.cell.2022.01.021.

13. Kathiriya, I. S., Nora, E. P. & Bruneau, B. G. Investigating the transcriptional control of cardiovascular development. Circ. Res. 116, 700–714 (2015).

14. George, R. M. & Firulli, A. B. Hand Factors in Cardiac Development. Anat. Rec. 302, 101–107 (2019).

15. MacGrogan, D., Münch, J. & de la Pompa, J. L. Notch and interacting signalling pathways in cardiac development, disease, and regeneration. Nat. Rev. Cardiol. 15, 685–704 (2018).

16. Buckingham, M., Meilhac, S. & Zaffran, S. Building the mammalian heart from two sources of myocardial cells. Nat. Rev. Genet. 6, 826–835 (2005).

17. Lescroart, F. et al. Defining the earliest step of cardiovascular lineage segregation by single-cell RNA-seq. Science 359, 1177–1181 (2018).

18. De Bono, C. et al. Single-cell transcriptomics uncovers a non-autonomous Tbx1-dependent genetic program controlling cardiac neural crest cell development. Nat. Commun. 14, 1551 (2023).

19. Quijada, P., Trembley, M. A. & Small, E. M. The role of the epicardium during heart development and repair. Circ. Res. 126, 377–394 (2020).

20. Kelly, R. G. The heart field transcriptional landscape at single-cell resolution. Dev. Cell 58, 257–266 (2023).

21. Lyons, I. et al. Myogenic and morphogenetic defects in the heart tubes of murine embryos lacking the homeo box gene Nkx2-5. Genes Dev. 9, 1654–1666 (1995).

22. Molkentin, J. D., Lin, Q., Duncan, S. A. & Olson, E. N. Requirement of the transcription factor GATA4 for heart tube formation and ventral morphogenesis. Genes Dev. 11, 1061–1072 (1997).

23. Lin, Q., Schwarz, J., Bucana, C. & Olson, E. N. Control of mouse cardiac morphogenesis and myogenesis by transcription factor MEF2C. Science 276, 1404–1407 (1997).

24. Bruneau, B. G. et al. A murine model of Holt-Oram syndrome defines roles of the T-box transcription factor Tbx5 in cardiogenesis and disease. Cell 106, 709–721 (2001).

25. Srivastava, D. et al. Regulation of cardiac mesodermal and neural crest development by the bHLH transcription factor, dHAND. Nat. Genet. 16, 154–160 (1997).

26. Lindsay, E. et al. Tbx1 haploinsufficiency in the DiGeorge syndrome region causes aortic arch defects in mice. Nature 410, 97–101 (2001).

27. Zeisberg, E. M. et al. Morphogenesis of the right ventricle requires myocardial expression of Gata4. J. Clin. Invest. 115, 1522–1531 (2005).

28. Tsuchihashi, T. et al. Hand2 function in second heart field progenitors is essential for cardiogenesis. Dev. Biol. 351, 62–69 (2011).

29. Rivera-Feliciano, J. et al. Development of heart valves requires Gata4 expression in endothelial-derived cells. Development 133, 3607–3618 (2006).

30. VanDusen, N. J. et al. Hand2 is an essential regulator for two Notch-dependent functions within the embryonic endocardium. Cell Rep. 9, 2071–2083 (2014).

31. Liu, J. et al. Gata4 regulates hedgehog signaling and Gata6 expression for outflow tract development. PLoS Genet. 15, e1007711 (2019).

32. de Soysa, T. Y. et al. Single-cell analysis of cardiogenesis reveals basis for organ-level developmental defects. Nature 572, 120–124 (2019).

33. Yuan, X., Scott, I. C. & Wilson, M. D. Heart Enhancers: Development and Disease Control at a Distance. Front. Genet. 12, 642975 (2021).

34. George, R. M. et al. Single cell evaluation of endocardial Hand2 gene regulatory networks reveals HAND2-dependent pathways that impact cardiac morphogenesis. Development 150, (2023).

35. Kathiriya, I. S., et al. Reduced TBX5 dosage undermines developmental control of atrial cardiomyocyte identity in a model of human atrial disease. bioRxivorg (2025) doi:10.1101/2025.08.16.669546.

36. Pu, W. T., Ishiwata, T., Juraszek, A. L., Ma, Q. & Izumo, S. GATA4 is a dosage-sensitive regulator of cardiac morphogenesis. Dev. Biol. 275, 235–244 (2004).

37. Anene-Nzelu, C. G., Lee, M. C. J., Tan, W. L. W., Dashi, A. & Foo, R. S. Y. Genomic enhancers in cardiac development and disease. Nat. Rev. Cardiol. (2021) doi:10.1038/s41569-021-00597-2.

38. Blue, G. M. et al. Advances in the Genetics of Congenital Heart Disease: A Clinician’s Guide. J. Am. Coll. Cardiol. 69, 859–870 (2017).

39. Xiao, F. et al. Functional dissection of human cardiac enhancers and noncoding de novo variants in congenital heart disease. Nat. Genet. (2024) doi:10.1038/s41588-024-01669-y.

40. Dickel, D. E. et al. Genome-wide compendium and functional assessment of in vivo heart enhancers. Nat. Commun. 7, 12923 (2016).

41. Armendariz, D. A. et al. CHD-associated enhancers shape human cardiomyocyte lineage commitment. Elife 12, (2023).

42. Vincentz, J. W. et al. Variation in a Left Ventricle-Specific Hand1 Enhancer Impairs GATA Transcription Factor Binding and Disrupts Conduction System Development and Function. Circ. Res. 125, 575–589 (2019).

43. van Eif, V. W. W. et al. Genome-Wide Analysis Identifies an Essential Human TBX3 Pacemaker Enhancer. Circ. Res. 127, 1522–1535 (2020).

44. Smemo, S. et al. Regulatory variation in a TBX5 enhancer leads to isolated congenital heart disease. Hum. Mol. Genet. 21, 3255–3263 (2012).

45. Lupiáñez, D. G. et al. Disruptions of topological chromatin domains cause pathogenic rewiring of gene-enhancer interactions. Cell 161, 1012–1025 (2015).

46. Spielmann, M., Lupiáñez, D. G. & Mundlos, S. Structural variation in the 3D genome. Nat. Rev. Genet. 19, 453–467 (2018).

47. Baudic, M. et al. TAD boundary deletion causes PITX2-related cardiac electrical and structural defects. Nat. Commun. 15, 3380 (2024).

48. Gorkin, D. U. et al. An atlas of dynamic chromatin landscapes in mouse fetal development. Nature 583, 744–751 (2020).

49. Hocker, J. D. et al. Cardiac cell type-specific gene regulatory programs and disease risk association. Sci. Adv. 7, eabf1444 (2021).

50. Ameen, M. et al. Integrative single-cell analysis of cardiogenesis identifies developmental trajectories and non-coding mutations in congenital heart disease. Cell 185, 4937–4953.e23 (2022).

51. Zhou, P. et al. GATA4 Regulates Developing Endocardium Through Interaction With ETS1. Circ. Res. 131, e152–e168 (2022).

52. Osterwalder, M. et al. Enhancer redundancy provides phenotypic robustness in mammalian development. Nature 554, 239–243 (2018).

53. Abassah-Oppong, S. et al. A gene desert required for regulatory control of pleiotropic Shox2 expression and embryonic survival. Nat. Commun. 15, (2024).

54. Man, J. C. K. et al. An enhancer cluster controls gene activity and topology of the SCN5A-SCN10A locus in vivo. Nat. Commun. 10, 4943 (2019).

55. van den Boogaard, M., et al. Identification and Characterization of a Transcribed Distal Enhancer Involved in Cardiac Kcnh2 Regulation. Cell Rep. 28, 2704–2714.e5 (2019).

56. Galang, G. et al. ATAC-Seq Reveals an Isl1 Enhancer That Regulates Sinoatrial Node Development and Function. Circ. Res. 127, 1502–1518 (2020).

57. Yamaguchi, N. et al. An Anterior Second Heart Field Enhancer Regulates the Gene Regulatory Network of the Cardiac Outflow Tract. Circulation 148, 1705–1722 (2023).

58. Jindal, G. A. et al. Single-nucleotide variants within heart enhancers increase binding affinity and disrupt heart development. Dev. Cell (2023) doi:10.1016/j.devcel.2023.09.005.

59. Morikawa, Y. & Cserjesi, P. Cardiac neural crest expression of Hand2 regulates outflow and second heart field development. Circ. Res. 103, 1422–1429 (2008).

60. McFadden, D. G. et al. A GATA-dependent right ventricular enhancer controls dHAND transcription in the developing heart. Development 127, 5331–5341 (2000).

61. Blow, M. J. et al. ChIP-Seq identification of weakly conserved heart enhancers. Nat. Genet. 42, 806–810 (2010).

62. Schachterle, W., Rojas, A., Xu, S.-M. & Black, B. L. ETS-dependent regulation of a distal Gata4 cardiac enhancer. Dev. Biol. 361, 439–449 (2012).

63. Vincentz, J. W. et al. HAND transcription factors cooperatively specify the aorta and pulmonary trunk. Dev. Biol. 476, 1–10 (2021).

64. Cannavò, E. et al. Shadow Enhancers Are Pervasive Features of Developmental Regulatory Networks. Curr. Biol. 26, 38–51 (2016).

65. Kvon, E. Z., Waymack, R., Gad, M. & Wunderlich, Z. Enhancer redundancy in development and disease. Nat. Rev. Genet. 22, 324–336 (2021).

66. Tamura, M., Amano, T. & Shiroishi, T. The Hand2 gene dosage effect in developmental defects and human congenital disorders. Curr. Top. Dev. Biol. 110, 129–152 (2014).

67. Xin, M., Olson, E. N. & Bassel-Duby, R. Mending broken hearts: cardiac development as a basis for adult heart regeneration and repair. Nat. Rev. Mol. Cell Biol. 14, 529–541 (2013).

68. Phan, M. H. Q. et al. Conservation of regulatory elements with highly diverged sequences across large evolutionary distances. Nat. Genet. 57, 1524–1534 (2025).

69. Kvon, E. Z. et al. Comprehensive In Vivo Interrogation Reveals Phenotypic Impact of Human Enhancer Variants. Cell 180, 1262–1271.e15 (2020).

70. ENCODE Project Consortium et al. Expanded encyclopaedias of DNA elements in the human and mouse genomes. Nature 583, 699–710 (2020).

71. Osterwalder, M. et al. HAND2 targets define a network of transcriptional regulators that compartmentalize the early limb bud mesenchyme. Dev. Cell 31, 345–357 (2014).

72. Laurent, F. et al. HAND2 Target Gene Regulatory Networks Control Atrioventricular Canal and Cardiac Valve Development. Cell Rep. 19, 1602–1613 (2017).

73. Chen, S., Lee, B., Lee, A. Y.-F., Modzelewski, A. J. & He, L. Highly Efficient Mouse Genome Editing by CRISPR Ribonucleoprotein Electroporation of Zygotes. J. Biol. Chem. 291, 14457–14467 (2016).

74. He, A. et al. Dynamic GATA4 enhancers shape the chromatin landscape central to heart development and disease. Nat. Commun. 5, 4907 (2014).

75. Akerberg, B. N. et al. A reference map of murine cardiac transcription factor chromatin occupancy identifies dynamic and conserved enhancers. Nat. Commun. 10, 4907 (2019).

76. Will, A. J. et al. Composition and dosage of a multipartite enhancer cluster control developmental expression of Ihh (Indian hedgehog). Nat. Genet. 49, 1539–1545 (2017).

77. Montavon, T. et al. A regulatory archipelago controls Hox genes transcription in digits. Cell 147, 1132–1145 (2011).

78. Malkmus, J. et al. Spatial regulation by multiple Gremlin1 enhancers provides digit development with cis-regulatory robustness and evolutionary plasticity. Nat. Commun. 12, 5557 (2021).

79. Wong, E. S. et al. Deep conservation of the enhancer regulatory code in animals. Science 370, (2020).

80. Kemmler, C. L. et al. Next-generation plasmids for transgenesis in zebrafish and beyond. Development 150, (2023).

81. Kemmler, C. L. et al. Conserved enhancers control notochord expression of vertebrate Brachyury. Nat. Commun. 14, 6594 (2023).

82. Simon, C. S. et al. A Gata4 nuclear GFP transcriptional reporter to study endoderm and cardiac development in the mouse. Biol. Open 7, (2018).

83. Galli, A. et al. Distinct roles of Hand2 in initiating polarity and posterior Shh expression during the onset of mouse limb bud development. PLoS Genet. 6, e1000901 (2010).

84. Rowton, M. et al. Hedgehog signaling activates a mammalian heterochronic gene regulatory network controlling differentiation timing across lineages. Dev. Cell (2022) doi:10.1016/j.devcel.2022.08.009.

85. Lotto, J. et al. Cell diversity and plasticity during atrioventricular heart valve EMTs. Nat. Commun. 14, 5567 (2023).

86. Farhat, B. et al. Understanding the cell fate and behavior of progenitors at the origin of the mouse cardiac mitral valve. Dev. Cell 59, 339–350.e4 (2024).

87. Harel, I. et al. Pharyngeal mesoderm regulatory network controls cardiac and head muscle morphogenesis. Proc. Natl. Acad. Sci. U. S. A. 109, 18839–18844 (2012).

88. Liu, X. et al. Single-Cell RNA-Seq of the Developing Cardiac Outflow Tract Reveals Convergent Development of the Vascular Smooth Muscle Cells. Cell Rep. 28, 1346–1361.e4 (2019).

89. Xiao, Y. et al. Hippo signaling plays an essential role in cell state transitions during cardiac fibroblast development. Dev. Cell 45, 153–169.e6 (2018).

90. Nomaru, H. et al. Single cell multi-omic analysis identifies a Tbx1-dependent multilineage primed population in murine cardiopharyngeal mesoderm. Nat. Commun. 12, 6645 (2021).

91. Kolander, K. D., Holtz, M. L., Cossette, S. M., Duncan, S. A. & Misra, R. P. Epicardial GATA factors regulate early coronary vascular plexus formation. Dev. Biol. 386, 204–215 (2014).

92. Barnes, R. M. et al. Hand2 loss-of-function in Hand1-expressing cells reveals distinct roles in epicardial and coronary vessel development. Circ. Res. 108, 940–949 (2011).

93. Pollex, T. et al. Enhancer-promoter interactions become more instructive in the transition from cell-fate specification to tissue differentiation. Nat. Genet. (2024) doi:10.1038/s41588-024-01678-x.

94. Kosicki, M. et al. VISTA Enhancer browser: an updated database of tissue-specific developmental enhancers. Nucleic Acids Res. 53, D324–D330 (2025).

95. Richter, F. et al. Genomic analyses implicate noncoding de novo variants in congenital heart disease. Nat. Genet. 52, 769–777 (2020).

96. Fernandez-Perez, A. et al. Hand2 Selectively Reorganizes Chromatin Accessibility to Induce Pacemaker-like Transcriptional Reprogramming. Cell Rep. 27, 2354–2369.e7 (2019).

97. Ernst, J. & Kellis, M. ChromHMM: automating chromatin-state discovery and characterization. Nat. Methods 9, 215–216 (2012).

98. Anderson, K. M. et al. Transcription of the non-coding RNA upperhand controls Hand2 expression and heart development. Nature 539, 433–436 (2016).

99. Han, X. et al. The lncRNA Hand2os1/Uph locus orchestrates heart development through regulation of precise expression of Hand2. Development 146, (2019).

100. Monti, R. et al. Limb-Enhancer Genie: An accessible resource of accurate enhancer predictions in the developing limb. PLoS Comput. Biol. 13, e1005720 (2017).

101. Losa, M. et al. A spatio-temporally constrained gene regulatory network directed by PBX1/2 acquires limb patterning specificity via HAND2. Nat. Commun. 14, 3993 (2023).

102. Ferguson, C. A., Firulli, B. A., Zoia, M., Osterwalder, M. & Firulli, A. B. Identification and characterization of Hand2 upstream genomic enhancers active in developing stomach and limbs. Dev. Dyn. (2023) doi:10.1002/dvdy.646.

103. Pampari, A. et al. ChromBPNet: bias factorized, base-resolution deep learning models of chromatin accessibility reveal cis-regulatory sequence syntax, transcription factor footprints and regulatory variants. bioRxivorg (2025) doi:10.1101/2024.12.25.630221.

104. Cao, Y. et al. In Vivo Dissection of Chamber-Selective Enhancers Reveals Estrogen-Related Receptor as a Regulator of Ventricular Cardiomyocyte Identity. Circulation (2023) doi:10.1161/CIRCULATIONAHA.122.061955.

105. Franke, M. et al. Formation of new chromatin domains determines pathogenicity of genomic duplications. Nature 538, 265–269 (2016).

106. Yanagisawa, H., Clouthier, D. E., Richardson, J. A., Charité, J. & Olson, E. N. Targeted deletion of a branchial arch-specific enhancer reveals a role of dHAND in craniofacial development. Development 130, 1069–1078 (2003).

107. Dekker, J. et al. Spatial and temporal organization of the genome: Current state and future aims of the 4D nucleome project. Mol. Cell (2023) doi:10.1016/j.molcel.2023.06.018.

108. Nicodemi, M. & Prisco, A. Thermodynamic pathways to genome spatial organization in the cell nucleus. Biophys. J. 96, 2168–2177 (2009).

109. Chiariello, A. M., Annunziatella, C., Bianco, S., Esposito, A. & Nicodemi, M. Polymer physics of chromosome large-scale 3D organisation. Sci. Rep. 6, 29775 (2016).

110. Conte, M. et al. Polymer physics indicates chromatin folding variability across single-cells results from state degeneracy in phase separation. Nat. Commun. 11, 3289 (2020).

111. Kragesteen, B. K. et al. Dynamic 3D chromatin architecture contributes to enhancer specificity and limb morphogenesis. Nat. Genet. 50, 1463–1473 (2018).

112. Esposito, A. et al. Polymer physics reveals a combinatorial code linking 3D chromatin architecture to 1D chromatin states. Cell Rep. 38, 110601 (2022).

113. Bianco, S. et al. Polymer physics predicts the effects of structural variants on chromatin architecture. Nat. Genet. 50, 662–667 (2018).

114. Laugwitz, K.-L., Moretti, A., Caron, L., Nakano, A. & Chien, K. R. Islet1 cardiovascular progenitors: a single source for heart lineages? Development 135, 193–205 (2008).

115. Zhou, P., He, A. & Pu, W. T. Regulation of GATA4 transcriptional activity in cardiovascular development and disease. Curr. Top. Dev. Biol. 100, 143–169 (2012).

116. Grego-Bessa, J. et al. Notch signaling is essential for ventricular chamber development. Dev. Cell 12, 415–429 (2007).

117. Shelton, E. L. & Yutzey, K. E. Twist1 function in endocardial cushion cell proliferation, migration, and differentiation during heart valve development. Dev. Biol. 317, 282–295 (2008).

118. Camenisch, T. D. et al. Disruption of hyaluronan synthase-2 abrogates normal cardiac morphogenesis and hyaluronan-mediated transformation of epithelium to mesenchyme. J. Clin. Invest. 106, 349–360 (2000).

119. Conway, S. & Molkentin, J. Periostin as a heterofunctional regulator of cardiac development and disease. Curr. Genomics 9, 548–555 (2008).

120. Delgado, I. et al. Control of mouse limb initiation and antero-posterior patterning by Meis transcription factors. Nat. Commun. 12, 1–13 (2021).

121. Dickel, D. E. et al. Ultraconserved Enhancers Are Required for Normal Development. Cell 172, 491–499.e15 (2018).

122. Haberle, V. et al. Transcriptional cofactors display specificity for distinct types of core promoters. Nature 570, 122–126 (2019).

123. Hörnblad, A., Bastide, S., Langenfeld, K., Langa, F. & Spitz, F. Dissection of the Fgf8 regulatory landscape by in vivo CRISPR-editing reveals extensive intra- and inter-enhancer redundancy. Nat. Commun. 12, 439 (2021).

124. Cunningham, T. J., Lancman, J. J., Berenguer, M., Dong, P. D. S. & Duester, G. Genomic Knockout of Two Presumed Forelimb Tbx5 Enhancers Reveals They Are Nonessential for Limb Development. Cell Rep. 23, 3146–3151 (2018).

125. Hay, D. et al. Genetic dissection of the α-globin super-enhancer in vivo. Nat. Genet. 48, 895–903 (2016).

126. Long, H. K., Prescott, S. L. & Wysocka, J. Ever-Changing Landscapes: Transcriptional Enhancers in Development and Evolution. Cell 167, 1170–1187 (2016).

127. Loubiere, V., de Almeida, B. P., Pagani, M. & Stark, A. Developmental and housekeeping transcriptional programs display distinct modes of enhancer-enhancer cooperativity in Drosophila. Nat. Commun. 15, 8584 (2024).

128. Wang, X. & Goldstein, D. B. Enhancer domains predict gene pathogenicity and inform gene discovery in complex disease. Am. J. Hum. Genet. 106, 215–233 (2020).

129. Perry, M. W., Boettiger, A. N., Bothma, J. P. & Levine, M. Shadow enhancers foster robustness of Drosophila gastrulation. Curr. Biol. 20, 1562–1567 (2010).

130. Frankel, N. et al. Phenotypic robustness conferred by apparently redundant transcriptional enhancers. Nature 466, 490–493 (2010).

131. Hong, J.-W., Hendrix, D. A. & Levine, M. S. BREVIA Shadow Enhancers as a Source of Evolutionary Novelty. Science 321, (2008).

132. Antosova, B. et al. The gene regulatory network of lens induction is wired through Meis-dependent shadow enhancers of Pax6. PLoS Genet. 12, e1006441 (2016).

133. Sagai, T. et al. SHH signaling directed by two oral epithelium-specific enhancers controls tooth and oral development. Sci. Rep. 7, 13004 (2017).

134. Munshi, N. V. et al. Cx30.2 enhancer analysis identifies Gata4 as a novel regulator of atrioventricular delay. Development 136, 2665–2674 (2009).

135. Stefanovic, S. et al. GATA-dependent regulatory switches establish atrioventricular canal specificity during heart development. Nat. Commun. 5, 3680 (2014).

136. Ranade, S. S. et al. Single Cell Epigenetics Reveal Cell-Cell Communication Networks in Normal and Abnormal Cardiac Morphogenesis. bioRxiv 2022.07.25.501458 (2022) doi:10.1101/2022.07.25.501458.

137. Holman, A. R. et al. Single-cell multi-modal integrative analyses highlight functional dynamic gene regulatory networks directing human cardiac development. Cell Genom. 100680 (2024).

138. van Duijvenboden, K., de Boer, B. A., Capon, N., Ruijter, J. M. & Christoffels, V. M. EMERGE: a flexible modelling framework to predict genomic regulatory elements from genomic signatures. Nucleic Acids Res. 44, e42 (2016).

139. Corces, M. R. et al. Single-cell epigenomic analyses implicate candidate causal variants at inherited risk loci for Alzheimer’s and Parkinson’s diseases. Nat. Genet. 52, 1158–1168 (2020).

140. Zhu, K. et al. Multi-omic profiling of the developing human cerebral cortex at the single-cell level. Sci. Adv. 9, eadg3754 (2023).

141. Paliou, C. et al. Preformed chromatin topology assists transcriptional robustness of Shh during limb development. Proc. Natl. Acad. Sci. U. S. A. 116, 12390–12399 (2019).

142. Furlong, E. E. M. & Levine, M. Developmental enhancers and chromosome topology. Science 361, 1341–1345 (2018).

143. McGurk, K. A. et al. Genetic and phenotypic architecture of human myocardial trabeculation. *Nat*. Cardiovasc. Res. (2024) doi:10.1038/s44161-024-00564-3.

144. Spurrell, C. H. et al. Genome-wide fetalization of enhancer architecture in heart disease. Cell Rep. 40, 111400 (2022).

145. Ritter, N. et al. The lncRNA Locus Handsdown Regulates Cardiac Gene Programs and Is Essential for Early Mouse Development. Dev. Cell 50, 644–657.e8 (2019).

146. Amemiya, C. T. et al. The African coelacanth genome provides insights into tetrapod evolution. Nature 496, 311–316 (2013).

147. Yuan, X. et al. Heart enhancers with deeply conserved regulatory activity are established early in zebrafish development. Nat. Commun. 9, 4977 (2018).

148. Jindal, G. A. & Farley, E. K. Enhancer grammar in development, evolution, and disease: dependencies and interplay. Dev. Cell 56, 575–587 (2021).

149. de Almeida, B. P. et al. Targeted design of synthetic enhancers for selected tissues in the Drosophila embryo. Nature (2023) doi:10.1038/s41586-023-06905-9.

150. Taskiran, I. I. et al. Cell-type-directed design of synthetic enhancers. Nature 626, 212–220 (2024).

151. Osterwalder, M. et al. Characterization of Mammalian In Vivo Enhancers Using Mouse Transgenesis and CRISPR Genome Editing. Methods Mol. Biol. 2403, 147–186 (2022).

152. Concordet, J.-P. & Haeussler, M. CRISPOR: intuitive guide selection for CRISPR/Cas9 genome editing experiments and screens. Nucleic Acids Res. 46, W242–W245 (2018).

153. Tissières, V. et al. Gene Regulatory and Expression Differences between Mouse and Pig Limb Buds Provide Insights into the Evolutionary Emergence of Artiodactyl Traits. Cell Rep. 31, 107490 (2020).

154. Stuart, T. et al. Comprehensive Integration of Single-Cell Data. Cell 177, 1888–1902.e21 (2019).

155. Granja, J. M. et al. ArchR is a scalable software package for integrative single-cell chromatin accessibility analysis. Nat. Genet. 53, 403–411 (2021).

156. Stuart, T., Srivastava, A., Madad, S., Lareau, C. A. & Satija, R. Single-cell chromatin state analysis with Signac. Nat. Methods 18, 1333–1341 (2021).

157. Zhang, Y. et al. Model-based analysis of ChIP-Seq (MACS). Genome Biol. 9, R137 (2008).

158. Ernst, J. & Kellis, M. Chromatin-state discovery and genome annotation with ChromHMM. Nat. Protoc. 12, 2478–2492 (2017).

159. Cerezo, M. et al. The NHGRI-EBI GWAS Catalog: standards for reusability, sustainability and diversity. Nucleic Acids Res. 53, D998–D1005 (2025).

160. Quinlan, A. R. & Hall, I. M. BEDTools: a flexible suite of utilities for comparing genomic features. Bioinformatics 26, 841–842 (2010).

161. Langmead, B. & Salzberg, S. L. Fast gapped-read alignment with Bowtie 2. Nat. Methods 9, 357–359 (2012).

162. Li, H. et al. The Sequence Alignment/Map format and SAMtools. Bioinformatics 25, 2078–2079 (2009).

163. Feng, J., Liu, T., Qin, B., Zhang, Y. & Liu, X. S. Identifying ChIP-seq enrichment using MACS. Nat. Protoc. 7, 1728–1740 (2012).

164. Rouco, R. et al. Cell-specific alterations in Pitx1 regulatory landscape activation caused by the loss of a single enhancer. Nat. Commun. 12, 7235 (2021).

165. Wingett, S. et al. HiCUP: pipeline for mapping and processing Hi-C data. F1000Res. 4, 1310 (2015).

166. Durand, N. C. et al. Juicer Provides a One-Click System for Analyzing Loop-Resolution Hi-C Experiments. Cell Syst 3, 95–98 (2016).

167. Abdennur, N. et al. Cooltools: Enabling high-resolution Hi-C analysis in Python. PLoS Comput Biol 20, e1012067 (2024).

168. Barbieri, M. et al. Complexity of chromatin folding is captured by the strings and binders switch model. Proc. Natl. Acad. Sci. U. S. A. 109, 16173–16178 (2012).

169. Knight, P. A. & Ruiz, D. A fast algorithm for matrix balancing. IMA J. Numer. Anal. 33, 1029–1047 (2013).

170. Brackley, C. A., Taylor, S., Papantonis, A., Cook, P. R. & Marenduzzo, D. Nonspecific bridging-induced attraction drives clustering of DNA-binding proteins and genome organization. Proc. Natl. Acad. Sci. U. S. A. 110, E3605–11 (2013).

171. Thompson, A. P. et al. LAMMPS - a flexible simulation tool for particle-based materials modeling at the atomic, meso, and continuum scales. Comput. Phys. Commun. 271, 108171 (2022).

172. Plimpton, S. Fast parallel algorithms for short-range molecular dynamics. J. Comput. Phys. 117, 1–19 (1995).

173. Rosa, A. & Everaers, R. Structure and dynamics of interphase chromosomes. PLoS Comput. Biol. 4, e1000153 (2008).

174. Conte, M. et al. Loop-extrusion and polymer phase-separation can co-exist at the single-molecule level to shape chromatin folding. Nat. Commun. 13, 4070 (2022).

175. De Gennes, P. G. Scaling concepts in polymer physics. Cornell University Press, Ithaca N.Y. (1979).

176. Dobin, A. et al. STAR: ultrafast universal RNA-seq aligner. Bioinformatics 29, 15–21 (2013).

177. Liao, Y., Smyth, G. K. & Shi, W. featureCounts: an efficient general purpose program for assigning sequence reads to genomic features. Bioinformatics 30, 923–930 (2014).

178. Love, M. I., Huber, W. & Anders, S. Moderated estimation of fold change and dispersion for RNA-seq data with DESeq2. Genome Biol. 15, 550 (2014).

179. Hao, Y. et al. Dictionary learning for integrative, multimodal and scalable single-cell analysis. Nat. Biotechnol. (2023) doi:10.1038/s41587-023-01767-y.

180. Bonev, B. et al. Multiscale 3D Genome Rewiring during Mouse Neural Development. Cell 171, 557–572.e24 (2017).

181. Fang, M., Xiang, F.-L., Braitsch, C. M. & Yutzey, K. E. Epicardium-derived fibroblasts in heart development and disease. J. Mol. Cell. Cardiol. 91, 23–27 (2016).

182. Rodríguez-Carballo, E. et al. The HoxD cluster is a dynamic and resilient TAD boundary controlling the segregation of antagonistic regulatory landscapes. Genes Dev. 31, 2264–2281 (2017).

183. Li, H. et al. Segregation of morphogenetic regulatory function of Shox2 from its cell fate guardian role in sinoatrial node development. Commun Biol 7, 385 (2024).

